# Zero-mode waveguide nanowells for single-molecule detection in living cells

**DOI:** 10.1101/2023.06.26.546504

**Authors:** Sora Yang, Nils Klughammer, Anders Barth, Marvin E. Tanenbaum, Cees Dekker

**Author notes:** **For correspondence:** (MET); (CD). These senior authors contributed equally to this work (Marvin E. Tanenbaum, Cees Dekker). These authors contributed equally to this work (Sora Yang, Nils Klughammer, Anders Barth).

## Abstract

Single-molecule fluorescence imaging experiments generally require sub-nanomolar protein concentrations to isolate single protein molecules, which makes such experiments challenging in live cells due to high intracellular protein concentrations. Here, we show that single-molecule observations can be achieved in live cells through a drastic reduction in the observation volume using overmilled zero-mode waveguides (ZMWs - subwavelength-size holes in a metal film). Overmilling of the ZMW in a palladium film creates a nanowell of tunable size in the glass layer below the aperture, which cells can penetrate. We present a thorough theoretical and experimental characterization of the optical properties of these nanowells over a wide range of ZMW diameters and overmilling depths, showing an excellent signal confinement and a five-fold fluorescence enhancement of fluorescent molecules inside nanowells. ZMW nanowells facilitate live-cell imaging, as cells form stable protrusions into the nanowells. Importantly, the nanowells greatly reduce cytoplasmic background fluorescence, enabling detection of individual membrane-bound fluorophores in the presence of high cytoplasmic expression levels, which could not be achieved with TIRF microscopy. Zero-mode waveguide nanowells thus provide great potential to study individual proteins in living cells.

**Graphical abstract:** 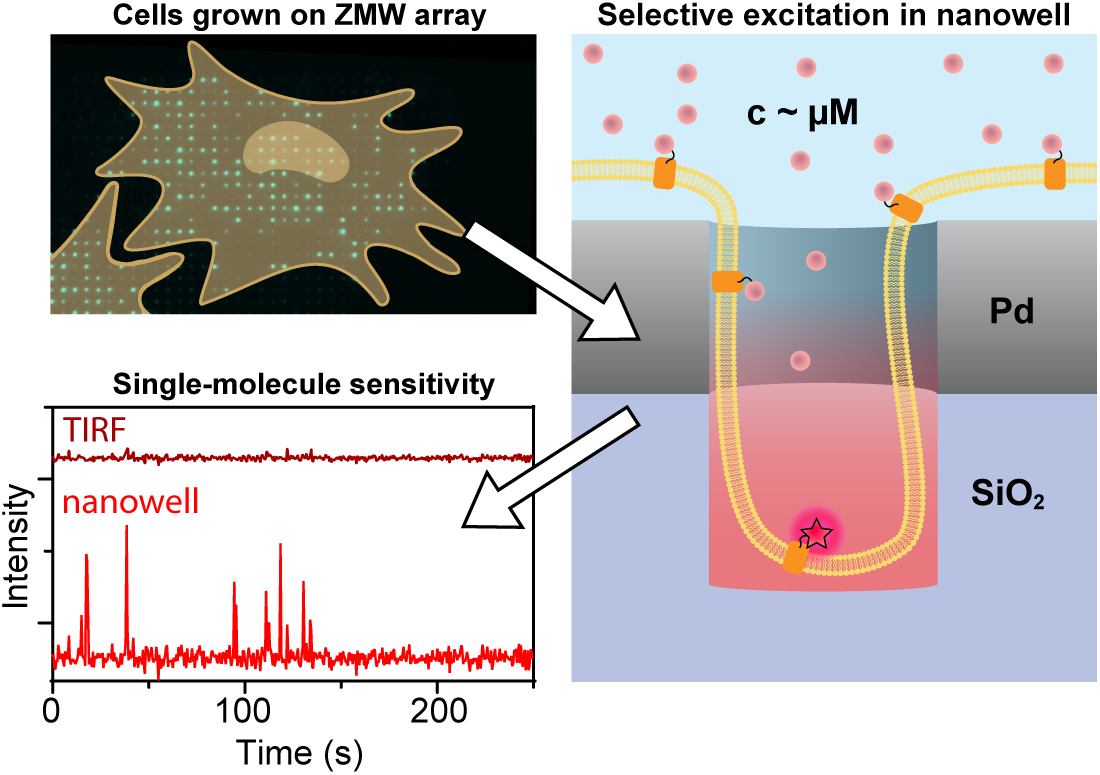

## Introduction

Single-molecule techniques are widely applied to study the behavior of biomolecules or biomolecular complexes, providing mechanistic insights into individual steps of biological processes that would otherwise be averaged out in bulk experiments (***Hinterdorfer and Oijen, 2009***). Imaging-based approaches have been especially powerful in studying single nucleic acid and protein molecules, as they allow tracking of individual biomolecules in space and time. Central to all single-molecule fluorescence imaging techniques is the ability to detect and distinguish a single molecule of interest over the background of fluorescent molecules that are freely diffusing through the solution. The ability to isolate a single molecule by imaging therefore depends on the concentrations of fluorescent molecules and the observation volume; if multiple freely diffusing molecules are present within the observation volume, the isolation of one specific molecule of interest becomes very challenging.

In in vitro experiments, single-molecule observation can easily be achieved by using low concentrations of fluorescent molecules, which limits the number of molecules in the observation volume. However, weak biomolecular interactions (*K*_*d*_ > 1 µM) that require high concentrations cannot be studied at the nano-to picomolar concentrations that are typically employed in in vitro single-molecule experiments. Moreover, studying biomolecules in their natural habitat, the crowded environment of live cells, is also very challenging, as protein concentrations in cells are often in the high nanomolar to micromolar range (***Milo and Phillips, 2015***), which is incompatible with singlemolecule observations. Fundamentally, the concentration limit for single-molecule observation is bounded by the size of the observation volume, which can be minimized using used common optical sectioning methods such as confocal microscopy, total internal reflection microscopy (TIRF), or light-sheet microscopy (***Liu et al., 2015b***). Despite such improvements, the volumes remain on the order of femtoliters, which puts the concentration limit for isolating single molecules at ≈ 1 nM (***Kubitscheck, 2017***).

A much more drastic confinement of the observation volume can be achieved using zero-mode waveguides (ZMW), which are subwavelength apertures in a metal film. Owing to their small size (∼100 nm), ZMWs effectively block the propagation of incident light of wavelengths above a characteristic cutoff wavelength *λ*_*c*_, *λ* > *λ*_*c*_ = 1.7*d*, where *d* is the diameter of the aperture. Within the ZMW, an evanescent field forms which to first order follows an exponential decay as 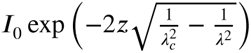 (***Jackson, 1962***). Typical decay lengths are on the scale of several tens to hundred of nanometers, depending on the ZMW diameter, the wavelength of incident light in the surrounding medium, and the ZMW material. Thus, by providing observation volumes in the zeptoliter range, ZMWs enable single-molecule studies at even micromolar concentrations (***Levene et al., 2003***). ZMWs made from gold or aluminum have been extensively studied (***Rigneault et al., 2005***; ***Levene et al., 2003***; ***Gérard et al., 2008***; ***Aouani et al., 2009***; ***Martin et al., 2016***; ***Wu et al., 2019***; ***Al Masud et al., 2020***; ***Patra et al., 2022***) and used for a variety of in vitro single-molecule applications (***Samiee et al., 2005***, ***2006***; ***Auger et al., 2014***; ***Assad et al., 2016***; ***Larkin et al., 2017***; ***Baibakov et al., 2019***; ***Hoyer et al., 2022***), and notably for DNA sequencing (***Rhoads and Au, 2015***). Recently, we have introduced the use of palladium for free-standing ZMWs (***Klughammer and Dekker, 2021***), which were applied to the in vitro study of nuclear transport (***Klughammer et al., 2023***). Palladium offers excellent mechanical and chemical stability, can easily be modified via thiol chemistry (***Love et al., 2003***, ***2005***; ***Klughammer et al., 2023***), and provides reduced photoluminescence in the blue spectral region compared to gold (***Mooradian, 1969***; ***Boyd et al., 1986***; ***Klughammer and Dekker, 2021***). Importantly, Pd is compatible with live cell experiments due to its low cytotoxicity (***Jiang et al., 2004***).

While the vast majority of studies applying ZMWs to single-molecule measurements have been performed in vitro, a few studies have shown that ZMWs made of aluminum can be applied to single-molecule imaging of cellular (membrane) proteins as well, as cells can form protrusions that penetrate into ZMWs (***Wenger et al., 2007-02***; ***Moran-Mirabal et al., 2007***; ***Richards et al., 2012***), which has enabled single-molecule observation of membrane composition (***Wenger et al., 2007-02***) and membrane channels (***Richards et al., 2012***). Inspired by this work, we hypothesized that the creation of nanowells in the glass coverslip below the ZMWs (see ***Figure 1*** A) provide a means of finetuning the size of the observation volume and result in excellent optical properties, while allowing cellular protrusions to enter the nanowells. (***Holzmeister et al., 2014***; ***Gregor et al., 2019***). Moving the observation volume slightly away from the ZMW cavity can potentially lead to an increase of the single-molecule fluorescence signal due to enhancement of the excitation field or modulation of the radiative and non-radiative rates by the metal, as has previously been shown for aluminum (***Miyake et al., 2008***; ***Tanii et al., 2013***; ***Jiao et al., 2014***) and gold (***Wu et al., 2019***). Overmilling could also allow cells to penetrate more deeply through the ZMWs and allow facile imaging not only of membrane bound proteins, but of proteins in the cytoplasm too, greatly expanding the potential applications of ZMW imaging of living cells.

**Figure 1.**
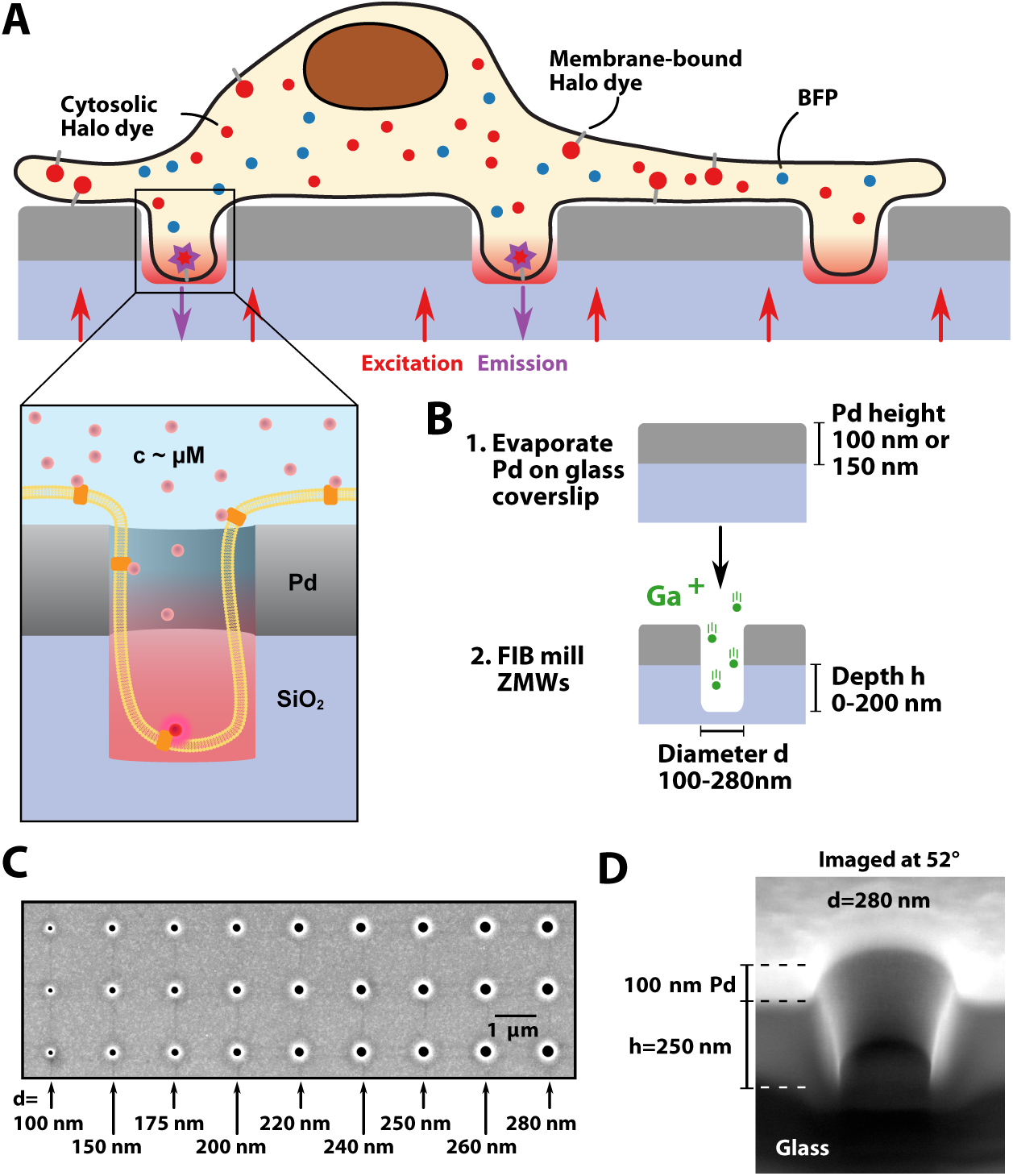
Schematic of the experiment and fabrication of overmilled ZMWs. **A:** Schematic of a cell on top of an array of overmilled ZMWs. Nanowells below Pd ZMWs allow for the observation of single membrane bound fluorophores despite a high abundance of cytoplasmic fluorophores. **B:** Pd is evaporated onto a glass coverslip and ZWMs are created by local focused ion beam milling. Pore diameters used in the study ranged between 100 nm to 280 nm and overmilling depths ranged between 0 nm to 200 nm. **C:** SEM image showing ZMWs with different pore diameters. **D:** The depth of milling was measured by cutting through the pores with a focused ion beam, and measuring the height when imaging under an angle of 52°. **Figure 1—Figure Supplement 1.** Additional SEM images and layouts of version 1 and 2 arrays.

Here, we establish palladium ZMWs nanowells as a tool for single-molecule studies in live cells (***Figure 1*** A). We fabricated ZMW arrays using focused ion beam (FIB) milling, which allowed us to survey a wide range of diameters and overmilling depths to optimize the design for both optical performance and cell compatibility. Finite-difference time-domain (FDTD) simulations of the excitation intensity and fluorescence emission showed an effective reduction of the observation volume to the nanowell below the ZMW and suggested a potential fluorescence enhancement due to the focusing of the excitation intensity within the well, facilitated by the formation of a standing wave below the metal layer. The theoretical results are corroborated by single-molecule experiments on freely diffusing fluorophores which confirmed the signal confinement and showed an up to five fold fluorescence enhancement. Using live-cell imaging, we show that human osteosarcoma U2OS cells readily protruded into the nanowells, protruding more efficiently when ZMWs were overmilled. Cell protrusions remained stable over the timescale of minutes, enabling singlemolecule observation of individual membrane-bound fluorophores even in the presence of high cytoplasmic concentrations of the same fluorophores. This was only possible due to the efficient suppression of cytoplasmic background signal by the ZMW, whereas conventional TIRF microscopy did not allow single molecules to be followed in this setting. Oblique illumination of the nanowells lead to a further reduction of the background levels. Due to their excellent cell compatibility, overmilled Pd ZMWs can be readily applied for single-molecule studies of biological processes in living cells at physiological concentrations.

## Results

### Fabrication of Pd ZMWs on glass

To fabricate nanowells, we first applied a thin (100 nm or 150 nm) palladium layer to standard glass coverslips covered with a 5 nm Ti adhesion layer by physical vapor deposition (***Figure 1*** B). In contrast to previous studies that used aluminum (***Wenger et al., 2007-02***; ***Moran-Mirabal et al., 2007***; ***Richards et al., 2012***), we chose palladium due to its suitability for nanostructuring, good chemical stability, low photoluminescence in the visible spectrum, and low cytotoxicity (***Jiang et al., 2004***; ***Love et al., 2005***; ***Klughammer and Dekker, 2021***). Palladium surfaces can also easily be functionalized using thiols, which provides a strategy for the specific immobilization of molecules and thus can be used for surface passivation via self-assembled monolayers or may be useful for promoting cell adhesion for certain cell types (***Love et al., 2003***). As in our previous studies (***Klughammer and Dekker, 2021***; ***Klughammer et al., 2023***), we used focused ion beam (FIB) milling to create pores in the metal layer, which allows the precise tuning of pore diameters and pore depths within a single array (***Figure 1*** B). We manufactured arrays containing pores of different sizes and depths, including larger marker holes for identification of the different areas within the arrays (***Figure 1***—***Figure Supplement 1***). Typically, 16 arrays were placed on a single glass coverslip, each containing ≈3000 nanowells of varying diameter and depth (***Figure 1*** C, ***Figure 1***—***Figure Supplement 1***). Pore diameters were chosen to range between 100 nm to 280 nm based on a previous study that showed cell protrusion into ZMWs (***Richards et al., 2012***). The depth of the nanowells was varied by overmilling into the glass surface below the palladium layer up to 200 nm. An exemple cross-section is shown in ***Figure 1*** D.

### Simulating the optical properties of palladium ZMW nanowells

To guide the selection of the optimal width and depth of the well below the ZMW, we performed finite-difference time-domain (FDTD) simulations of the excitation electromagnetic field and dipole emission within overmilled ZMWs (***Figure 2*** A). These simulations allow us to assess the spatial distribution of the excitation intensity, the modulation of the fluorescence quantum yield of the fluorophore, and the fraction of signal directed towards the detection side, which together define the detectable signal from within the nanowell as the product of these quantities.

**Figure 2.**
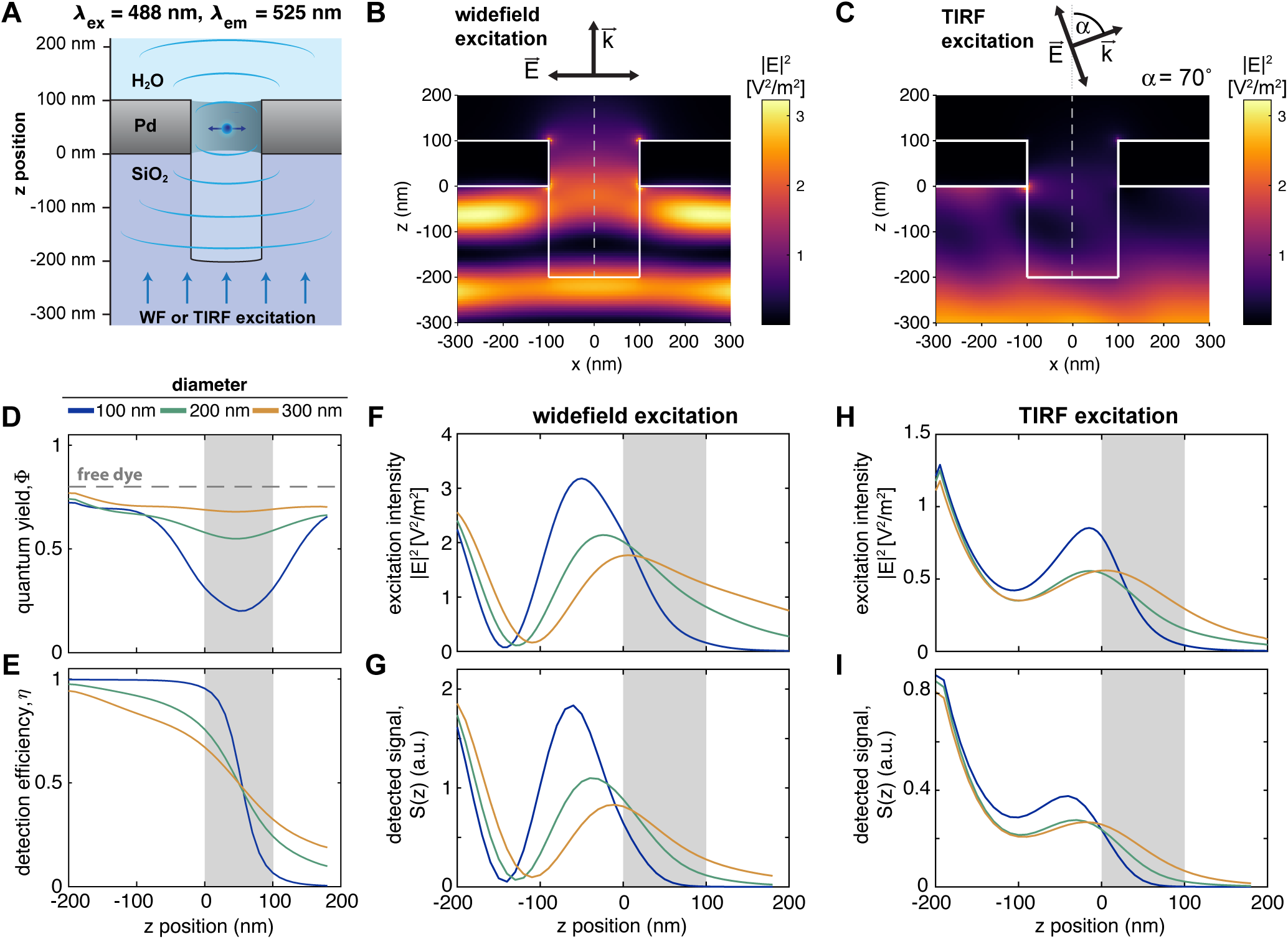
FDTD simulations of the excitation field and fluorescence emission within a nanowell underneath a ZMW. **A:** Schematic of the simulation setup. A dipole was placed at varying depths within the aperture and excited by a plane wave incident from the bottom (widefield, WF) or under an angle of 70° resembling conditions used in TIRF microscopy. **B,C:** Resulting distributions of the excitation field intensity for widefield (B) or TIRF (C) excitation for a ZMW diameter of 200 nm and an overmilling depth of 200 nm. The electric field is polarized along the x-axis. **D,E:** Computed quantum yield and detection efficiency of the dye Alexa488 as a function of the z-position along the central pore axis (dashed line in B,C). **F-I:** Z-profiles of the excitation intensity along the central pore axis (F,H) and the detected signal *S*(*z*) (G,I) under widefield (F,G) and TIRF (H,I) excitation. The position of the metal membrane is indicated as a gray shaded area. **Figure 2—Figure Supplement 1.** FDTD simulations of excitation field intensity distributions under widefield excitation at *λex* = 488 nm. **Figure 2—Figure Supplement 2.** FDTD simulations of excitation field intensity distributions under widefield excitation at *λex* = 640 nm. **Figure 2—Figure Supplement 3.** FDTD simulations of excitation field intensity distributions under TIRF illumination at *λex* = 488 nm. **Figure 2—Figure Supplement 4.** FDTD simulations of excitation field intensity distributions under TIRF illumination at *λex* = 640 nm. **Figure 2—Figure Supplement 5.** FDTD simulations of excitation field intensity distributions under confocal illumination at *λex* = 488 nm. **Figure 2—Figure Supplement 6.** FDTD simulations of excitation field intensity distributions under confocal illumination at *λex* = 640 nm. **Figure 2—Figure Supplement 7.** FDTD simulations of fluorescence emission and detected signal from overmilled ZMWs for Alexa488. **Figure 2—Figure Supplement 8.** FDTD simulations of fluorescence emission and detected signal from overmilled ZMWs for JFX650. **Figure 2—Figure Supplement 9.** Overview of radiative and non-radiative rates obtained from FDTD simulations of fluorescence emission within overmilled ZMWs. **Figure 2—Figure Supplement 10.** FDTD simulations of dipole emission as a function of the distance to the pore walls. **Figure 2—Figure Supplement 11.** Estimation of background signal using FDTD simulations.

To cover the different excitation modes applied in this study, we probed the excitation field distribution at wavelengths of 488 nm and 640 nm upon excitation by a plane wave (widefield), under an angled illumination as used in TIRF microscopy (***Figure 2*** B, C, F, H, and ***Figure 2***—***Figure Supplement 1***–4), as well as upon excitation by a focused beam (***Figure 2***—***Figure Supplement 5***–6). As expected, the zero-mode waveguide effectively blocks the propagation of the excitation light under all conditions for small pore diameters of 100 nm or below, as evident from the profiles of the excitation intensity along the pore axis (***Figure 2*** F, H). At large pore diameters of 200 nm and above, a finite amount of excitation light propagates beyond the ZMW. Due to the reflective surface of the metal, a standing wave is formed on the detection side which leads to an undulation of the excitation field intensity within the overmilled volume (***Figure 2*** B) (***Tanii et al., 2013***; ***Jiao et al., 2014***; ***Wu et al., 2019***). Under TIRF illumination at an angle of 70°, the first maximum of the standing wave pattern is shifted to longer distances from the metal surface compared to widefield excitation because the magnitude of the wave vector orthogonal to the metal surface is reduced (***Figure 2*** C). This results in a reduced excitation intensity within the well, but also provides a more even intensity distribution with an intensity maximum at the bottom of the well. Additionally, the propagation of light through the ZMW is reduced under TIRF illumination compared to widefield excitation, which may limit background cytoplasmic fluorescence in imaging experiments (***Figure 2*** H).

In addition to modulating the excitation field, the metal nanostructure affects the quantum yield of the fluorophore by modulating radiative and non-radiative decay rates, which we assess by simulating dipole emission at varying depths along the central axis of the nanowell (***Figure 2*** A,D and ***Figure 2***—***Figure Supplement 7***–8). Within the ZMW, the radiative rate is reduced while the non-radiative rate is strongly increased due to coupling to the metal nanostructure (grey area in ***Figure 2***—***Figure Supplement 9***). As the distance to the metal increases, the non-radiative losses decrease, while the radiative rate remains relatively constant within the volume beneath the ZMW. Overall, within the proximity of the ZMW, these effects lead to a strong predicted reduction of the quantum yield (***Figure 2*** D) and hence the fluorescence lifetime (***Figure 2***—***Figure Supplement 7*** D, ***Figure Supplement 8*** D), as will be assessed experimentally below. Within the nanowell below the ZMW, the modulation of the decay rates was only weakly dependent on the lateral position (***Figure 2***—***Figure Supplement 10***). Finally, we consider the fraction of the fluorescence emission that can be detected in the experiment, i.e., the signal emitted towards lower side of the ZMW facing the objective lens. Part of the dipole emission from within the ZMW is lost as it propagates towards the upper side of the ZMW that faces away from the objective lens, leading to a sharp decay of the detection efficiency within the ZMW (***Figure 2*** E). Below the ZMW, the effective detection efficiency of the dipole emission is increased approximately two-fold compared to the absence of a metal nanostructure because the metal layer acts as a mirror and propagation of radiation through the ZMW is blocked (***Figure 2*** E and ***Figure 2***—***Figure Supplement 7***,8). Overall, these processes lead to a more effective restriction of the detected signal to the well below the ZMW compared to what is expected from the excitation intensity alone.

The end result is a near-complete suppression of background signals originating from the top side of the ZMW. Under widefield excitation, the simulations predict a background level of 3 % at a pore diameter of 100 nm in a 100 nm thin Pd film, which increases to 10% at 300 nm diameter (numbers are given for overmilling depth of 200 nm, ***Figure 2***—***Figure Supplement 11*** A,D). Under TIRF excitation, the background level decreases further by approximately a factor of two compared to widefield excitation because the excitation intensity is more effectively confined to the nanowell, reaching an excellent signal-to-background ratio of ∼25 even for a large pore diameter of 300 nm at an overmilling depth of 200 nm (***Figure Supplement 11*** E).

In summary, the FDTD simulations show that the observed signal remains effectively confined to the overmilled volume and the ZMW even for pore diameters of up to 300 nm (***Figure 2*** G,I and ***Figure 2***—***Figure Supplement 7***–11). Notably, no enhancement of the fluorescence emission is expected as the presence of the metal waveguide is found to significantly reduce the fluorescence quantum yield (***Figure 2*** D). On the other hand, the excitation field is enhanced in the proximity of the metal surface due to the formation of the standing wave, reaching peak intensities that are up to three-times higher compared to the absence of waveguide (***Figure 2*** B,C,F,H), and the detection efficiency is increased two-fold because the dipole emission is directed towards the detection side (***Figure 2*** E). Together, these effects lead to a significant enhancement of the detected signal from the nanowells.

### Nanowells provide signal confinement and enhancement

To corroborate the theoretical results, we performed measurements on freely diffusing dyes in water in the nanowells using confocal excitation, see ***Figure 3*** A. As expected, the detected fluorescence signal of the dyes Alexa488 and JFX650 in the nanowells increased with both the pore diameter and milling depth (***Figure 3*** B and ***Figure 3***—***Figure Supplement 1*** and C,H,M of ***Figure 3***— ***Figure Supplement 2*** and 3). Time traces of the signal within the wells showed fluctuations originating from the diffusion of fluorophores (***Figure 3*** C). Using fluorescence correlation spectroscopy (FCS), we estimated the number of particles within the nanowells which ranged between 0 and 20 (***Figure 3*** D and ***Figure 3***—***Figure Supplement 2***–3). We observed a linear scaling of the particle number with the milling depth and quadratic scaling with the pore diameter as expected for the cylindrical wells (***Figure 3***—***Figure Supplement 4*** A,B). While the corresponding volumes scaled well with predictions, absolute volumes estimated by FCS exceeded the volume of the nanowells including the ZMW volume by a factor of 2-6 (***Figure 3***—***Figure Supplement 4*** C,D). Comparable deviations had also been observed in previous studies (***Levene et al., 2003***; ***Rigneault et al., 2005***; ***Lenne et al., 2008***; ***Wu et al., 2019***), and were attributed to the signal contribution of many dim fluorophores from the highly concentrated solution that leaks through the ZMW from the other side (***Levene et al., 2003***). This explanation is further supported by the fact that the volume mismatch is largest at high volumes (***Figure 3***—***Figure Supplement 4*** C,D), since large ZMW diameters showed a higher background signal also in FDTD simulations (***Figure 2*** G–I, ***Figure 2***—***Figure Supplement 11***). The residence time of the dyes within the nanowell, as seen from the decay of the FCS curves, likewise increased with the size of the well (***Figure 3***—***Figure Supplement 1***–2).

**Figure 3.**
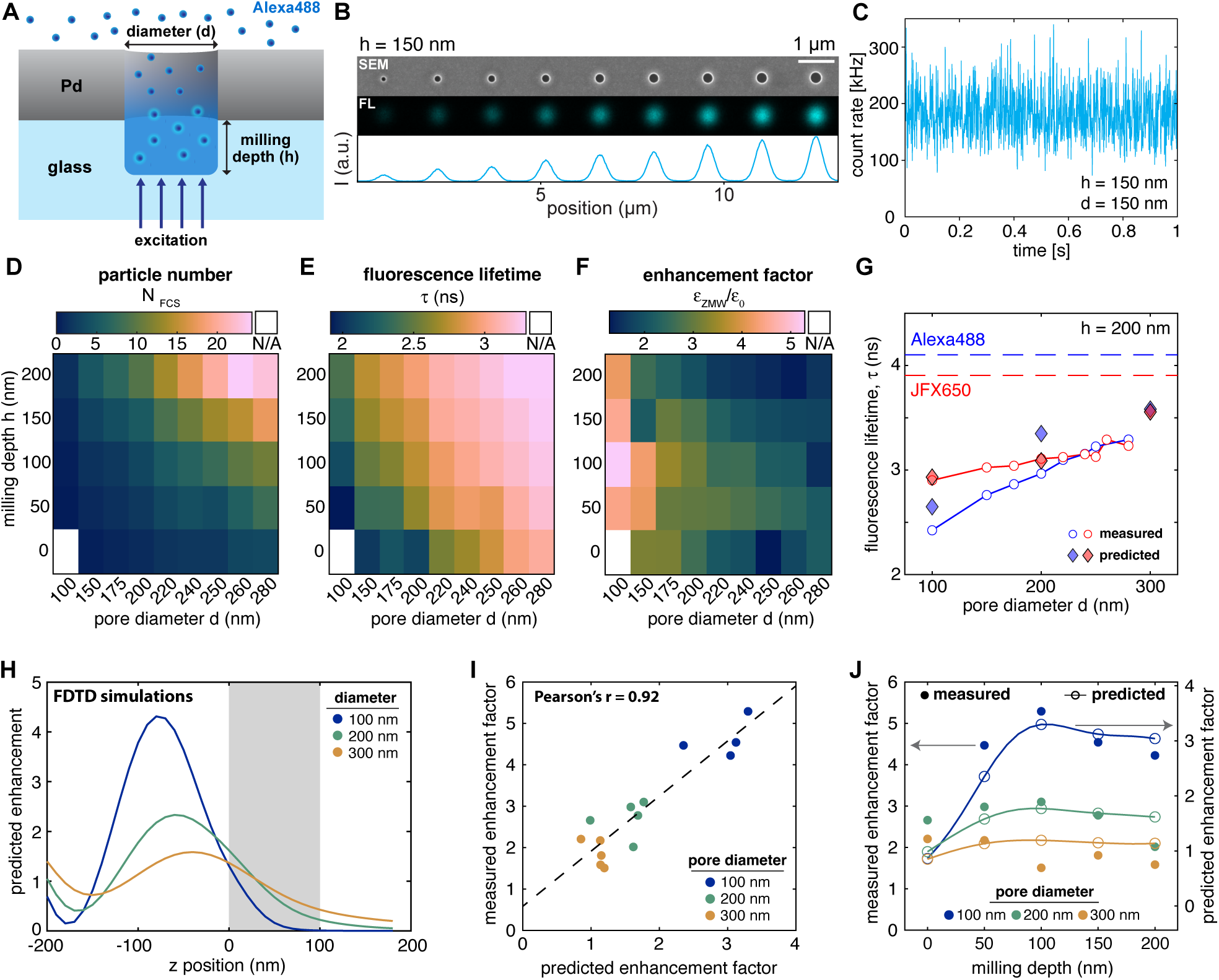
Experimental characterization of fluorescence properties in ZMWs. **A:** Schematic of a ZMW with freely diffusing Alexa488 dye. **B:** SEM (top) and confocal fluorescence (middle) image of a pore array with a milling depth of 150 nm. The fluorescence image was acquired at 1 µM concentration of Alexa488. The intensity profile of the fluorescence image is shown below. The scalebar corresponds to 1 µm. **C:** Example fluorescence time trace (binning: 1 ms) acquired at a concentration of 500 nM Alexa488 for a ZMW with a diameter of 280 nm and no overmilling (h = 0 nm). **D-F:** Heatmaps of the average number of particles in the observation volume *NFCS*, fluorescence lifetime *τ*, and signal enhancement factor defined as the ratio of the counts per molecule in the ZMW compared to free diffusion, *ε*_ZMW_∕*ε*_0_ acquired for 500 nM of Alexa488. Data marked as N/A could not be quantified due to insufficient signal. **G:** Comparison of measured and predicted fluorescence lifetimes from FDTD simulations for an overmilling depth of 200 nm. The lifetimes of the free dyes are shown as dashed lines. **H:** Predicted signal enhancement compared to a free-diffusion experiment as a function of the z position obtained from FDTD simulations (see ***Figure Supplement 7*** for details). **I:** Linear regression of the measured versus the predicted signal enhancement. **J:** Comparison of measured and predicted enhancement factors as a function of the overmilling depth. **Figure 3—Figure Supplement 1.** Experimental characterization of photophysics and diffusion within ZMWs. **Figure 3—Figure Supplement 2.** Extracted parameters for the dye Alexa488 in overmilled Pd ZMWs. **Figure 3—Figure Supplement 3.** Extracted parameters for the dye JFX650 in overmilled Pd ZMWs. **Figure 3—Figure Supplement 4.** Quantification of the observation volume in overmilled ZMWs. **Figure 3—Figure Supplement 5.** Comparison of extracted parameters for the dyes Alexa488 and JFX650 in ZMW. **Figure 3—Figure Supplement 6.** Correlation between fluorescence lifetime and molecular brightness in ZMW for Alexa488 and JFX650. **Figure 3—Figure Supplement 7.** FDTD simulations of excitation field and fluorescence emission in the absence of a Pd layer. **Figure 3—Figure Supplement 8.** Comparison of experimental and predicted enhancement factors in overmilled Pd ZMWs.

To gauge the amount of radiative and non-radiative rate enhancement experienced by the dyes, we quantified the excited state fluorescence lifetime (***Figure 3*** E), where a reduced lifetime indicates a stronger enhancement of either radiative or non-radiative relaxation. Fluorescence decays were well described by a mono-exponential model function (***Figure 3***—***Figure Supplement 1***). We observed the strongest modulation of the fluorescence lifetime for small pore diameters and shallow wells where the dye is restricted within the proximity of the metal aperture. The predicted signal-averaged fluorescence lifetimes from the FDTD simulations showed excellent quantitative agreement with the experimental values (***Figure 3*** G).

To test for a potential enhancement of the signal emanating from the nanowells, we define a signal enhancement factor by comparing the molecular brightness of the fluorophore (as measured from the FCS analysis) within the nanowell to the free-diffusion value. For both dyes, a signal enhancement of 2-5 was observed across the entire parameter space (***Figure 3*** F, ***Figure 3***—***Figure Supplement 5***). The largest enhancement was observed at small pore diameters, reaching a maximum value of 4-5 at a pore diameter of 100 nm and a milling depth of 100 nm for Alexa488. Higher molecular brightness correlated with a reduced fluorescence lifetime (***Figure 3***—***Figure Supplement 6***). Notably, the highest signal enhancement was not obtained at zero overmilling where the dyes are confined to the ZMW, but rather increased as the nanowell extended into the glass up to a depth of 100 nm, after which the signal enhancement was reduced.

To understand the observed enhancement, we computed the theoretical enhancement factor from the FDTD simulations (***Figure 3*** H, ***Figure 3***–***Figure Supplement 7***). This enhancement originates predominantly from a focusing of the excitation light due to the formation of the standing wave, as no quantum yield enhancement was found to be present due to losses to the metal nanostructure (***Figure 2***–***Figure Supplement 9***). Accordingly, the z-profile of the enhancement factor is dominated by the profile of the excitation field. Experimental and predicted enhancement factors showed excellent correlation, although the experimental enhancement factors slightly exceeded the predicted values (***Figure 3*** I). The predicted enhancement factors reproduced well the experimental trends obtained for different pore diameters and milling depths, confirming the maximum of the enhancement at a depth of approximately 100 nm (***Figure 3*** J).

In summary, overmilled palladium ZMWs provide an excellent confinement of the detected signal to the volume in a nanowell below the metal aperture for pore diameters up to 300 nm, while simultaneously offering up to a five-fold signal enhancement by focusing the excitation power within the nanowell.

### Membrane protrusions of live cells can be imaged through Pd ZMWs

To investigate the extent of cell membrane protrusion into ZMWs, cell membranes were fluorescently labeled using a blue fluorescent protein (BFP) fused to the transmembrane domain of the transmembrane protein CD40 (CD40TM-BFP). Cells expressing CD40TM-BFP were grown on a Pd film containing arrays of ZMWs with different diameters and depths (***Figure 1***—***Figure Supplement 1***). Based on their morphology, cells appeared healthy on Pd surfaces for at least two days. When imaged through the ZMWs, we observed BFP signal in many nanowells (***Figure 4*** A). To verify the presence of cells on the ZMW arrays, coverslips were flipped upside-down to image the entire cells using widefield fluorescence microscopy (***Figure 4***—***Figure Supplement 1***). The location of BFP-positive nanopores corresponded well to the position of cells on the nanopore array, indicating that the observed fluorescence in nanopores originated from cells that had membrane protrusions in the pores. To examine the relationship between pore size and cell membrane protrusion into pores, the BFP fluorescence intensities inside nanowells were analyzed for different pore sizes (***Figure 4*** B and ***Figure 4***—***Figure Supplement 1*** C). The amount of protrusions into nanowells, as assessed from the BFP signal, decreased both with the decreased pore diameter and milling depth, with the smallest pore diameter (*d* = 100 nm) only showing a very small amount (few percent) of occupied pores (***Figure 4*** C). Efficient cell protrusions into nanowells with an occupancy of up to 50 % occured for pore diameter above 200 nm and milling depths above 50 nm. Importantly, the low occupancy for pores with no overmilling confirmed that the creation of nanowells with overmilling is key to observation of the cell protrusions. Increased area of the glass surface in overmilled nanowells may facilitates the cell adhesion within the nanowells, leading to improved protrusion in nanowells compared to ZMWs without overmilling

Having established that cells protrude into the nanowells, we assessed the stability and dynamics of the signals from the cell protrusions. For this, we generated a monoclonal cell line that expressed cytoplasmic BFP to assess whether cytoplasmic proteins could be imaged on experimentally relevant timescales within the nanowells (***Figure 5*** A). Based on the efficiency of cell protrusion into nanopores of different sizes (***Figure 4*** C), we narrowed the range of pore diameters down to a range of 170 nm to 250 nm (***Figure 1***—***Figure Supplement 1***). We observed BFP cytoplasmic fluorescence of varying intensity in a large proportion of the pores (***Figure 5*** A and ***Figure 5***—***Figure Supplement 1***). To assess the stability of the cell protrusions in nanowells, we monitored the BFP signal within individual pores over time (***Figure 5*** C,D). For approximately 50 % of pores with BFP signal, the intensity remained largely constant over the acquisition time of 5 minutes. However for a subset of pores, large fluctuations of the BFP intensity were observed, suggesting movement of cell protrusions in and out of the observation volume (***Figure 5***—***Figure Supplement 2***). Shallow pores (ℎ = 50 nm) exhibited a stable signal more frequently compared to deeper pores (***Figure 5***— ***Figure Supplement 2*** C), suggesting that pore depth influenced the stability of cell protrusions in the nanowell. When analyzing BFP intensities among different pores, we found a clear bimodal distribution for all pore sizes, with BFP-positive pores either having a high or low BFP signal (with <10 % of the signal obtained for pores with high intensity, ***Figure 5*** B,E). We hypothesized that the high BFP intensities originated from pores containing well-defined cell protrusions, while low in-tensities reflected pores that were covered by a cell, but in which cells did not insert a protrusion (see cartoons in ***Figure 5*** B). The signal from pores with low BFP intensities then would reflect the fluorescence signal originating from above the ZMW. In our simulations, we found that oblique angle illumination (i.e., TIRF) reduced light penetration through the ZMWs and could thus, in theory, reduce this cytoplasmic signal leaking through the ZMWs (***Figure 2*** F-I and ***Figure 2***—***Figure Supplement 11***).To test this, we measured BFP intensities under both widefield and TIRF illumination. The BFP intensity for pores with low signal was indeed markedly reduced when pores were imaged under TIRF illumination compared to widefield illumination (***Figure 5*** H and ***Figure 5***—***Figure Supplement 3***). The average BFP intensity of pores showing a stable high-BFP signal positively correlated with pore size (***Figure 5*** F), again indicating that the BFP intensity represents the cytoplasmic volume inside the pore. The results support the hypothesis that pores with low BFP signal were covered by cells but their membrane did not penetrate into the pores, while the nanowell was occupied by a cell protrusion in the case of a high BFP signal.

**Figure 4.**
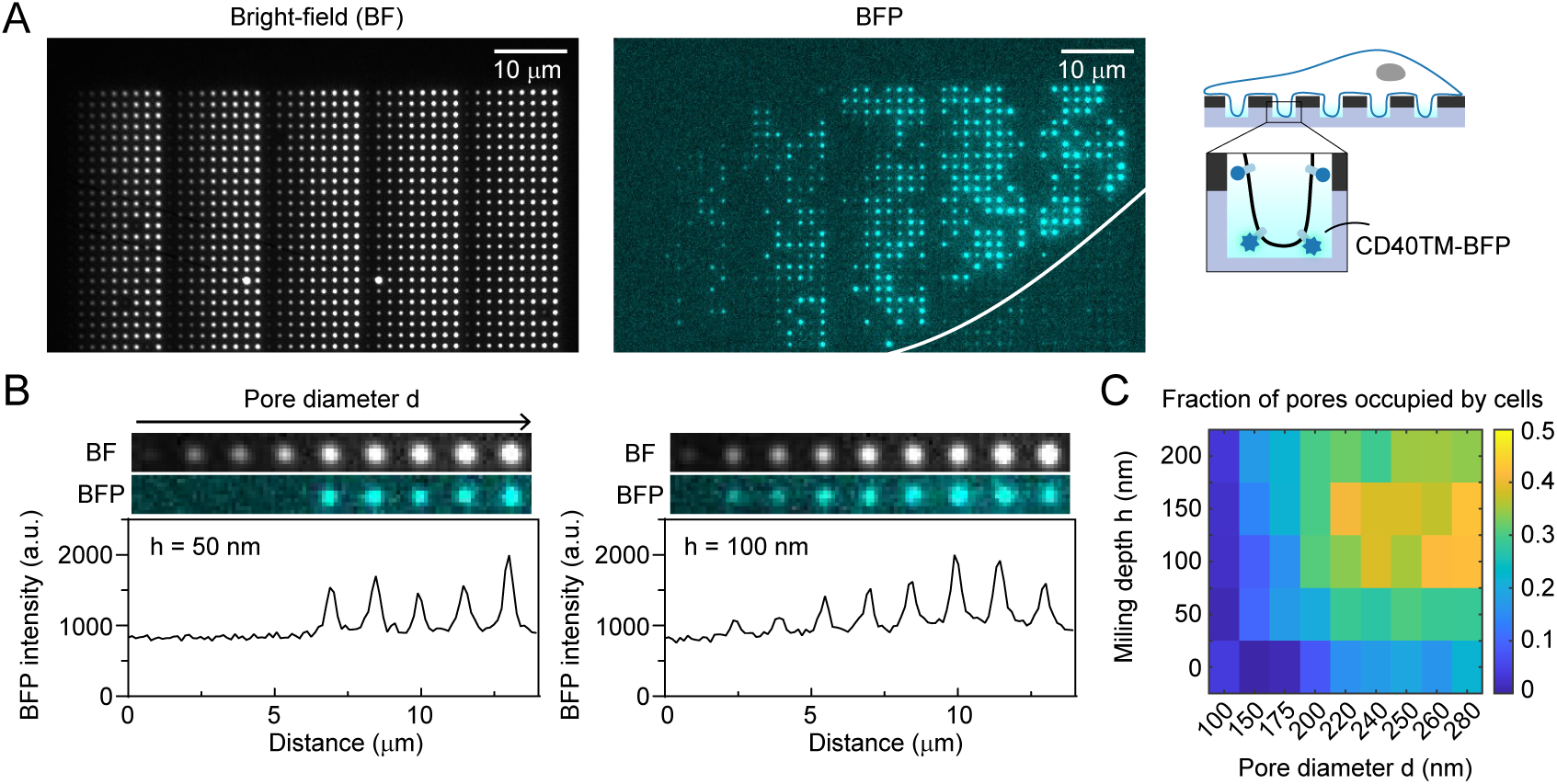
The cell membrane protrudes into nanowells. **A:** U2OS cells expressing CD40TM-BFP on a ZMW array of version 1. Bright-field (left) and BFP fluorescence (right) images are shown. White outline represents an estimated outline of a cell on the surface. **B:** Intensity profiles of BFP fluorescence along different pore diameters (100 nm to 280 nm) for a milling depth of 50 nm (left) and 100 nm (right). **C:** Fraction of pores with detectable BFP signal for each pore size, as determined from 7367 pores potentially covered by cells. **Figure 4—Figure Supplement 1.** Images from flipped conformation confirming cell’s presence on the ZMW array.

**Figure 5.**
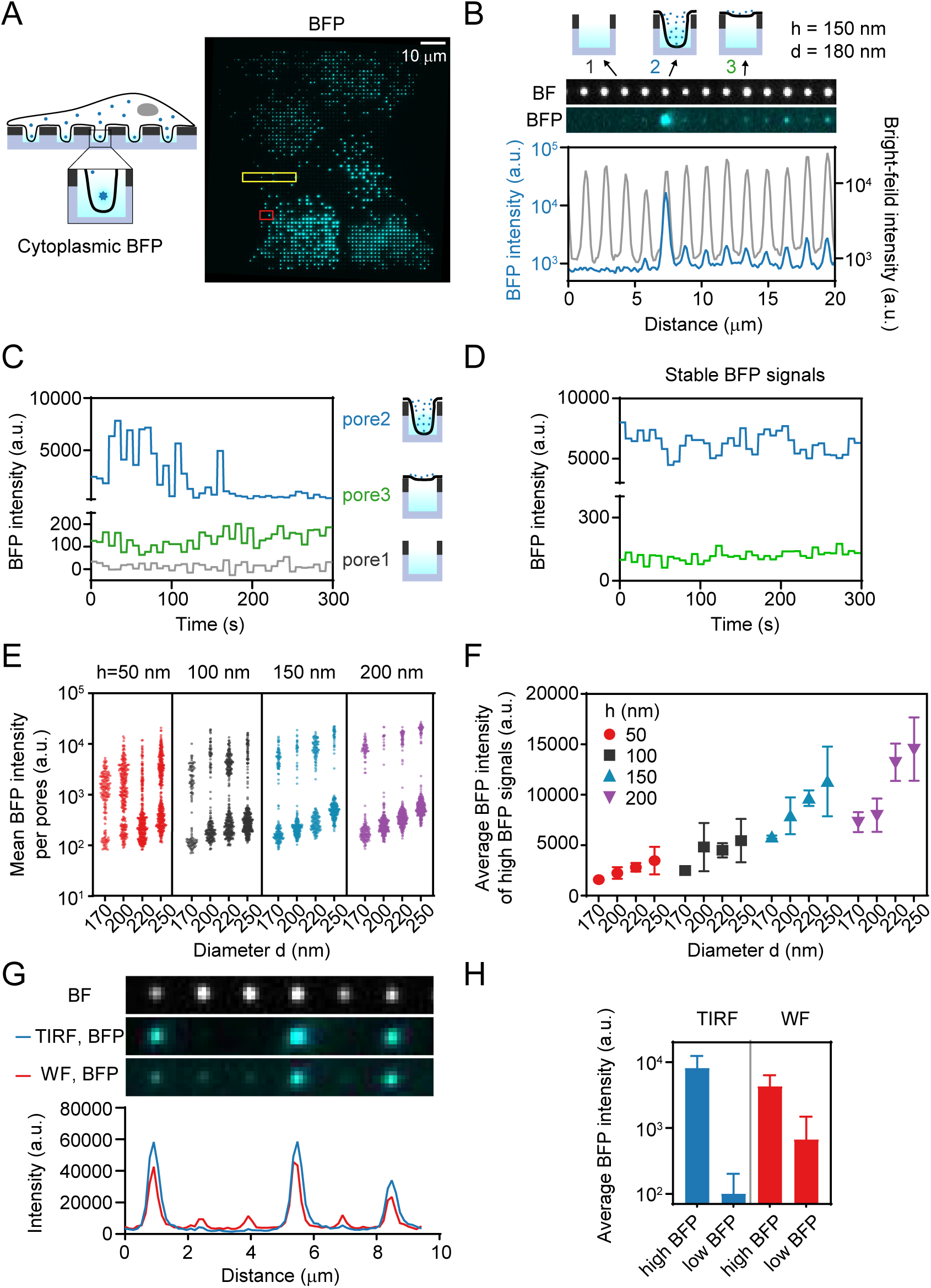
Imaging of single fluorophores in live cells that protrude into nanowells. **A:** U2OS cells expressing cytoplasmic BFP were grown on the ZMW arrays of version 2 and imaged using transmission light (left) or BFP fluorescence (right). Scale bar: 10 µm. **B:** Intensity profiles of BFP and transmission light for pores in the yellow box in (A). **C:** BFP fluorescence time traces of the pores indicated in B. **D:** Representative time traces of BFP intensity for pores with stable high (blue line) or stable low (green line) BFP signal intensity. The pores correspond to the pores denoted by the red box in (A). Images were acquired every 7.5 s. **E:** Each dot represents the average BFP intensity of a time trace for an individual pore showing stable signal. There are two distinct populations for each pore size. The number of measurements per pore size ranges from 113 to 413. **F:** Average BFP intensity (Mean ± sd from 3 independent experiments) of the high BFP signals in (E). **G:** BFP intensity profiles under widefield and TIRF illumination. Under TIRF illumination, peak intensities are increased while background levels are reduced. **H:** Average BFP intensities of high and low intensity pores under TIRF or WF illumination. Error bars represent the standard deviation. **Figure 5—Figure Supplement 1.** Brightfield and BFP-fluorescence images of array version 2 **Figure 5—Figure Supplement 2.** Representative examples of BFP fluorescence intensity time traces for individual pores. **Figure 5—Figure Supplement 3.** TIRF illumination vs. widefield illumination

### Cytoplasmic background can be efficiently suppressed using Pd ZMWs

Having confirmed that cell protrusions remained stable within nanopores over a timescale of minutes, we next tested whether single protein molecules could be visualized within the ZMWs. As a model, we used the transferrin receptor (TfR), a transmembrane protein involved in iron delivery into the cells. To achieve specific labeling TfR, we fused it to the HaloTag, a small protein tag that can be covalently labeled with fluorescent dyes (***Los et al., 2008***). We generated a cell line, stably expressing HaloTag-TfR as well as cytoplasmic BFP. BFP was used as a marker to identify which pores where occupied by cell protrusions (***Figure 6*** A). For single-molecule imaging, we chose the JFX650 dye as the fluorophore for labeling HaloTag-TfR (JFX650-HaloTag ligand) due to its high brightness and photostability (***Grimm et al., 2021***). Conventional TIRF microscopy on glass coverslips confirmed the correct localization of the HaloTag-TfR in the plasma membrane and identified the optimal dye concentration to visualize single-HaloTag-TfR molecules (***Figure 6*** B). Cells were then cultured on ZMWs and imaged using TIRF illumination. HaloTag-TfR signal was monitored from pores with stable cell protrusion as evidenced by constant high cytoplasmic BFP signal over the duration of the experiment (***Figure 6*** C,D). Only pores with high BFP signal intensities exhibited HaloTag-TfR signal (***Figure 6***—***Figure Supplement 1***), further supporting that pores with low BFP signal originated from cells lying on top of the pore without protruding into the pore.

**Figure 6.**
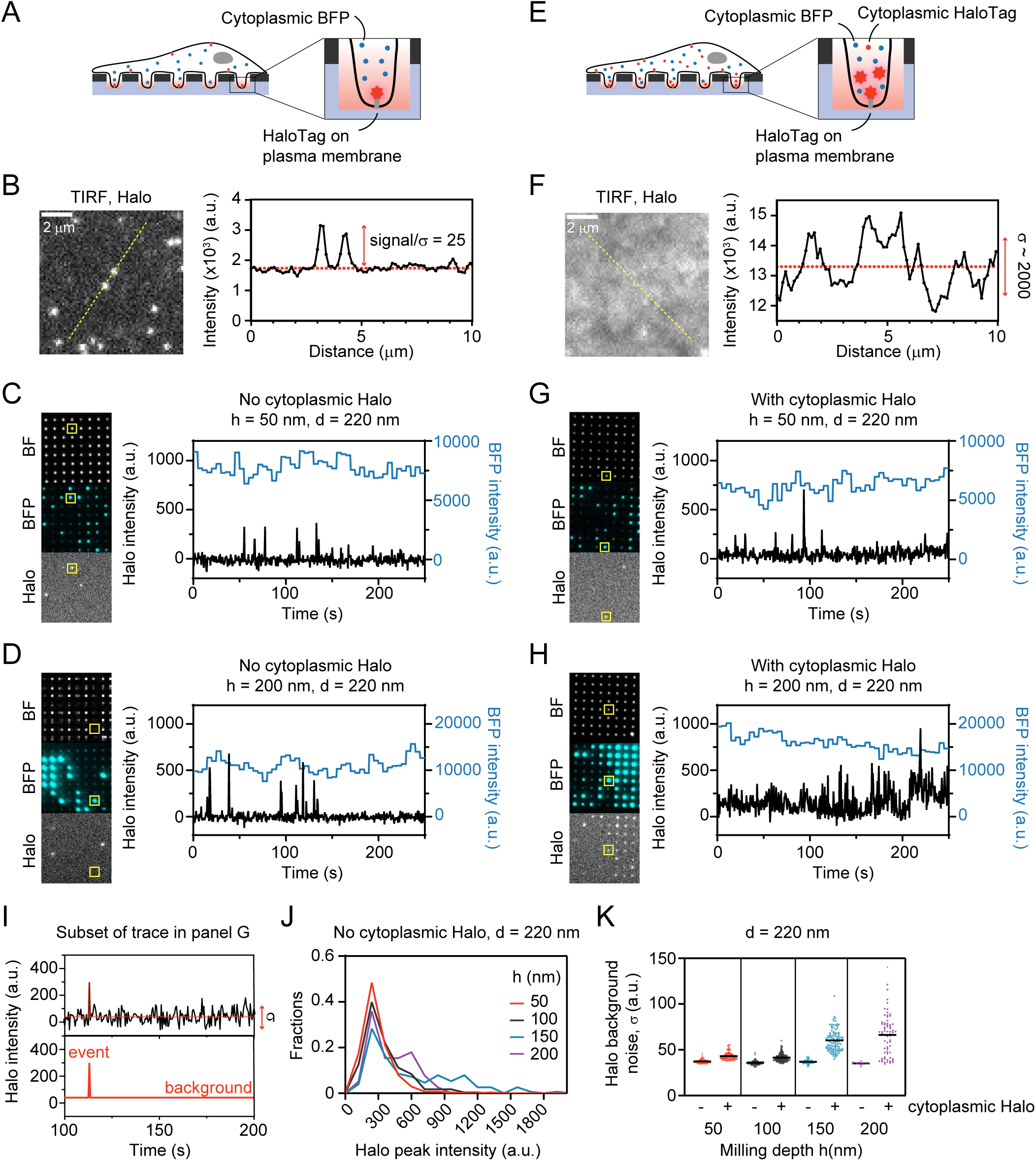
Suppression of cytoplasmic background signal using Pd ZMWs. **A:** Schematic of U2OS cells expressing cytoplasmic BFP and HaloTag-TfR localized in the plasma membrane. **B:** Representative TIRF image of JFX650-labeled HaloTag-TfR. Graph on the right represents an intensity profile along the yellow line. **C,D:** Representative images of bright-field (BF), BFP, and JFX650-HaloTag acquired through nanopores. Fluorescence time trace of a single pore, representing the yellow boxed area in the images. Pore diameter *d* = 220 nm, milling depth *d* = 50 nm (C) and *d* = 200 nm (D). Time interval, 5 s for BFP, 500 ms for JFX650-Halo. **E:** Expression of cytoplasmic HaloTag in the same cell line as in (A). **F:** Representative TIRF image of JFX650-labeled HaloTag-TfR. Graph on the right represents an intensity profile through the yellow line. Scale bar 2 µm. **G,H:** Representative images of BF, BFP, and JFX650-HaloTag acquired under the same imaging condition as C,D, but from the cell line additionally expressing cytoplasmic HaloTag. Fluorescence time trace of a single pore representing the yellow boxed area in the images. Pore diameter *d* = 220 nm, milling depth ℎ = 50 nm (G) and ℎ = 200 nm (H). Time interval, 5 s for BFP, 500 ms for JFX650-Halo. **I:** Analysis of the Halo intensity time trace using a hidden Markov model to determine the Halo peak intensity and background noise of each trace. The background noise is defined as the standard deviation (*σ*) of the background intensity. **J:** Distribution of Halo peak intensities for pores with different milling depths and a constant pore diameter of 220 nm, obtained from the cell line without cytoplasmic HaloTag described in (A). **K:** Background noise *σ* in the red HaloTag channel from individual pores. Each dot represents a single pore. The mean is indicated by black bars. Number of pores: *n* = 86, 133, 116, 177, 72, 113, 14, 69 (in the same order as the graph). **Figure 6—Figure Supplement 1.** BFP and JFX650-HaloTag signal of pores showing low BFP signal.

Next, we probed whether the reduction of the observation volume provided by the ZMWs would allow the detection of single membrane-bound molecules in the presence of a high cytoplasmic background in the same detection channel. To this end, we generated a cell line that expresses high levels of freely-diffusing cytoplasmic HaloTag in addition to membrane-localized HaloTag-TfR (***Figure 6*** E). TIRF microscopy is the gold standard for reduction of fluorescence background from high levels of cytoplasmic proteins. However, at high cytoplasmic Halo expression, TIRF microscopy no longer yielded sufficient background fluorescence reduction to observe single HaloTag-TfR proteins on the membrane, as can be seen in ***Figure 6*** F. In contrast, when cells were cultured on ZMWs with a 50 nm milling depth, single HaloTag-TfR molecules could readily be observed, with a signal-to-noise comparable to when no cytoplasmic HaloTag was present (***Figure 6*** C,G). This shows that the ZMWs effectively suppressed the background signal from the cytoplasmic HaloTag and allowed observation of membrane-localized HaloTag-TfR molecules. In contrast, ZMWs with a 200 nm milling depth, which yield larger optical volumes due to increased nanowell sizes, resulted in significantly higher HaloTag fluorescence background, rendering it impossible to distinguish single HaloTag-TfR molecules from cytoplasmic background signal (***Figure 3*** H).

To quantify the signal-to-noise ratio of single HaloTag-TfR proteins, we applied a hidden Markov model (HMM) analysis to detect the signal spikes representing single HaloTag-TfR molecules diffusing in and out of nanowells (***Figure 6*** I). In the cell line without cytoplasmic HaloTag, no significant differences in the Halo peak intensity and background noise were observed across different milling depths (***Figure 6*** J,K). However, in the cell line expressing high cytoplasmic HaloTag, the background signal significantly increased for deeper pores, due to the larger cytoplasmic volume present within the observation volume (***Figure 3*** K). These results show that the milling depth plays a crucial role for background suppression. Taken together, this work reveals that imaging cells on overmilled ZMWs allows for the visualization of single fluorescent molecules even in the presence of high fluorescent background in the cytoplasm.

## Discussion

Here, we have introduced overmilled palladium ZMWs combined with TIRF illumination as a platform for single-molecule studies in live cells. By creating an attoliter-volume size-tunable nanowell below the ZMW, we achieved a highly confined observation volume that is efficiently penetrated by cell protrusions. The resulting reduction of the cytoplasmic volume combined with favorable optical properties of the nanowell enabled the observation of single fluorescently-labeled cellular membrane proteins even in the presence of high cytoplasmic concentrations of the same fluorophore.

We demonstrated an effective confinement of the observation volume to the nanowell, a strong rejection of background fluorescence originating from the other side of the ZMW and an up to 5-fold signal enhancement, both theoretically using FDTD simulations and by in vitro experiments of freely diffusing fluorophores (***Figure 2*** and ***Figure 3***). Significant non-radiative losses to the metal occurred for shallow milling depths and small pore diameters where the dyes are confined to the proximity of the metal, as evidenced by a strong decrease of the excited state lifetime of the fluorophore, reaching up to a 2.3-fold reduction. Similar changes of the fluorescence lifetime were reported for aluminum (***Rigneault et al., 2005***; ***Jiao et al., 2014***) and aluminum/gold alloy ZMWs (***Al Masud et al., 2020***), with a 2 to 6-fold reduction of the fluorescence lifetime for various fluorophores and ZMW diameters. Due to a lack of radiative enhancement, this results in a strong quantum yield reduction of up to 4-fold within ZMWs with small pore diameters, which however approaches the free dye value quickly with increasing distance from the metal (***Figure 2*** B).

Despite a ≈10 % reduction of the quantum yield throughout the observation volume, the signal from within the nanowells is enhanced by a factor of 2 to 5 due to the combination of two effects. First, the excitation intensity is focused within the nanowell due to the formation of a standing wave below the metal layer that leads to a 3-fold increase of the excitation intensity at its maximum (***Figure 2*** F, ***Tanii et al.*** (***2013***); ***Jiao et al.*** (***2014***)). Second, an approximate 2-fold increase of the detection efficiency arises because the emission is guided towards the detection side due to the reflective metal surface (***Figure 2*** E). The conclusions reached here for the green detection channel using the dye Alexa488 (*λ*_ex_ = 488 nm, *λ*_em_ = 525 nm) apply also to the far-red detection channel using the dye JFX650 (*λ*_ex_ = 640 nm, *λ*_em_ = 670 nm), both from the theoretical (***Figure 2***—***Figure Supplement 7***,8) and experimental side (***Figure 3***—***Figure Supplement 5***), and are thus expected to remain valid over the whole visible spectrum. While similar results are expected also for other metals such as gold and aluminum (***Rigneault et al., 2005***; ***Miyake et al., 2008***; ***Tanii et al., 2013***; ***Jiao et al., 2014***; ***Wu et al., 2019***; ***Gérard et al., 2008***; ***Lenne et al., 2008***; ***Al Masud et al., 2020***; ***Aouani et al., 2009***), palladium offers a simpler fabrication process compared to aluminum by eliminating the need for a SiO_2_ passivation layer, and a reduced photoluminescence in the green spectral range compared to gold (***Klughammer and Dekker, 2021***).

Our insights into the optical properties of the nanowells have a number of consequences for in vitro and in cellulo applications of ZMWs. For standard ZMWs without overmilling, the zeptolitersize observation volume results in very short dwell times of freely diffusing molecules in the observation volume, necessitating immobilization of the molecules of interest. Using overmilled ZMWs, we achieve dwell times for freely diffusing molecules that are comparable to residence times in a diffraction-limited confocal volume, enabling potential applications of overmilled ZMWs in singlemolecule spectroscopy and single-molecule FRET experiments (***Agam et al., 2023***; ***Lerner et al., 2021***). The larger volume of the overmilled ZMWs also provides the possibility to study large and flexible molecules, such as extended DNA/RNA molecules, that are otherwise difficult to confine to the small volume. Furthermore, the distance-dependent quenching by the metal can lead to a significant signal reduction in standard ZMWs (***Holzmeister et al., 2014***), which is avoided in overmilled ZMWs where molecules are at sufficient distance from the metal. The excitation intensity within standard ZMWs also generally remains limited when molecules are not directly immobilized on the glass surface due to the exponentially decaying evanescent field. This situation is resolved in overmilled ZMWs where the excitation power is effectively focused to the nanowell due to the formation of a standing wave, an effect that cannot be exploited in standard ZMWs. While an increase of the excitation intensity can also be achieved by increasing laser powers, the distinctive intensity distribution within the nanowell further improves the background suppression by preferential excitation of molecules below the ZMW. Lastly, the metal layer acts a mirror surface that leads to a more efficient collection of the fluorescence emission from within the nanowell. This provides a more efficient use of the limited signal in single-molecule experiments, which is especially crucial in live cell applications where photostabilization by oxygen scavenging and use of reducing-oxidizing agents is difficult. A drawback of nanowells compared to standard ZMWs is that that larger observation volume in nanowells limits single-molecule detection sensitivity when very high protein concentrations are used (micromolar-millimolar). Therefore, standard ZMWs and nanowells will each have their specific applications, with nanowells performing especially well in cell-based imaging.

Pd-based ZMW nanowells showed excellent compatibility with live-cell imaging. Cells grew read-ily on palladium-coated glass coverslips, adhered to the untreated metal surface, and showed healthy morphologies. Efficient protrusion into the nanowells was observed for most pore sizes, except for the smallest diameter of 100 nm, with the frequency of membrane protrusions increasing both with milling depth and pore diameter, in agreement with previous results on Al ZMWs (***Moran-Mirabal et al., 2007***). Time-dependent fluctuations of the fluorescence signals of cytoplasmic BFP confirmed that cells dynamically explored the nanowells over a timescale of minutes (***Figure 5*** C,D). Approximately 10 % of pores exhibited significant cytoplasmic signal that remained stable over a timescale of 5 min, showing that cells formed stable protrusions into the nanowells on relevant timescales for many biological processes. While the cytoplasmic background signal increased both with pore size (***Figure 5*** E, in agreement with ***Moran-Mirabal et al.*** (***2007***)), it was markedly reduced under TIRF illumination due to a reduced propagation of the excitation light through the ZMW and a more even excitation intensity within the nanowell (***Figure 5*** H, ***Figure 2*** F-I, ***Figure 2***—***Figure Supplement 11***).

We showed the applicability of the nanowells for single-molecule fluorescence experiments in live cells and its superiority compared to TIRF microscopy with respect to single-molecule observations. By following the fluorescence of single membrane-bound fluorophores, we found that signal spikes originating from single molecules were only observed for pores with stable protrusions. Most importantly, single-molecule signals could still be observed under conditions of high cytoplasmic expression, which did not allow for single-molecule experiments using conventional TIRF excitation (***Figure 6*** F-H). Despite the high concentration of fluorophores in the cytoplasm, we could achieve a signal-to-noise ratio of ≈ 7 for shallow pores up to a depth of 100 nm, with noise levels equivalent to when no cytoplasmic signal was present (***Figure 6*** J,K). The results confirm the excellent suppression of the cytoplasmic signal provided by the ZMWs nanowells, allowing monitoring of single fluorophores despite a high cytoplasmic background of the same fluorophore.

We envision that overmilled ZMWs will allow live-cell single-molecule fluorescence experiments in a wider range of cases. More specifically, it provides a benefit in cases where expression levels cannot be controlled, for example when studying endogenously expressed proteins of interest, or when weak interactions are studied requiring high concentrations of the interaction partners. While we have only tested the human osteosarcoma U2OS cell line in this study, live-cell ZMW imaging has been applied successfully in other cell lines, including COS-7 cells (***Wenger et al., 2007-02***), Rat basophilic leukemia (RBL) mast cells (***Moran-Mirabal et al., 2007***), and mouse neuroblastoma N2a cells (***Richards et al., 2012***), suggesting its potential for broad applications. To facilitate cell protrusion into the nanopores, surface coatings or functionalization with specific molecules or peptides can be employed as effective strategies (***García and Boettiger, 1992***; ***VandeVondele et al., 2003***; ***Jiang et al., 2004***), which may extend the application further to a variety of cell types. Since the total cellular membrane fraction that protrudes into the nanowells is small (≤ 1 %), efficient approaches for membrane recruitment and immobilization of low-concentration complexes will be required. A potential strategy could be through introduction of a designed transmembrane protein with an intracellular docking platform and extracellular binding to the glass wells below the Pd layer using silane chemistry (***Malekian et al., 2018***) or electrostatic interactions with the glass surface using poly-lysine (***VandeVondele et al., 2003***; ***Liu et al., 2015a***). We envision many applications of our method for the study of protein-protein and protein-RNA interactions in the cytoplasm, enzymatic activities of single proteins or complexes such as ribosomes (***Yan et al., 2016***) or proteasomes (***Bard et al., 2018***), and protein conformational dynamics by combination with single-molecule FRET (***Agam et al., 2023***). In this study, we focused on membrane-associated proteins, which have a longer residence time within the nanowells compared to cytoplasmic fluorophores. Nonetheless, it is likely that ZMW nanowells can similarly be adopted to study freely diffusing molecules which would increase the applicability of overmilled ZMWs even further.

## Conclusions

We introduced the use of overmilled zero-mode waveguides made of palladium combined with TIRF illumination for live-cell imaging. We performed a thorough theoretical and experimental characterization of the optical properties of the nanowells using FDTD simulations and fluorescence experiments of freely diffusing organic dyes, which together delineated the signal confinement and fluorescence enhancement within the overmilled nanowell volume. Live-cell experiments showed that cells readily protrude into the nanowells, enabling single-molecule fluorescence experiments with excellent signal-to-noise ratio, despite a high cytosolic concentration of the fluorophore. By scanning a wide range of pore diameters and milling depths, we provided comprehensive guidelines for future in vitro or in cellulo single-molecule studies where a compromise must be found between the required background suppression, the desired fluorescence enhancement, and the efficiency of cell protrusions.

## Methods and Materials

### Fabrication of Palladium Zero-Mode Waveguides

Standard borosilicate coverslips (#1.5H, Marienfeld, Germany) were cleaned by consecutive sonication in deionized water, isopropyl alcohol, acetone, and 1 M potassium hydroxide solution, washed with deionized water, and spin dried. A thin adhesion layer of 3 nm titanium was deposited at a rate of 0.05 nm/s under a base pressure of 3 × 10^−6^ torr in a Temescal FC2000 e-gun evaporator. In the same vacuum, a layer of Pd was immediately added on top of the Ti. Two versions of Pd ZMWs were used in this study, which differed by the thickness of the Pd layer. Either 100 nm of Pd was deposited at a rate of 0.1 nm/s (Version 1) or 150 nm at a rate of 0.2 nm/s (Version 2), both under a base pressure below 2 × 10^−6^ torr.

Nanopores were milled through the layers via focused ion beam (FIB) milling on a FEI Helios G4 CX FIB/SEM. To improve consistency, the focus and stigmation of the ion beam was optimized on a graphite standard sample before milling. For the pore arrays of version 1, a 33 pA beam with an acceleration voltage of 30 kV was used. For the pores of version 2, the beam current was set to 430 pA at the same voltage. Due to the higher beam current used for version 2, milling time was reduced to around 4 min per array coming at the cost of less well defined pore diameters (***Figure 1***— ***Figure Supplement 1***). The diameters of the resulting pores were measured using the immersion mode of the scanning electron microscope on the same machine. The depth and opening angle of the pores, resulting from overmilling into the glass surface, was measured by cutting through the pores with the ion beam and imaging under 52° incident angle. The diameters, depths and taper angles of the pores can be found in ***Figure 1*** C,D and ***Figure 1***—***Figure Supplement 1*** C-F.

Prior to experiments, ZMWs were thoroughly cleaned by consecutive sonication in deionized water, ethanol, isopropyl alcohol, acetone, and 1 M potassium hydroxide solution for about 10 min each and exposed to oxygen plasma at a power of 90 W for 1 min. The coverslips could be reused about 10 times, after which the Pd film started to show signs of degradation.

### Single-molecule measurements of free fluorophores

Measurements of freely diffusing fluorophores inside Pd ZMWs were performed on coverslips of version 1 on a Picoquant Microtime 200 microscope operated using the Symphotime software in a temperature controlled room at 21.5 ± 1.0 °C. Lasers were focused by an 60x Olympus UPSLAPO 60XW water immersion objective with a working distance of 280 µm and a numerical aperture of 1.2. Excitation at wavelengths of 640 nm and 485 nm was performed at powers of 10 µW as measured at the sample plane. Pulsed lasers were operated in pulsed interleaved excitation at a repetition frequency of 40 MHz (***Müller et al., 2005***). The molecular brightness of a solution of Alexa488 fluorophores was optimized by adjusting the correction collar of the objective prior to the experiment. The emission light was passed through a 50 µm pinhole, split by a dichroic mirror and was filtered by 525/50 or 600/75 optical band pass filters for the blue and red detection channels, respectively (Chroma, Bellow Falls). Fluorescence emission was detected on single-photon avalanche-diode detectors (PD5CTC and PD1CTC, Micro Photon Devices, Bolzano). Fluorophore solutions of 500 nM of Alexa488 and JFX650-HaloTag were supplemented by 0.1 % Tween20 (Fischer Scientific) to minimize surface adhesion of the fluorophores. Transmission light images of the pore arrays were used to locate the pores prior to fine tuning of the position of the laser focus to maximize the signal.

### FDTD Simulations of light fields inside Pd ZMWs

Three-dimensional FDTD simulations were performed using Lumerical FDTD (ANSYS Inc., USA) as described previously for the characterization of free-standing ZMWs (***Klughammer et al., 2023***) and a description is repeated here for completeness. The surrounding medium was modeled as water with a refractive index of 1.33 and the refractive indices of the 100 nm thick palladium membrane and the SiO_2_ layer were modelled according to (***Alterovitz et al., 1998***). For the simulation of the excitation field, the ZMW was illuminated by a total-field scattered-field source which was polarized in the x-direction. The source was set as a plane wave for widefield excitation and TIRF excitation under an incidence angle of 70°, and a Gaussian source with a numerical aperture of 1.2 for focused excitation. The simulation box size was 1×1×0.8 µm^3^ for widefield and TIRF excitation with a grid resolution of 5 nm. To correctly model the focused beam, a larger box of 4x4x0.8 µm^3^ was required for the Gaussian source, in which case a larger grid resolution of 50 nm was used to model the field further away from the nanowell keeping the 5 nm grid resolution close to the nanowell. The electromagnetic field intensity distributions, computed as the absolute value of the complex electric field, |*E*|^2^|, in the xz- and yz-planes at the center of the nanowell and in the xy-plane at the ZMW entry are s hown in ***Figure 2***—***Figure Supplement 1*** - ***Figure Supplement 6***. To model the flu-orescence emission, a dipole emitter was placed at varying z-positions along the central axis of of the nanowell. The radiated power was monitored on all sides of the simulation box (see below). To compute the detection efficiency, the radiated power was integrated only over the detection side below the palladium layer. For dipole emission, all reported quantities were averaged over horizontal and vertical orientations of the dipole to model isotropic emission. The power was only weakly affected by the lateral position of the emitter with respect to the center of the nanowell (***Figure 2***—***Figure Supplement 10***, ***Levene et al.*** (***2003***)).

### Estimation of quantum yield and fluorescence lifetimes

Quantum yields and fluorescence were computed as described previously (***Klughammer et al., 2023***) and this description is repeated here for completeness. In the absence of the nanostructure, the decay rate of the excited molecule is given by 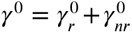, where 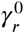 and 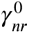 are the radiative and non-radiative decay rates. Here, 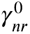 represents the rate of non-radiative relaxation to the ground state due to internal processes, which is assumed to be unaffected by the nanostructure. The intrinsic quantum yield of the fluorophore is defined as 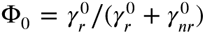 and was obtained from literature as Φ_0_ = 0.8 and 0.53 for Alexa488 and JFX650 (***Hellenkamp et al., 2018-09***; ***Grimm et al., 2021***).

Within the nanowell, the radiative decay rate *γ*_*r*_ is modified. Additionally, a non-radiative loss rate *γ*_loss_ arises due to absorption by the metal nanostructure (***Novotny and Hecht, 2006***). The quantum yield Φ in the presence of the ZMW is given by (***Bharadwaj and Novotny, 2007***):

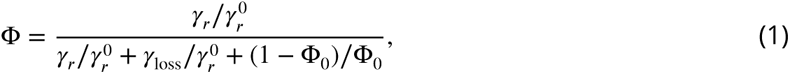

where 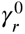 and *γ*_*r*_ are the radiative rates in the absence and the presence of the ZMW respectively. While absolute decay rates *γ*_*r*_, *γ*_loss_, and 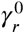 are inaccessible from FDTD simulations, relative rates normalized to the radiative rate in the absence of the ZMW, 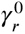, can be obtained from the power *P* radiated by the dipole (***Kaminski et al., 2007***) as:

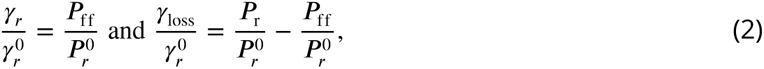

where *P*_*r*_ and 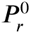 are the powers radiated by the dipole in the presence and absence of the ZMW, and *P*_ff_ is the power that is radiated into the far-field in the presence of the ZMW. See ***Figure 2***—***Figure Supplement 9*** for the obtained z-profiles of the normalized radiative and non-radiative rates.

To obtain the fluorescence lifetime *τ*, which is given by the inverse of the sum of all de-excitation rates, we use the relation *τ* = Φ∕*γ*_*r*_ in combination with eq. 1:

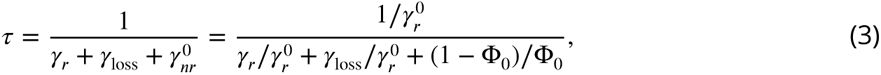

where the intrinsic radiative rate 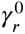 was estimated as 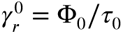 with the experimentally measured fluorescence lifetimes *τ*_0_ for Alexa488 and JFX650 of 4.0 ns and 3.9 ns. The detection efficiency *η* was estimated as the fraction of the power radiated towards the lower (detection) side of the ZMW, 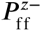, with respect to the total radiated power:

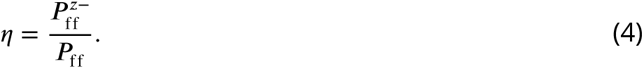

Finally, the total detected signal as a function of the z-position of the emitter within the nanowell was computed as the product of the excitation intensity *I*_ex_(*z*), detection efficiency *η*(*z*), and quantum yield Φ(*z*) as:

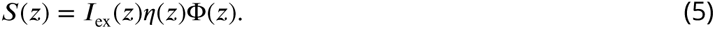

The computed detection efficiency *η*, quantum yield Φ, detected signal *S*(*z*) and lifetime *τ* as a function of the z-position within the ZMW are shown in ***Figure 2***—***Figure Supplement 7***,8.

### Estimation of signal enhancement factors

To estimate the theoretical signal enhancement factor, we performed simulations in the absence of the palladium layer and glass nanowell to mimic the free diffusion experiment (***Figure 3***—***Figure Supplement 7***). A 50 nm thin glass layer was added at the edge of the simulated volume (at *z* ≈ −4 µm) to account for the glass-water interface. The signal enhancement factor at each z-position was computed as the ratio of the detected signal in the nanowell and the free diffusion value:

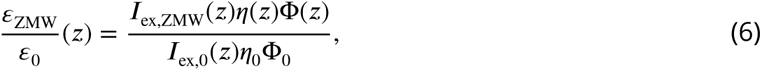

where Φ_0_ is the reference quantum yield. Here, we assume that 50% of the signal is detected in the free diffusion case (*η*_0_ = 0.5) and neglect detection losses due to the limited numerical aperture of the objective lens which are assumed to be identical for the compared conditions. The predicted average enhancement factors were then computed as the signal-weighted average along the central pore axis (*x* = 0, *y* = 0) from the bottom of the well of depth ℎ towards the end of the simulation box at height 200 nm:

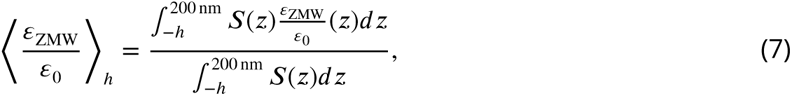

where *S*(*z*) is the detected signal as defined in eq. 5. See ***Figure 3***—***Figure Supplement 7*** for details.

### Live cell imaging on Pd ZMWs

#### Cell lines

Human U2OS (ATCC, HTB-96) and HEK 293T (ATCC, CRL-3216) cells were used for imaging and lentivirus production, respectively. They were grown in DMEM (4.5 g/L glucose, Gibco) with 5 % fetal bovine serum (Sigma-Aldrich) and 1 % penicillin / streptomycin (Gibco) and maintained at 37 °C with 5 % CO_2_. The cell lines were confirmed to be mycoplasma-free. Cell lines stably expressing transgenes were generated via lentiviral transduction. Lentivirus was produced by transfecting HEK 293T cells with polyethylenimine (Polysciences Inc.) and packaging vectors (psPAX2, pMD2.g) and the lentiviral plasmid of interest. The viral supernatant was collected 72 h after transfection. Cells were seeded for infection at ≈ 35 % confluency 24 h prior to lentivirus addition. The cells were spin-infected with the viral supernatant and Polybrene (10 µg/ml) for 90 min at 2000 rpm at 32 °C, then cultured for 48 h. Monoclonal cell lines expressing the transgene were isolated by single cell sorting into 96-well plates via FACS. The TfR coding sequence was amplified from Addgene plasmid # 133451.

#### Cell culture for imaging

Cells were seeded on ZMW coverslips in a 6-well plate at 40 % to 45 % confluency 1 day before the imaging experiment. The cell culture medium was replaced with imaging medium (pre-warmed CO_2_-independent Leibovitz’s-15 medium (Gibco) with 5 % fetal bovine serum and 1 % penicillin / streptomycin) 30 min prior to imaging. All live-cell imaging experiments were performed at 37 °C. For experiments with HaloTag expressing cell lines, they were incubated with 5 nM JFX650-Halo ligand in pre-warmed Leibovitz’s L15 medium for 10 min and washed three times.

#### Microscope and image acquisition

Live-cell imaging experiments were performed using a Nikon TI inverted microscope equipped with a TIRF illuminator, perfect focus system and NIS Element Software. A Nikon CFI Apochromat TIRF 100X 1.49 NA oil-immersion objective was used. The microscope was equipped with a temperature-controlled incubator. Bright-field and fluorescence images at each ZMW array position were recorded using an Andor iXon Ultra 888 EMCCD camera.

### Data analysis

#### Single-molecule fluorescence experiments

Fluorescence correlation spectroscopy (FCS) and lifeitme analysis was performed using the PAM software package (***Schrimpf et al., 2018***). Autocorrelation functions *G*(*t*_*c*_) were fit to a standard model function for 3D diffusion:

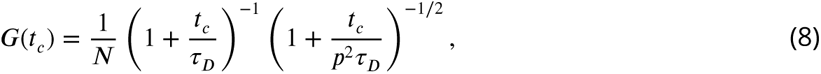

where *t*_*c*_ is the correlation time, *N* is the average number of particles in the observation volume, *τ*_*D*_ is the diffusion time, and *p* is a geometric factor that accounts for the axial elongation of the confocal volume (*p* = 3.4). While, strictly speaking, the 3D diffusion model is not applicable for the complex geometries in this study, we apply it here as a simple means to extract the amplitude and average decay time of the curves. The molecular brightness *ε* was calculated from the average signal ⟨*I*⟩ as *ε* = ⟨*I*⟩ ∕*N*. The effective volume was computed using the known concentration *c* of the fluorophore as *V*_FCS_ = *N*∕(*N*_*A*_*c*), where *N*_*A*_ is Avogadro’s number. Fluorescence decays of Alexa488 were fitted to a single-exponential decay that was convoluted with the instrument response function. Decays for JFX650 generally required two lifetimes to achieve a good fit, of which we report the average. The second component most likely originates from a fraction of dyes that were sticking to the surface.

#### Cell imaging

The images were analysed using custom-written software for MATLAB. The program automatically determined the positions of pores from bright-field images and calculated the fluorescence intensity of each pore from the fluorescence images. The BFP and JFX650-Halo fluorescence signals were obtained by calculating the mean intensity over a 7×7 pixel area around the pore and subtracting the background intensity determined from the outer edge pixels of the pore. To analyze the fraction of pores occupied by cells for each pore size (***Figure 4*** C), we estimated the area occupied by cells by manually determining an outline containing connected areas of occupied wells from 6 images. We analyzed the BFP intensity of each well in this area, with a total of 7367 wells analyzed. As a negative control, we measured the BFP intensity of the pores where no cells were present and set a threshold to determine positive BFP intensity pores. The total number of pores with a positive BFP signal above the threshold was 1890 (***Figure 4***—***Figure Supplement 1*** C). This panel illustrates the fraction of pores with positive BFP intensity among the analyzed pores for each pore size. For the analysis of the JFX650-Halo time traces, a hidden Markov model (vbFRET algorithm, (***Bronson et al., 2009***)) with the default setting of the algorithm was used to assign on and off states of the Halo signal.

## Data availability

All data underlying this study is made available in an open repository (***Yang et al., 2023***)

## Author contributions

C.D. and M.E.T. designed and supervised research. N.K. designed and fabricated nanostructures. A.B. and N.K. performed in vitro experiments. A.B. performed electromagnetic simulations and analyzed in vitro experiments. S.Y. performed and analyzed cell experiments. A.B., N.K., and S.Y. wrote the initial draft. All authors contributed to the final manuscript.

Taxonomy according to CRediT: A.B.: Methodology, Validation, Formal Analysis, Investigation, Writing — Original Draft Preparation, Writing — Review & Editing, Visualization, Funding Acquisition C.D.: Conceptualization, Writing — Review & Editing, Supervision, Project Administration, Funding Acquisition M.E.T.: Conceptualization, Writing — Review & Editing, Supervision, Project Administration, Funding Acquisition N.K.: Methodology, Validation, Investigation, Resources, Writing — Original Draft Preparation, Writing — Review & Editing, Visualization, Project Administration S.Y.: Methodology, Validation, Formal Analysis, Investigation, Writing — Original Draft Preparation, Writing — Review & Editing, Visualization, Funding Acquisition

## Acknowledgments

A.B, N.K. and C.D. acknowledge financial support by the NWO program OCENW.GROOT.2019.068, ERC Advanced Grant no. 883684, and the NanoFront and BaSyC programs of NWO/OCW. A.B. acknowledges funding from the European Union’s Horizon 2020 research and innovation program under the Marie Skłodowska-Curie Grant agreement no. 101029907. M.E.T., S.Y. were supported by the Oncode Institute, which is partly funded by the Dutch Cancer Society (KWF). S.Y. acknowledges funding from the European Union’s Horizon 2020 research and innovation programm under the Marie Skłodowska-Curie grant agreement no. 101026470. We thank Iris Bally and Ive Logister for help with experiments and Micha Müller for sharing CM40TM-BFP plasmid.

---

**Figure 1—Figure Supplement 1.**
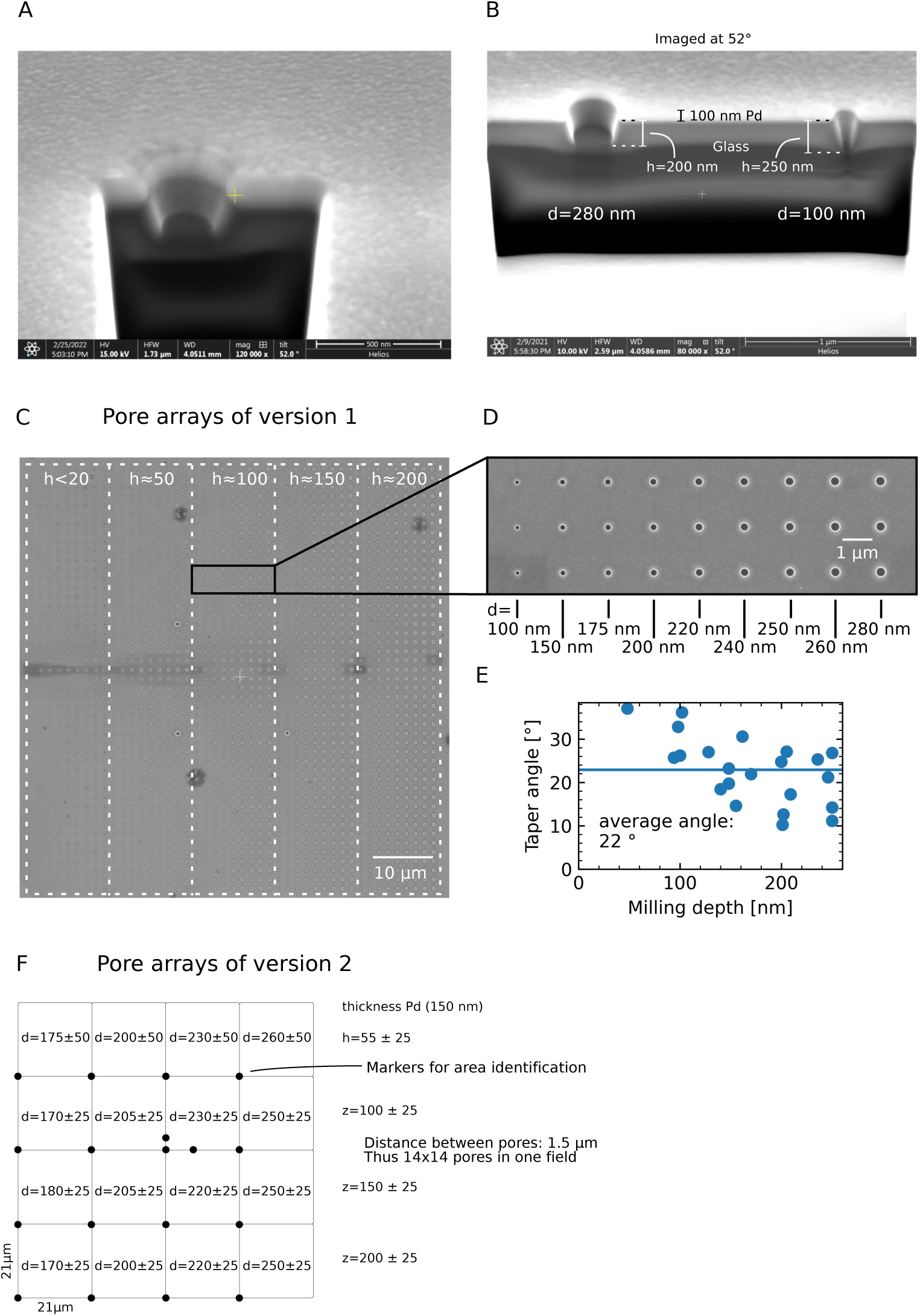
Additional SEM images and layouts of version 1 and 2 arrays. **A,B:** SEM images of overmilled Pd ZMWs, imaged under 52 °. Pd shows up as a bright layer and glass as a dark layer. **C:** An SEM image of a Pd ZMW array of version 1 containing five regions of different milling depths (h), each made from nine rows of pores with varying diameter. **D:** Zoom in of C. **E:** Taper angle vs. milling depth. For deeper pores, the edges were more perpendicular with an average of 22 ° (horizontal line). **F:** Diameters (d) and depths (h) in nm of ZMW arrays for arrays of version 2 together with their uncertainty (estimated from measuring several pores).

**Figure 2—Figure Supplement 1.**
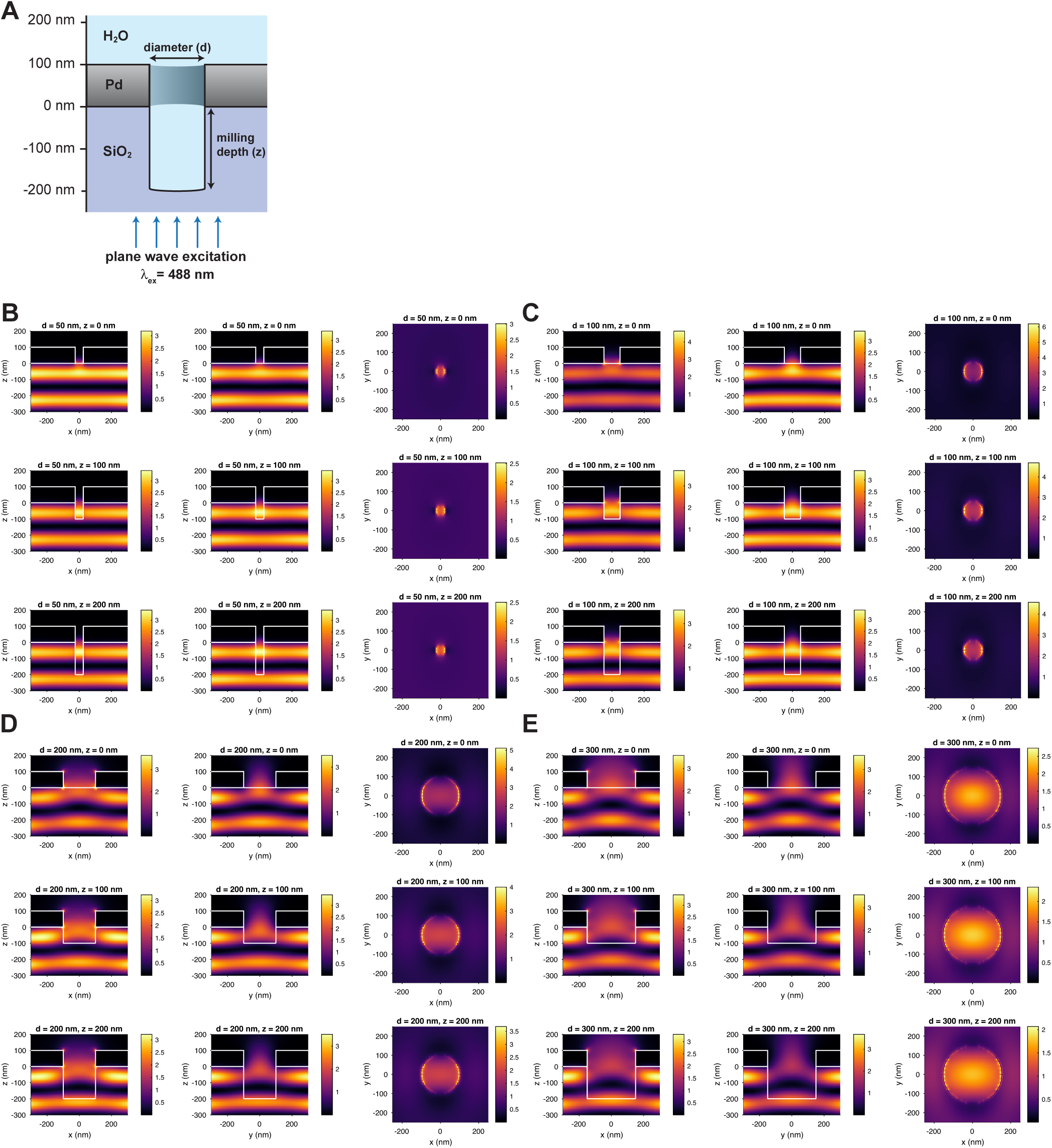
Excitation field intensity distributions from FDTD simulations under widefield excitation at. λ_ex_ = **488 nm. A:** Schematic of the simulation setup. **B-E:** Excitation field intensity distributions |*E*|^2^ in *V*^2^∕*m*^2^ in the x-z (left), y-z (middle) and x-y plane at the entrance to the ZMW (right) at pore diameters d of 50 nm (B), 100 nm (C), 200 nm (D), and 300 nm (E) and milling depths h of 0 nm (top), 100 nm (middle), and 200 nm (bottom). The electric field is polarized along the x-axis.

**Figure 2—Figure Supplement 2.**
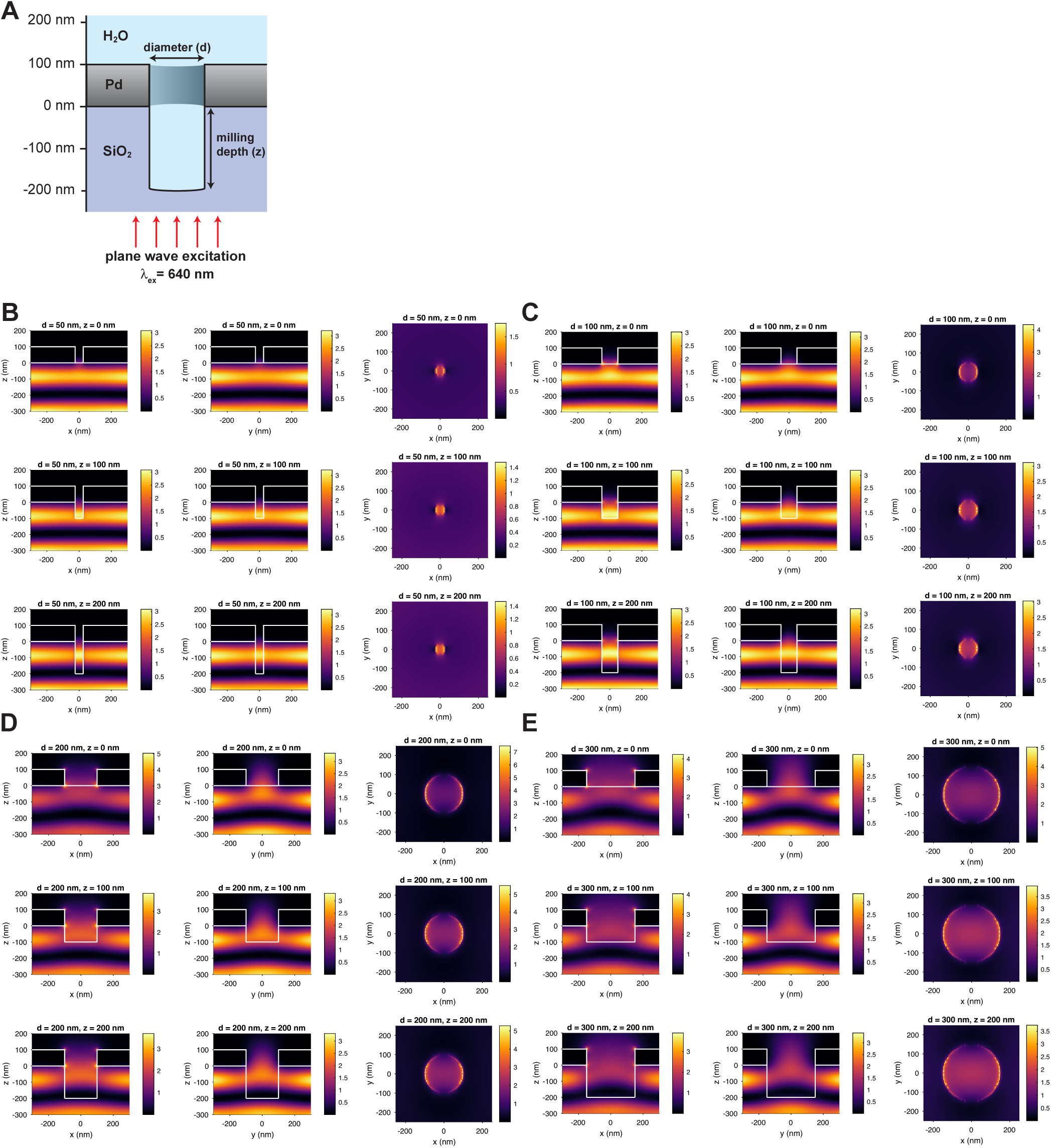
Excitation field intensity distributions from FDTD simulations under widefield excitation at λ_ex_ = 640 nm. **A:** Schematic of the simulation setup. **B-E:** Excitation field intensity distributions |*E*|^2^ in *V*^2^∕*m*^2^ in the x-z (left), y-z (middle) and x-y plane at the entrance to the ZMW (right) at pore diameters d of 50 nm (B), 100 nm (C), 200 nm (D), and 300 nm (E) and milling depths h of 0 nm (top), 100 nm (middle), and 200 nm (bottom). The electric field is polarized along the x-axis.

**Figure 2—Figure Supplement 3.**
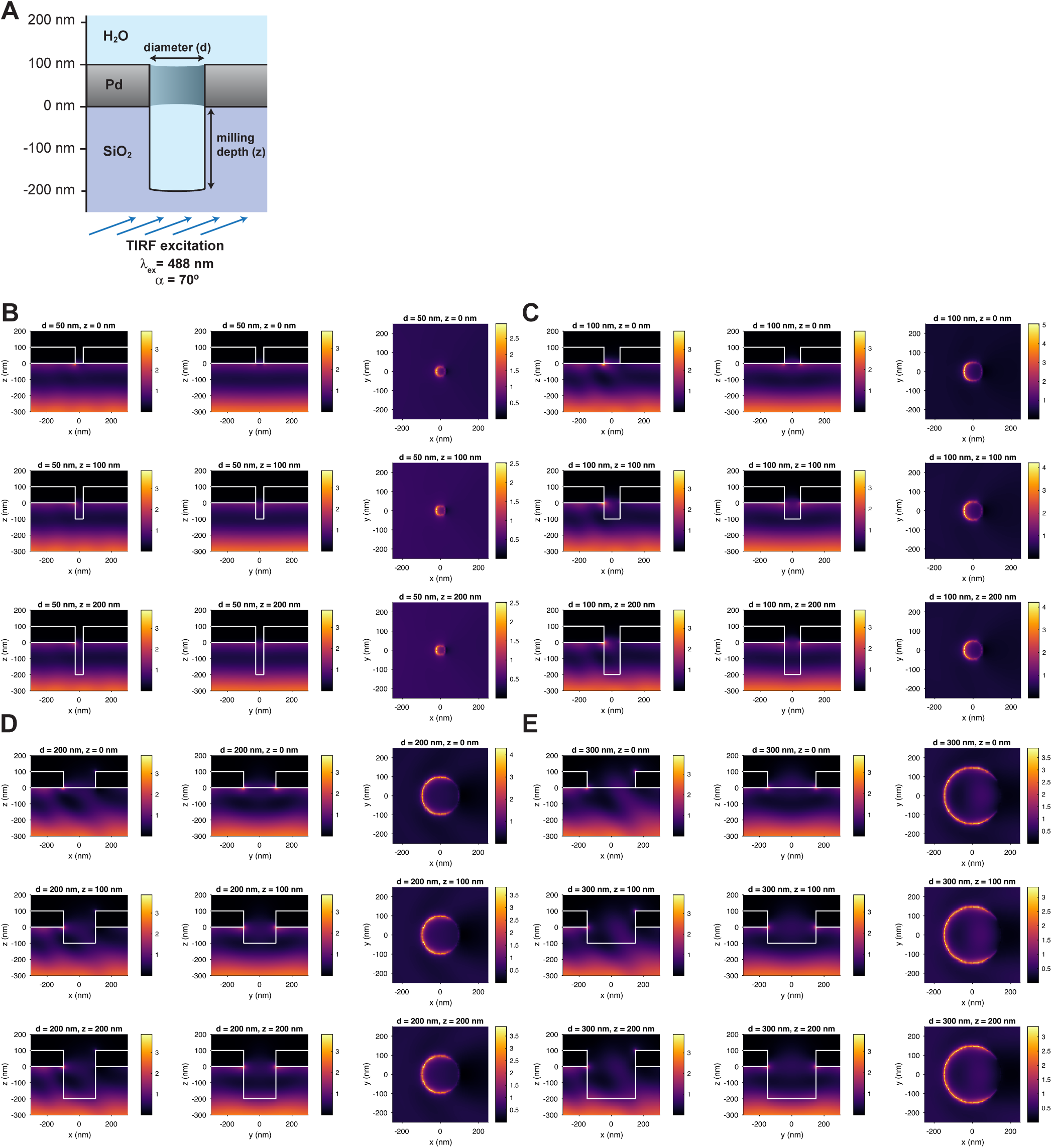
Excitation field intensity distributions from FDTD simulations under TIRF illumination at an angle of 70° at λ_ex_ = 488 nm. **A:** Schematic of the simulation setup. **B-E:** Excitation field intensity distributions |*E*|^2^ in *V*^2^/*m*^2^ in the x-z (left), y-z (middle) and x-y plane at the entrance to the ZMW (right) at pore diameters d of 50nm (B), 100nm (C), 200nm (D), and 300nm (E) and milling depths h of 0nm (top), 100nm (middle), and 200nm (bottom). The electric field is polarized along the x-axis.

**Figure 2—Figure Supplement 4.**
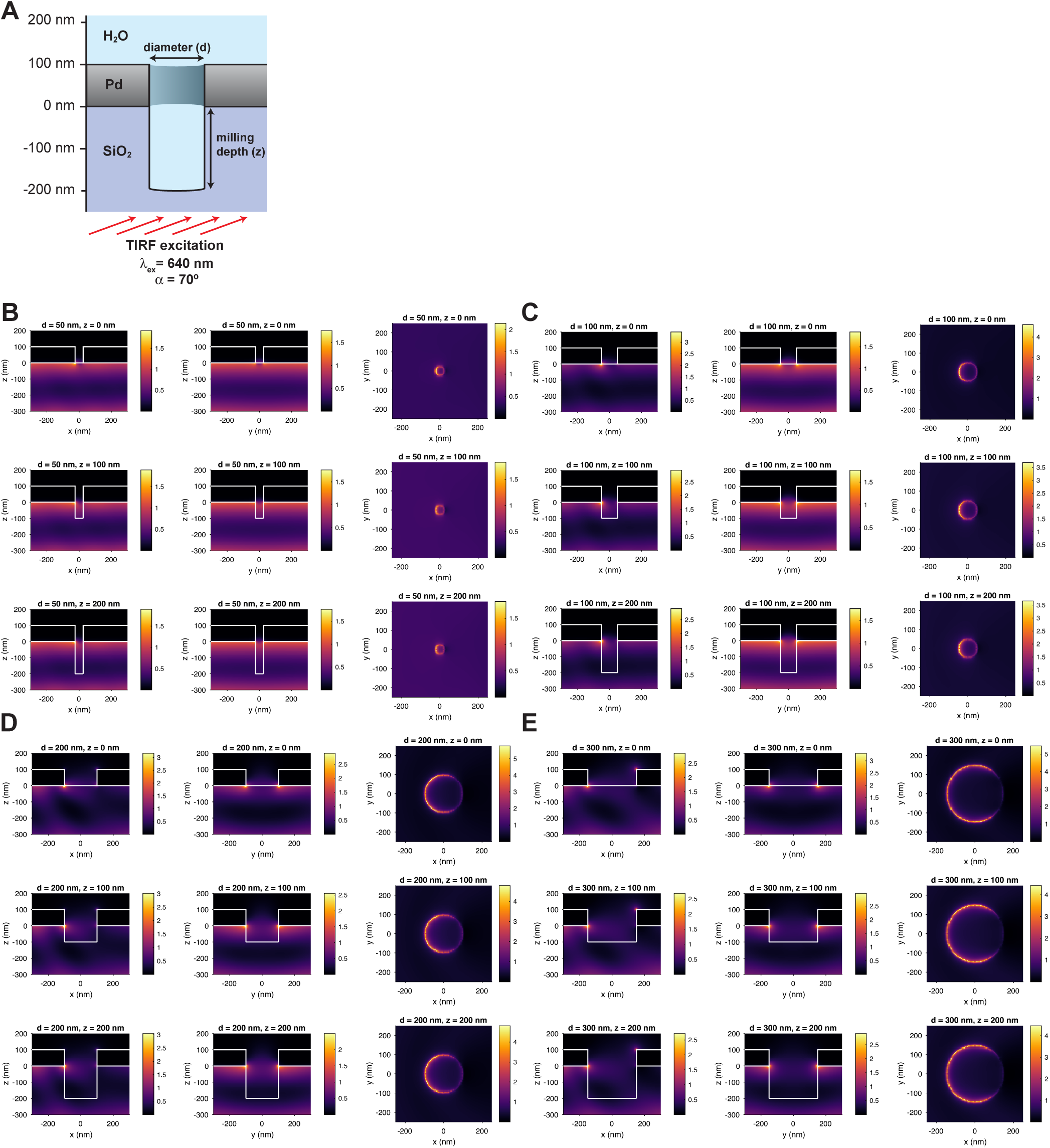
Excitation field intensity distributions from FDTD simulations under TIRF illumination at an angle of 70° at λ_ex_ = 640 nm. **A:** Schematic of the simulation setup. **B-E:** Excitation field intensity distributions |*E*|^2^ in *V*^2^∕*m*^2^ in the x-z (left), y-z (middle) and x-y plane at the entrance to the ZMW (right) at pore diameters d of 50 nm (B), 100 nm (C), 200 nm (D), and 300 nm (E) and milling depths h of 0 nm (top), 100 nm (middle), and 200 nm (bottom). The electric field is polarized along the x-axis.

**Figure 2—Figure Supplement 5.**
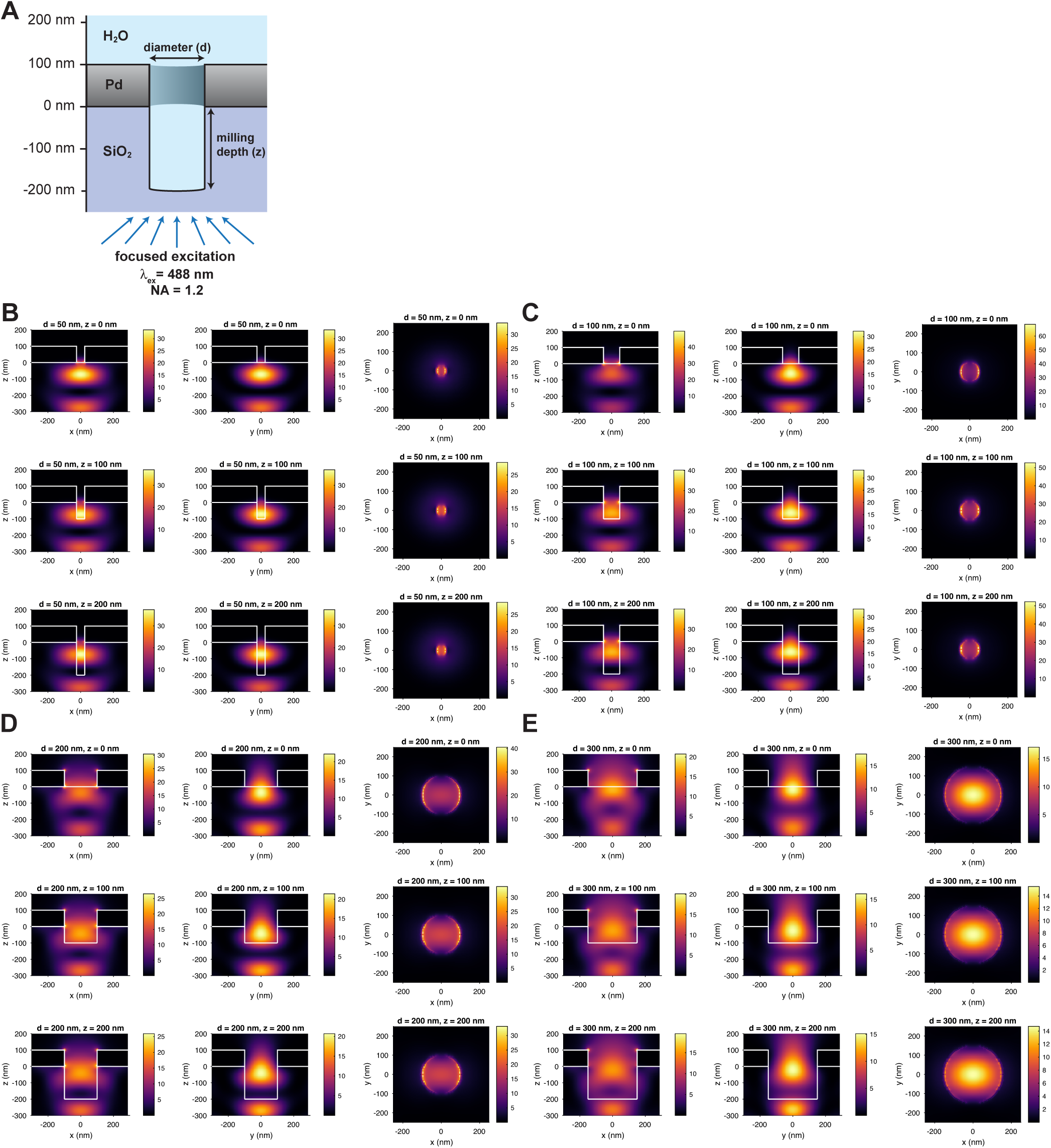
Excitation field intensity distributions from FDTD simulations under excitation by a focused Gaussian beam at NA = 1.2 and a wavelength of λ_ex_ = 488 nm. **A:** Schematic of the simulation setup. **B-E:** Excitation field intensity distributions |E| 2 in *V*^2^∕*m*^2^ in the x-z (left), y-z (middle) and x-y plane at the entrance to the ZMW (right) at pore diameters d of 50 nm (B), 100 nm (C), 200 nm (D), and 300 nm (E) and milling depths h of 0 nm (top), 100 nm (middle), and 200 nm (bottom). The electric field is polarized along the x-axis.

**Figure 2—Figure Supplement 6.**
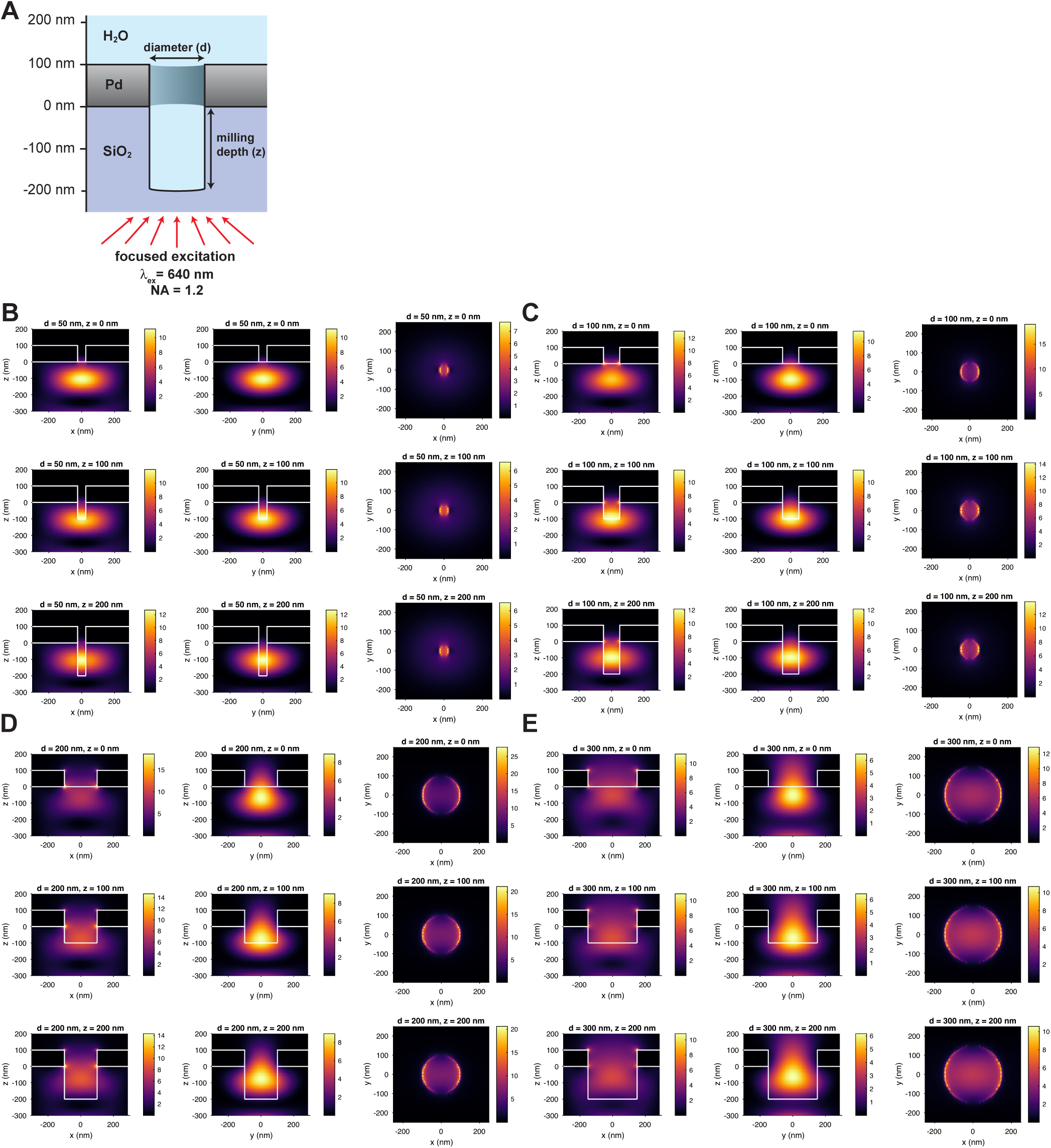
Excitation field intensity distributions obtained from FDTD simulations under excitation by a focused Gaussian beam at NA = 1.2 and λ_ex_ = 640 nm. **A:** Schematic of the simulation setup. **B-E:** Excitation field intensity distributions |*E*|^2^ in *V*^2^∕*m*^2^ in the x-y (left), y-z (middle) and x-y plane at the entrance to the ZMW (right) at pore diameters d of 50 nm (B), 100 nm (C), 200 nm (D), and 300 nm (E) and milling depths h of 0 nm (top), 100 nm (middle), and 200 nm (bottom). The electric field is polarized along the x-axis.

**Figure 2—Figure Supplement 7.**
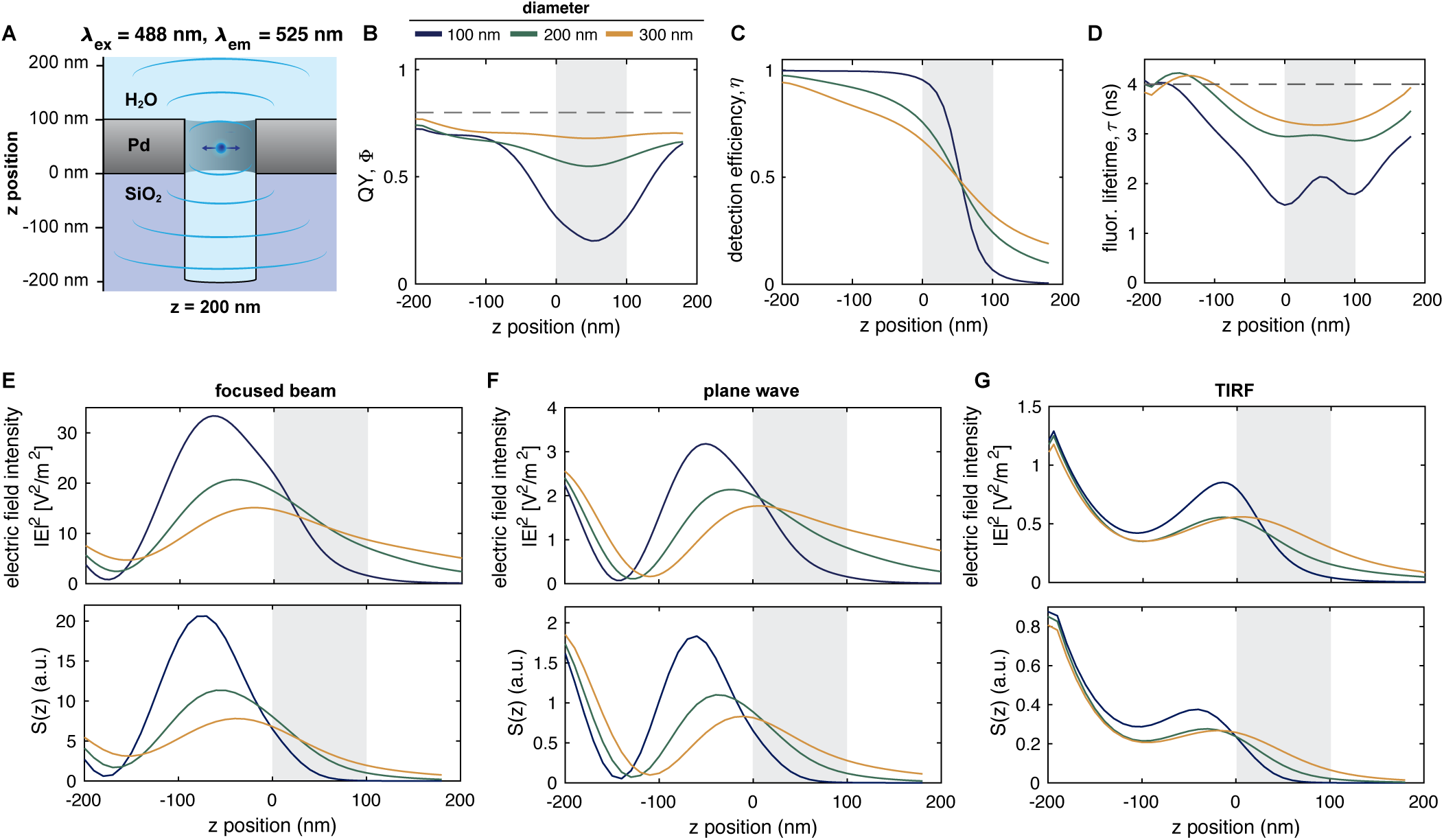
FDTD simulations of fluorescence emission and detected signal from overmilled ZMWs for Alexa488. **A:** Schematic of the simulation setup. **B-D:** Computed quantum yield Φ, detection efficiency η, and fluorescence lifetime τ profiles of the dye Alexa488 as a function of the z position. Dashed lines indicate the values in the absence of a ZMW. **E-G:** Z-profiles of the excitation intensity profiles along the central pore axis (top) and the total detected signal S(z) (bottom) under excitation by a focused Gaussian beam (E), plane wave (i.e., widefield) (F) or excitation under TIRF angle (G). The position of the metal membrane is indicated as a gray shaded area.

**Figure 2—Figure Supplement 8.**
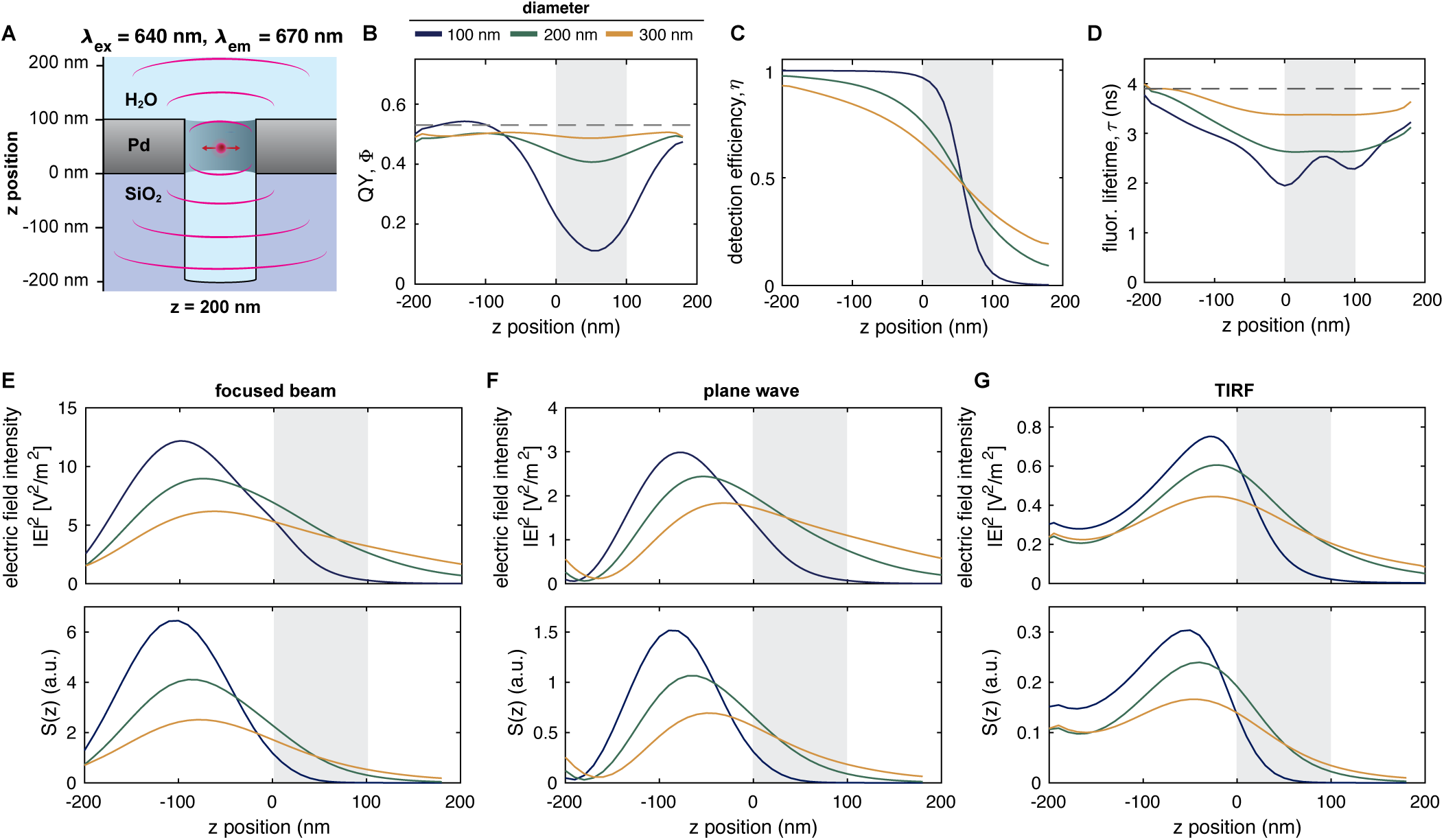
FDTD simulations of fluorescence emission and detected signal from overmilled ZMWs for JFX650. **A:** Schematic of the simulation setup. **B-D:** Computed quantum yield Φ, detection efficiency η, and fluorescence lifetime τ profiles of the dye JFX650 as a function of the z position. Dashed lines indicate the values in the absence of a ZMW. **E-G:** Z-profiles of the excitation intensity profiles along the central pore axis (top) and the total detected signal S(z) (bottom) under excitation by a focused Gaussian beam (E), plane wave (i.e., widefield) (F) or excitation under TIRF angle (G). The position of the metal membrane is indicated as a gray shaded area.

**Figure 2—Figure Supplement 9.**
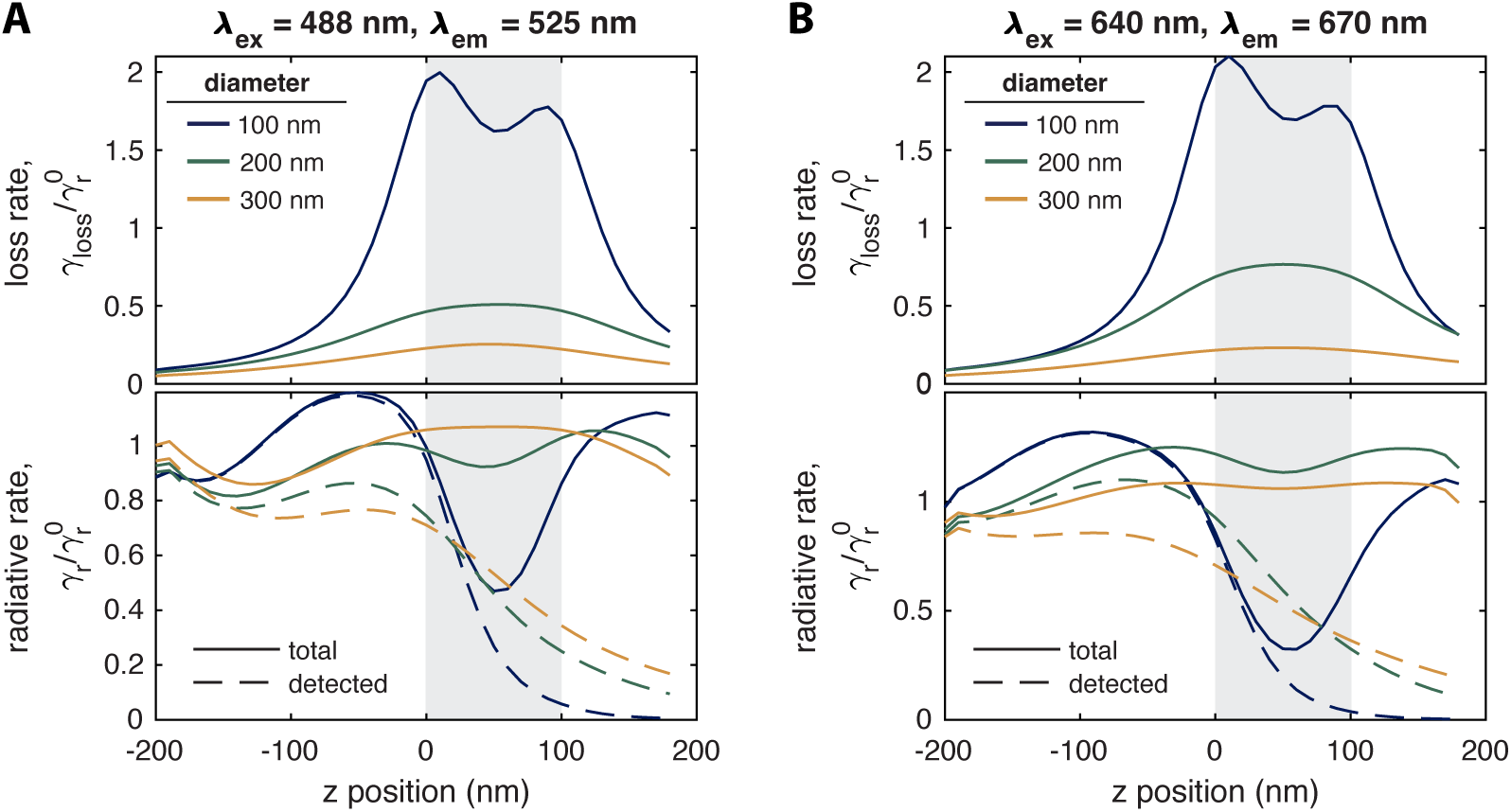
Overview of radiative and non-radiative rates obtained from FDTD simulations of fluorescence emission within overmilled ZMWs. Shown are the nonradiative loss rate (top) and radiative rate (bottom) in the presence of the metal nanostructure for the dyes Alexa488 (A) and JFX650 (B). The radiative rate towards the detection side is given as a dashed line, from which the detection efficiency is computed. Note that the given rates are normalized to the rates in the absence of the ZMW as described in the methods.

**Figure 2—Figure Supplement 10.**
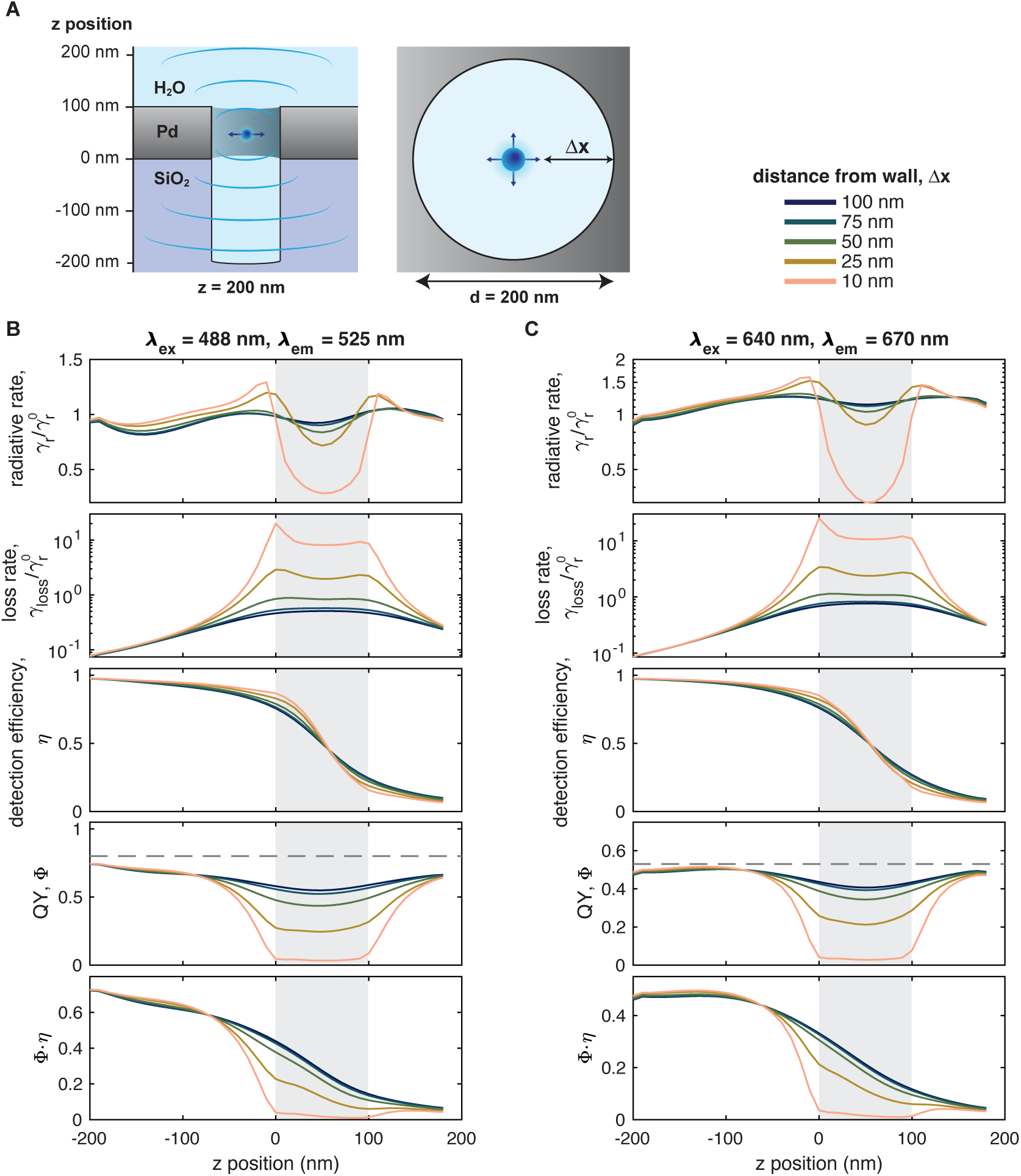
FDTD simulations of dipole emission as a function of the distance to the pore walls. **A:** Schematic of the simulation setup. The dipole was placed at varying distances Δx from the pore walls and the emission was monitored as a function of the z-position. **B,C:** Z-profiles of the normalized radiative and loss rates, the detection efficiency η, the quantum yield Φ, and the product of the detection efficiency and quantum yield, Φ ⋅ η for Alexa488 (B) and JFX650 (C) at the indicated excitation and emission wavelengths. The quantum yield of the free dye is shown as a dashed line. The dipole emission within the overmilled volume in the glass was found to be approximately independent of the lateral displacement within the pore at distance of ≈ 25 nm away from the ZMW. Within the ZMW, the non-radiative rate is strongly increased as the dipole approaches the pore wall, with significant non-radiative losses occurring at distances below 25 nm that result in a reduction of the quantum yield. Note that the loss rate is given on a log scale. The position of the palladium layer is indicated as a gray shaded area.

**Figure 2—Figure Supplement 11.**
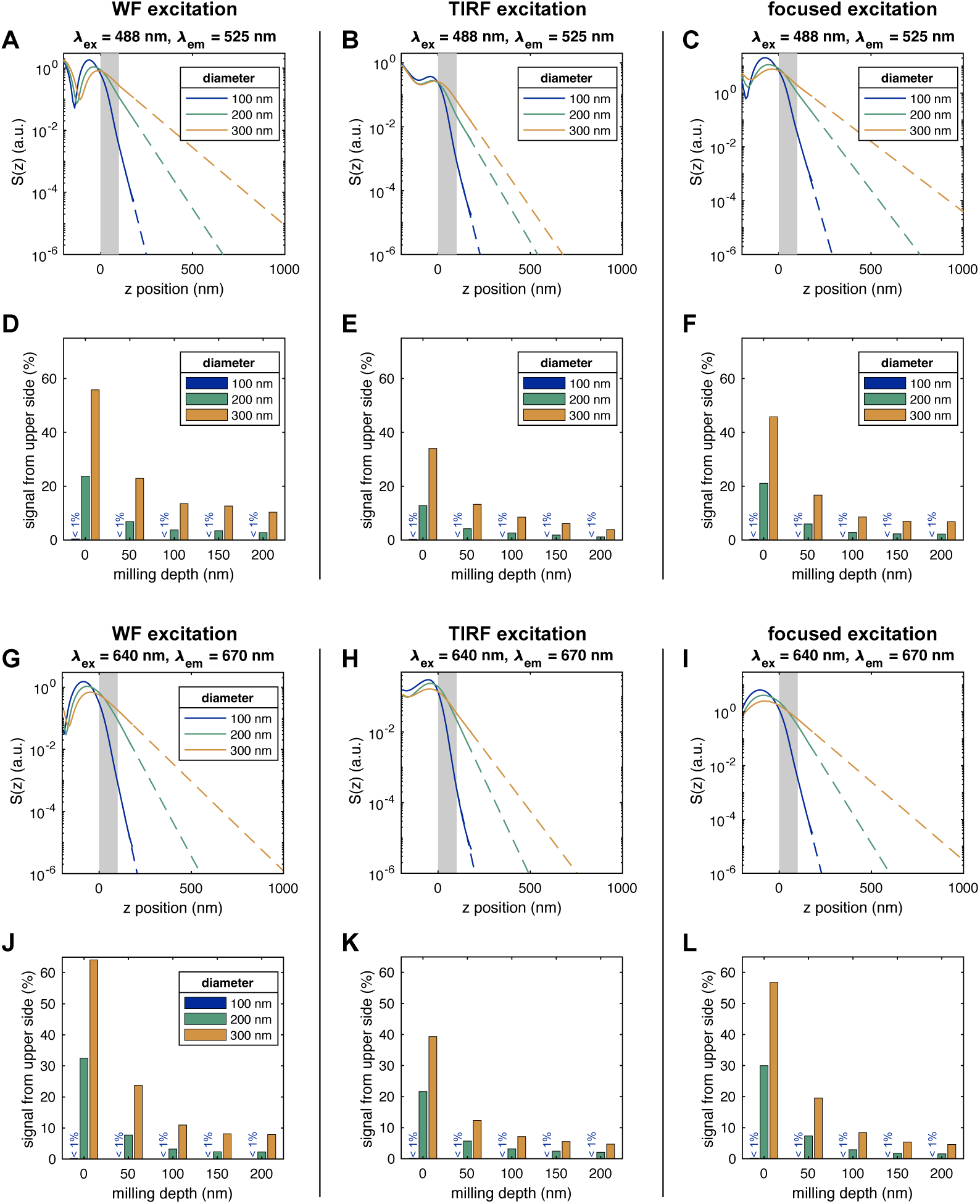
Estimation of background signal from FDTD simulations for the dyes Alexa488 (A-F) and JFX650 (G-L) under the different excitation modes. **A-C, G-I:** The detected signal S(z) obtained for widefield (WF), TIRF, and focused excitation (solid lines) is extrapolated by fitting the signal profile in the z-range from 100 nm to 200 nm to an exponential decay (dashed lines). **D-F, J-L:** From the extrapolated signal profiles, the amount of signal detected from the upper side (above a z position of 100 nm) is estimated for different pore diameters and milling depths. The bars for 100 nm are barely visible due to their small height.

**Figure 3—Figure Supplement 1.**
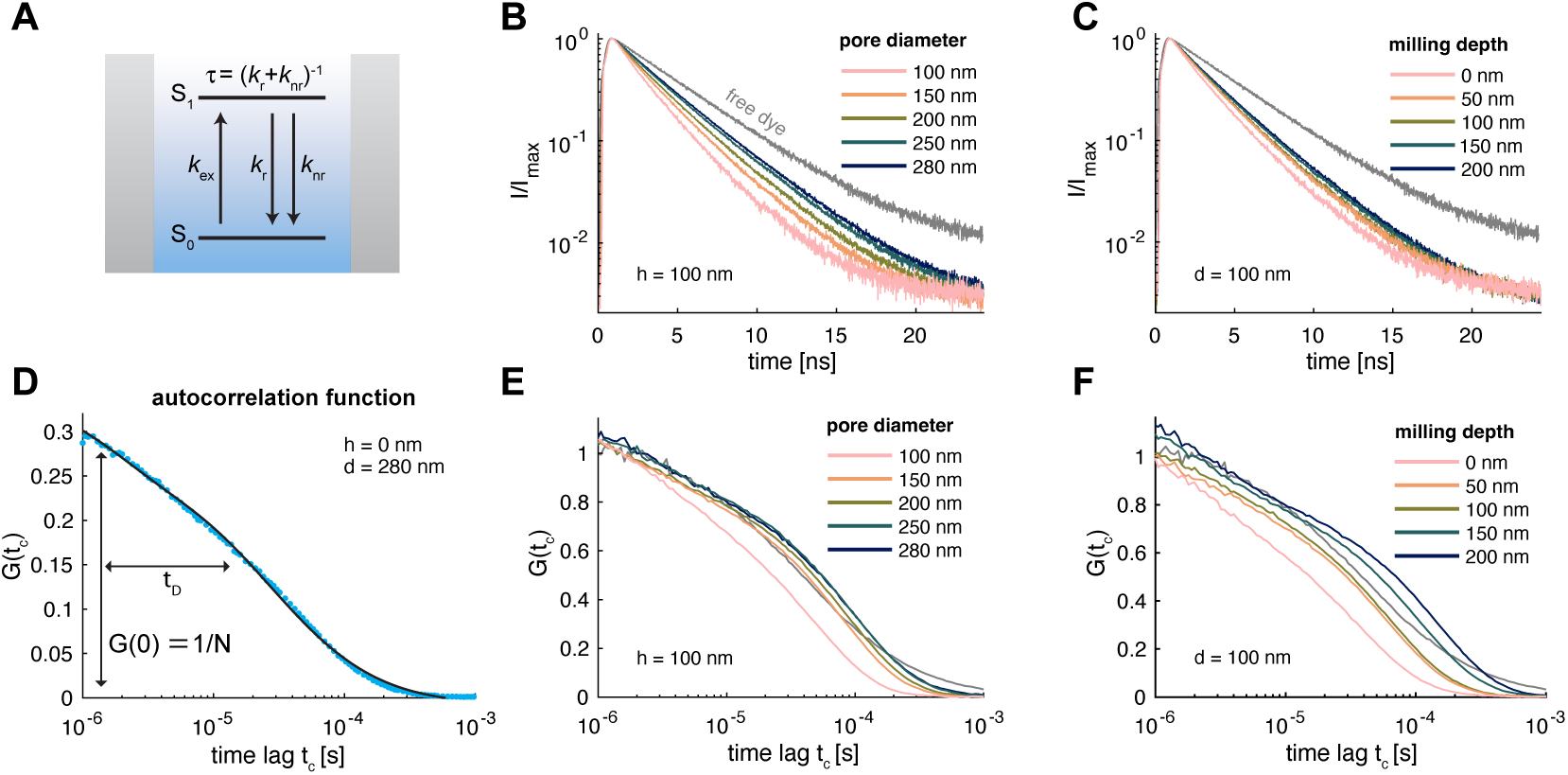
Experimental characterization of photophysics and diffusion within ZMWs. **A:** Jablonski scheme of the photophysics in the ZMW. Dyes are radiatively excited from the electronic ground state S_0_ to the first excited state S_1_ with rate k_ex_, from where relaxation can occur radiatively (k_r_) or non-radiatively (k_nr_). In the waveguide, all displayed rates change due to excitation field enhancement, plasmonic coupling, and metal-induced quenching. The excited state lifetime reports on the sum of the radiative and non-radiative rates. **B,C:** Fluorescence decays acquired at different pore diameters for a constant milling depth of 100 nm (B) and at different milling depths for a constant pore diameter of 100 nm (C). **D:** Autocorrelation function of the fluorescence time trace shown in Figure 3 C. The FCS analysis informs on the average number of particles (N) and the residence time of molecules within the ZMW (t_D_). **E,F:** FCS curves of the data shown in Figure 3 D-F. In B,C,E, and F, the curves for the free dye obtained from a free diffusion experiment are shown in gray.

**Figure 3—Figure Supplement 2.**
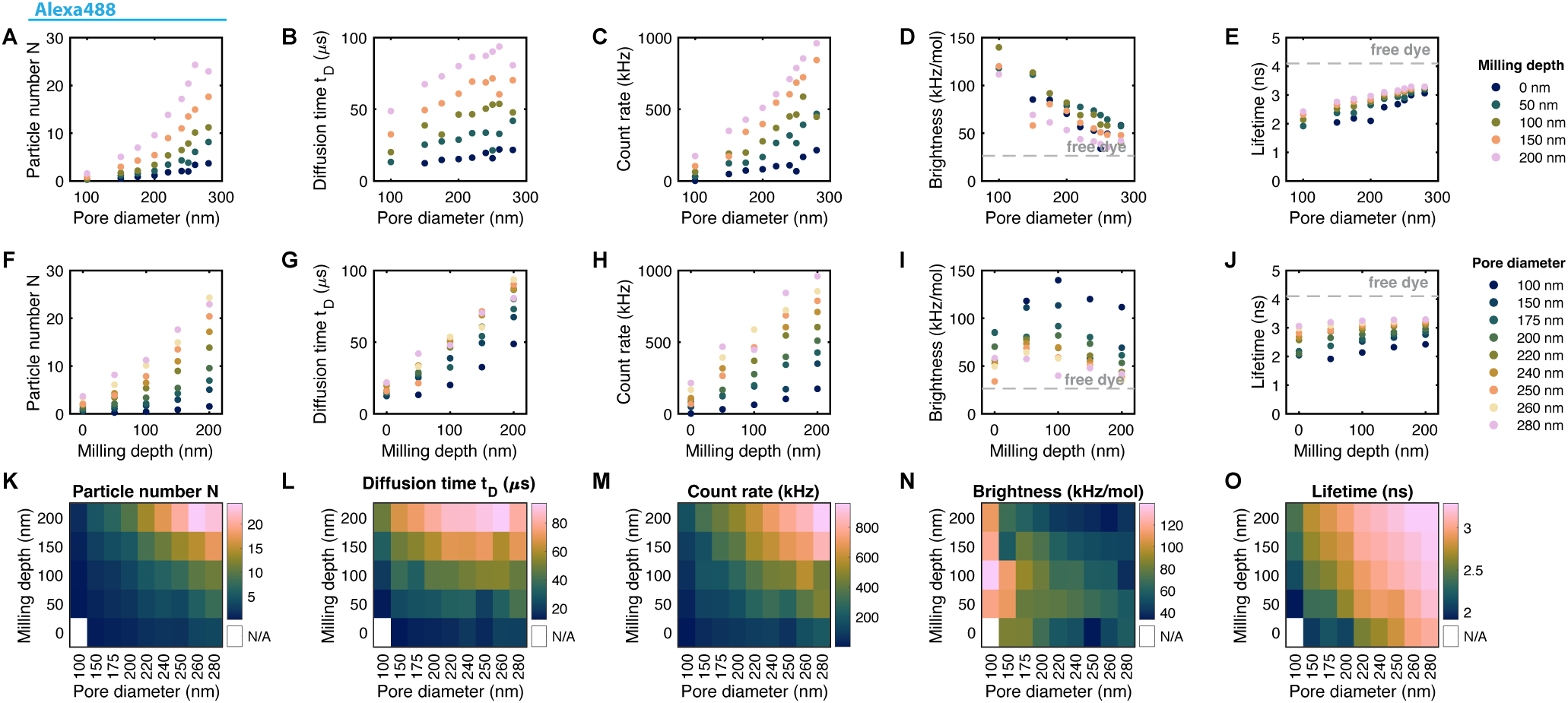
Extracted parameters for the dye Alexa488 in overmilled Pd ZMWs. Shown are the estimated particle number N, diffusion time t_D_, count rate, molecular brightness ε_ZMW_, and fluorescence lifetime τ as a function of the pore diameter at constant milling depth (A-E), as a function of the milling depth at constant pore diameter (F-J), and as heatmap plots (K-O). The molecular brightness and fluorescence lifetime of the free dye are indicated by gray dashed lines.

**Figure 3—Figure Supplement 3.**
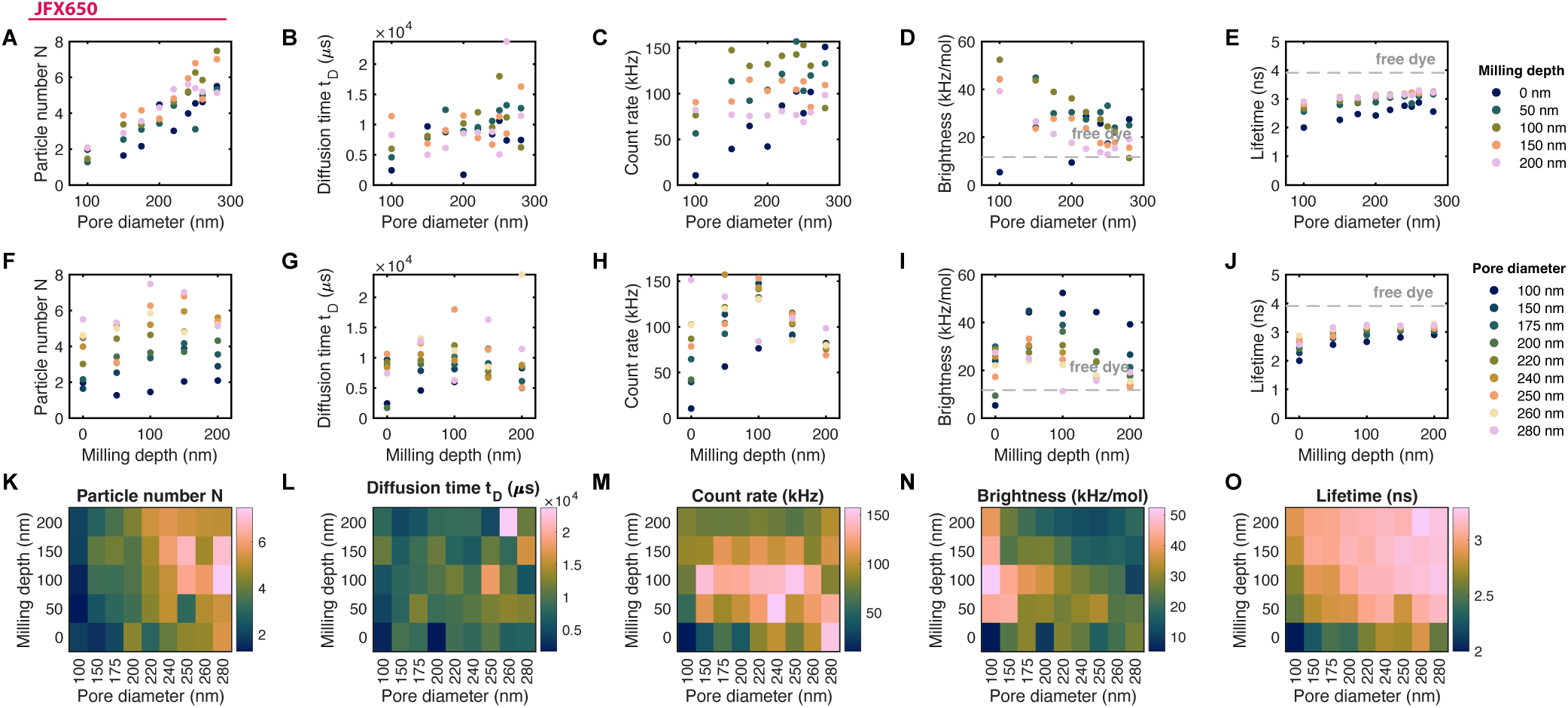
Extracted parameters for the dye JFX650 in overmilled Pd ZMWs. Shown are the estimated particle number N, diffusion time t_D_, count rate, molecular brightness ε_ZMW_, and fluorescence lifetime τ as a function of the pore diameter at constant milling depth (A-E), as a function of the milling depth at constant pore diameter (F-J), and as heatmap plots (K-O). The molecular brightness and fluorescence lifetime of the free dye are indicated by gray dashed lines. The robustness of the FCS analysis is markedly reduced compared to the Alexa488 dye due to significant sticking of the JFX650 dye to the glass and metal surfaces, as evident from the drastically prolonged diffusion times (B,G,L).

**Figure 3—Figure Supplement 4.**
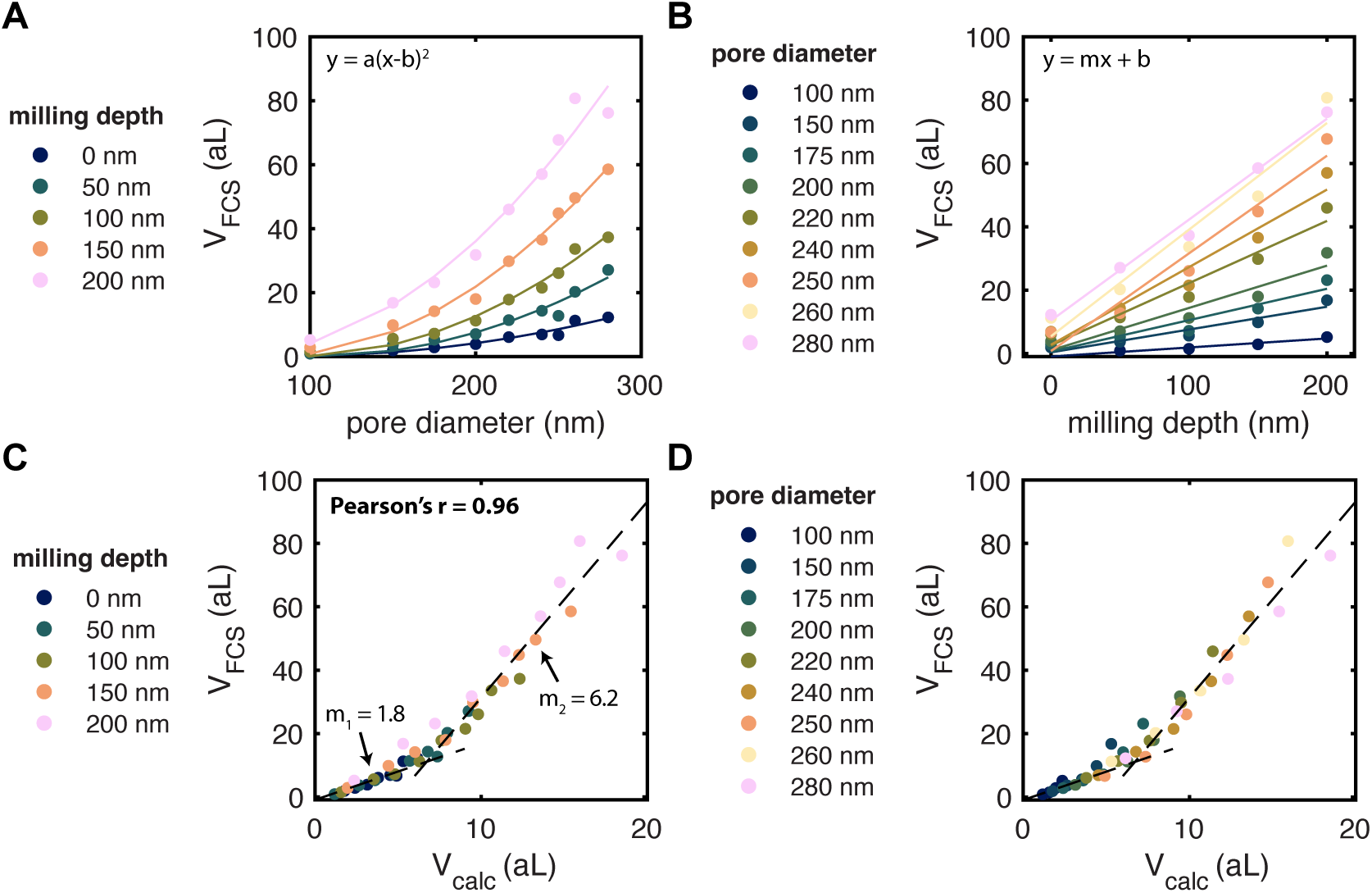
Quantification of the observation volume in overmilled ZMWs. **A,B:** Estimated effective volumes V_FCS_ as a function of the pore diameter (A) and milling depth (B). The effective volume V_FCS_ was estimated from the particle number N determined from FCS as V_FCS_ = N∕c, where c is the concentration of the dye (c = 500 nM). Note that the effective volume in FCS corresponds to a hypothetical volume with constant signal that contains N particles (***Xu and Webb, 2002***; ***Levene et al., 2003***). The effective volume shows a quadratic scaling with the pore diameter and linear scaling with the milling depth, as expected for the approximately cylindrical volume of the overmilled ZMWs. **C,D:** Comparison of the calculated volume of the overmilled aperture in the glass, V_calc_, and the experimentally determined effective volume, V_FCS_, color coded by milling depth (C) and pore diameter (D). V_calc_ is given by V_calc_ = (π∕4)d^2^(z + l), where d is the diameter, z the overmilling depth, and l is the thickness of the Pd layer (l = 100 nm). An excellent correlation is observed between the two quantities (Pearson’s correlation coefficient r = 0.96), however the measured volumes V_FCS_ are consistently overestimated by a factor of ≈ 2 for small pores (V_calc_ ≤ 5 aL) and ≈ 6 for large pores (V_calc_ ≥ 10 aL). Note that this overestimation is also present at small pore diameters where only little signal is detected from within the ZMW. Similar overestimation of the particle numbers within ZMW by FCS have previously been reported and attributed to a contribution of constant signal from many dim particles outside of the ZMW (***Levene et al., 2003***; ***Rigneault et al., 2005***; ***Lenne et al., 2008***; ***Wu et al., 2019***).

**Figure 3—Figure Supplement 5.**
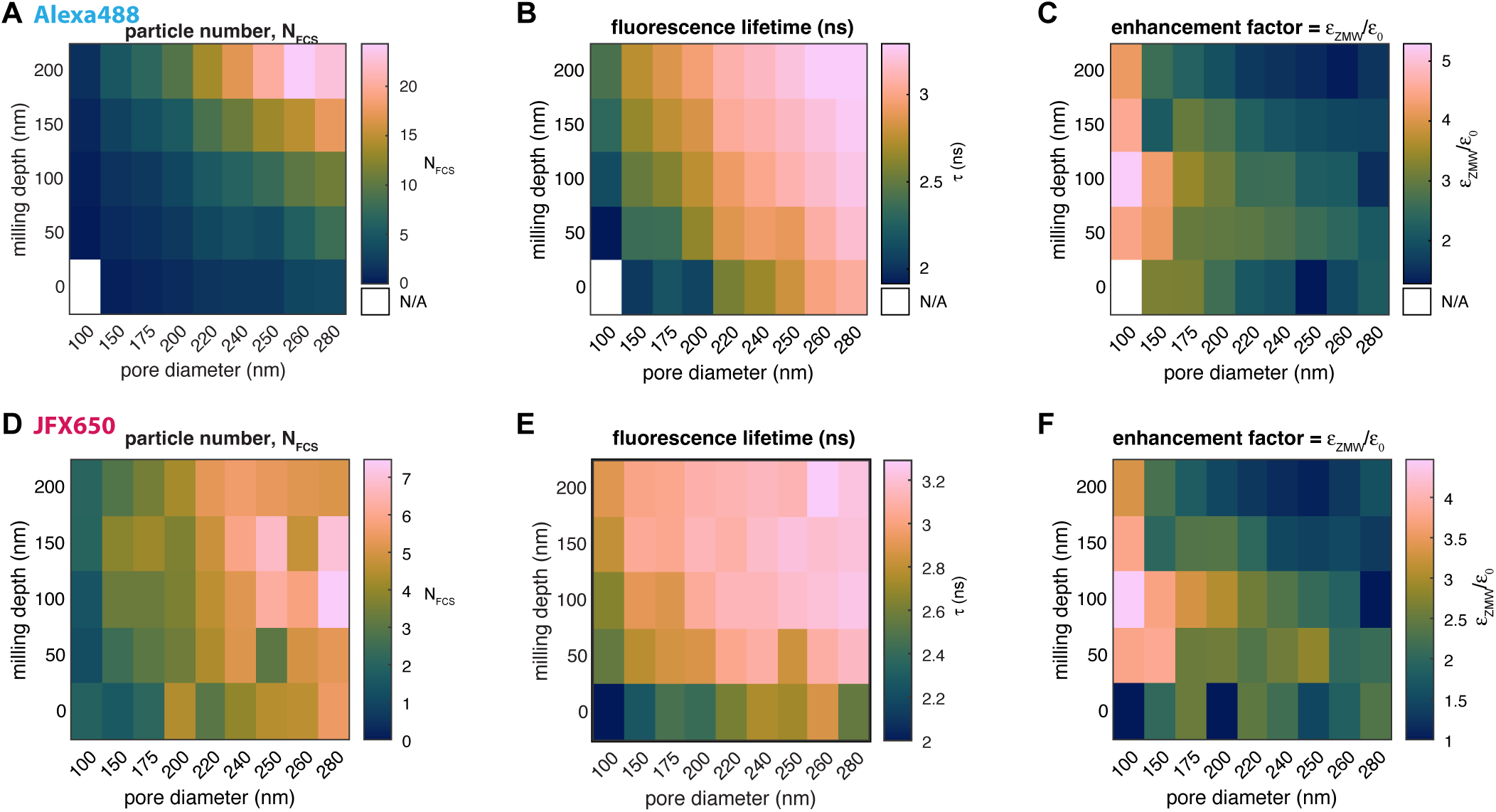
Comparison of extracted parameters for the dyes Alexa488 and JFX650 in ZMW. The heatmap plots show the average number of particles in the observation volume N, fluorescence lifetime τ, and signal enhancement factor for the dyes Alexa488 (A-C) and JFX650 (D-F). The enhancement factor is defined as the ratio of the counts per molecule in the ZMW compared to free diffusion, ε_ZMW_∕ε_0_. Data marked as N/A could not be quantified due to insufficient signal. The trends visible for Alexa488 are not as clear with JFX650 due to non-specific sticking interactions of the fluorophore with the surface, as evidenced by the increased diffusion time (see Figure 3—**Figure Supplement 3**).

**Figure 3—Figure Supplement 6.**
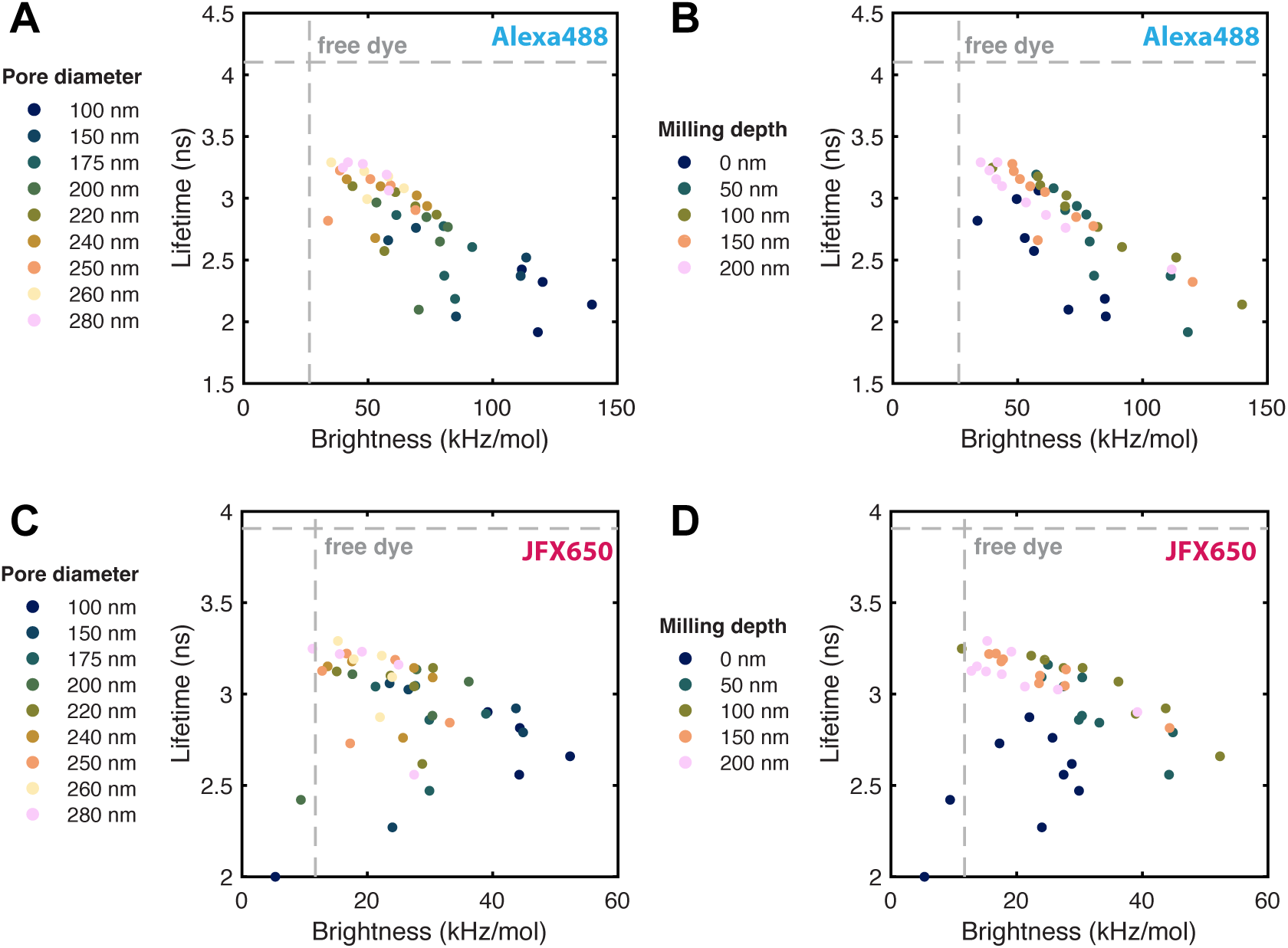
Correlation between fluorescence lifetime and molecular brightness in ZMW for Alexa488 (A,B) and JFX650 (C,D). The data is color coded either by pore diameter (A,C) or milling depth (B,D). The fluorescence lifetime is given by the inverse of the excited state decay rate. A reduction of the lifetime thus indicates the enhancement of the radiative and/or non-radiative relaxation rates due to the ZMW. A negative correlation is observed, where a lower lifetime corresponds with an increased molecular brightness.

**Figure 3—Figure Supplement 7.**
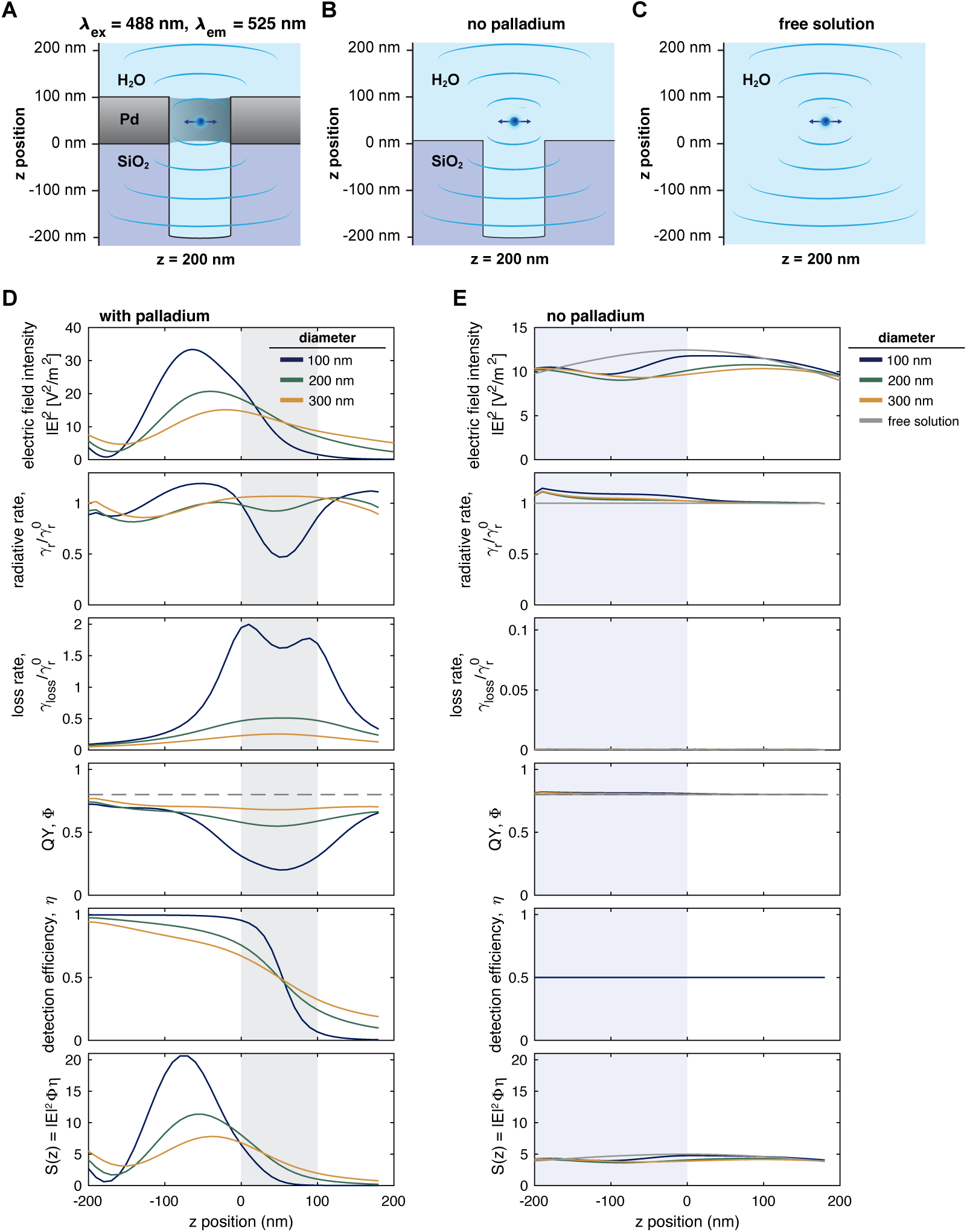
FDTD simulations of excitation field and fluorescence emission in the absence of a Pd layer. **A,B:** Schematic of the simulation setup with (A), without (B) the Pd layer and for the free solution case (C). **D,E:** Computed excitation field intensity E^2^, normalized radiative rate 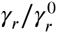, normalized loss rate 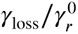 quantum yield Φ, detection efficiency η, and total fluorescence signal S(z) as a function of the z position in the presence (D) and absence (E) of the Pd layer. In D, the position of the metal membrane is indicated as a gray shaded area. In E, the position of the SiO_2_ layer is indicated as a blue shaded area (except for the free diffusion case). The detection efficiency was set to 0.5 in the absence of the Pd layer. A small radiative rate enhancement arises even in the absence of the Pd layer when the dipole is placed in the SiO_2_ nanocavity due to the Purcell effect (***Purcell, 1946***).

**Figure 3—Figure Supplement 8.**
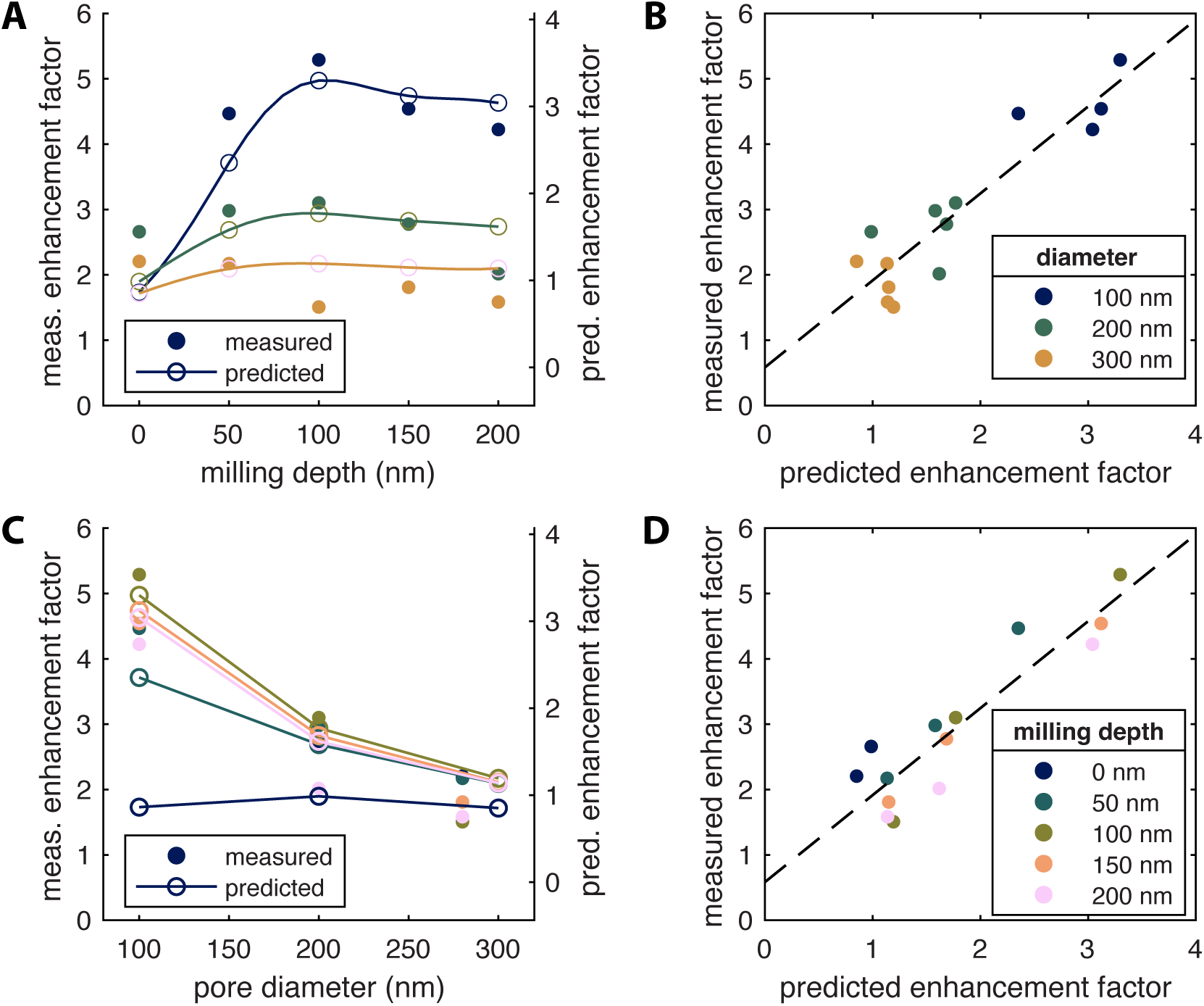
Comparison of experimental and predicted enhancement factors in overmilled Pd ZMWs. **A,C:** Measured and predicted enhancement factors as a function of the milling depths (A) or pore diameter (C). The scaling of the two y-axes was adjusted according to the results of the linear regression between measured and predicted enhancement factors. **B,D:** Plots of the measured versus the predicted enhancement factors, color coded by milling depth (B) or pore diameter (D). The solid line is a linear fit given by y = 1.33x + 0.58. The Pearson correlation coefficient is r = 0.92.

**Figure 4—Figure Supplement 1.**
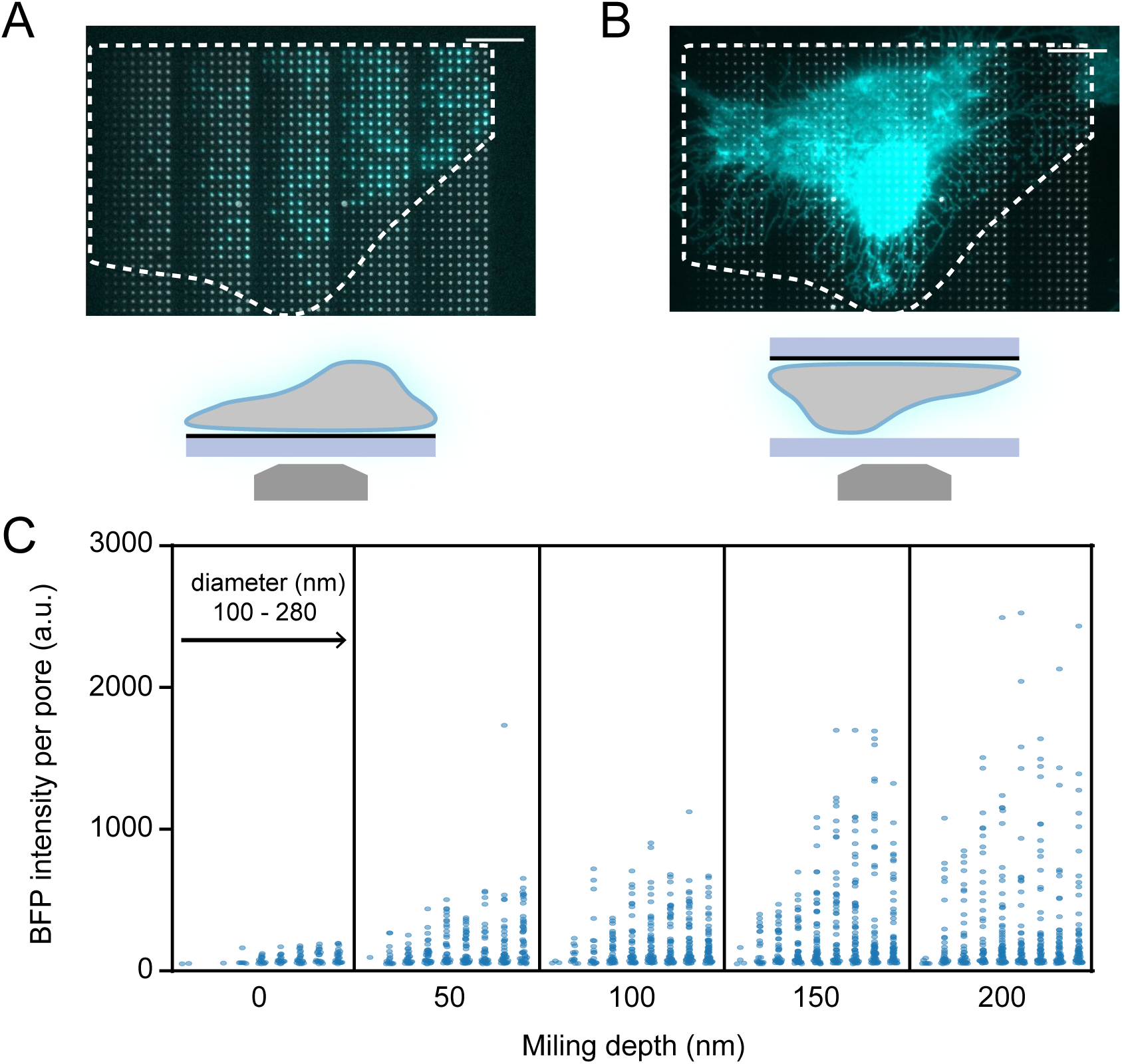
A: Overlay of bright-field and BFP fluorescence images from Figure 4, depicting cells attached to the coverslip. B: Overlay of bright-field and BFP fluorescence images obtained from the same field of view as in (A) for the flipped conformation, where cells attached to the Pd surface are not imaged through the ZMWs but from the open top side. Note that the image is mirrored to visualize cells in the same orientation as in (A). Scale bars: 10 µm. C: BFP fluorescence intensities of individual pores with varying sizes. Each dot represents a single pore. The total number of analyzed pores was 1890.

**Figure 5—Figure Supplement 1.**
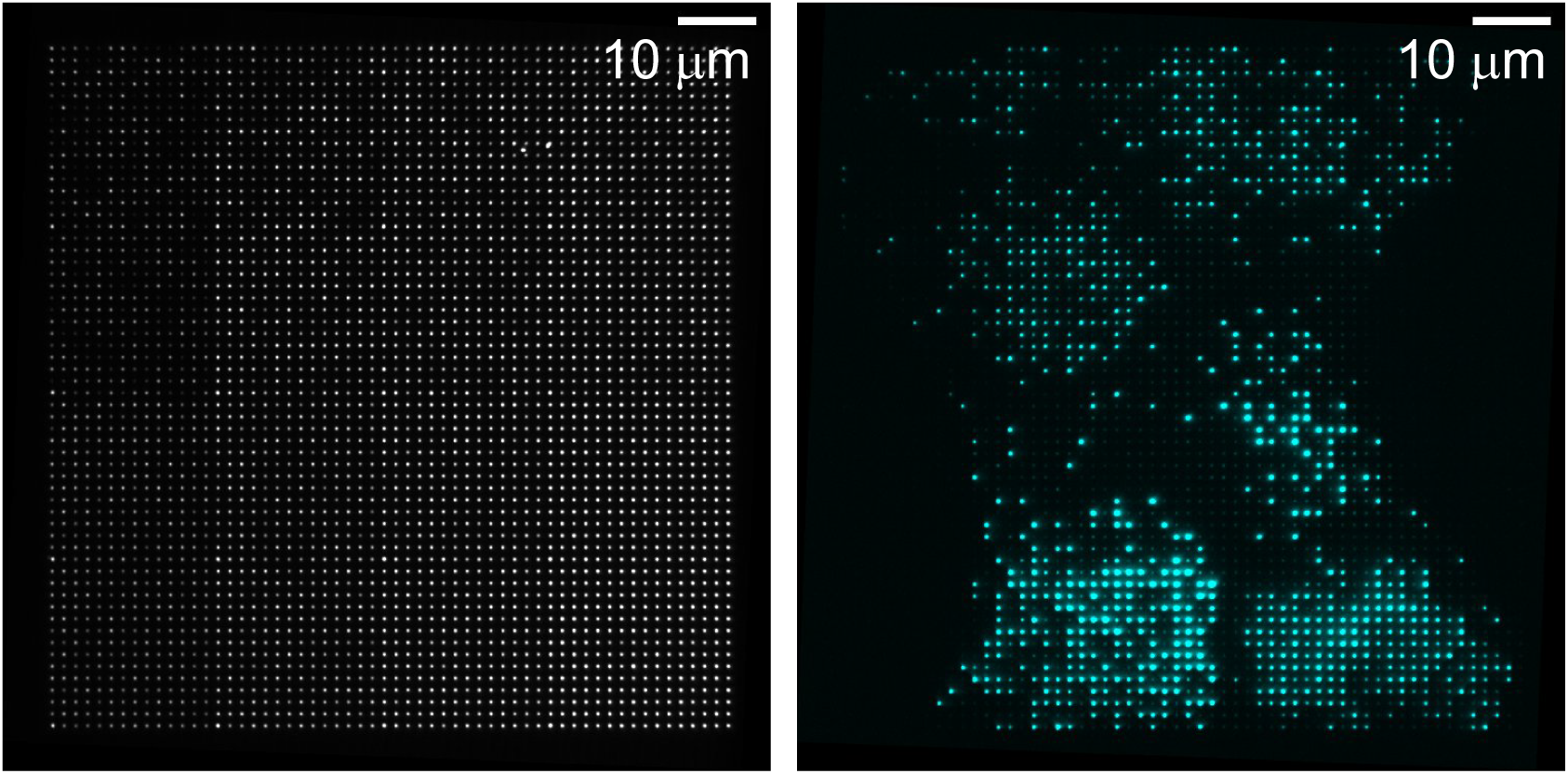
Brightfield and BFP-fluorescence images of array version 2. Orientation is the same as in Figure 1—**Figure Supplement 1** F. The brightfield image (left) shows the location of pores and the BFP-fluorescence image (right) shows their occupation.

**Figure 5—Figure Supplement 2.**
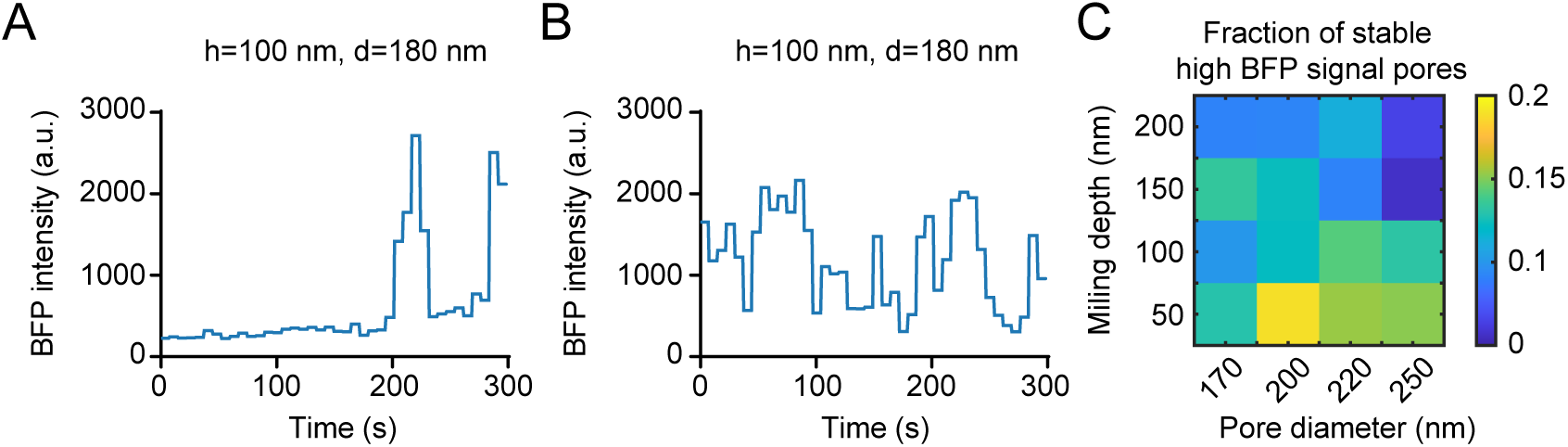
Representative examples of BFP fluorescence intensity switching between high and low signal. **A,B:** Representative examples of BFP fluorescence intensity time traces for individual pores where the intensity switched between the high and low intensity levels. **C:** Percentage of pores with high stable BFP signal for the duration of the movie (300 s). The total number of pores analyzed is 30907, with approximately 1900 pores for each pore size.

**Figure 5—Figure Supplement 3.**
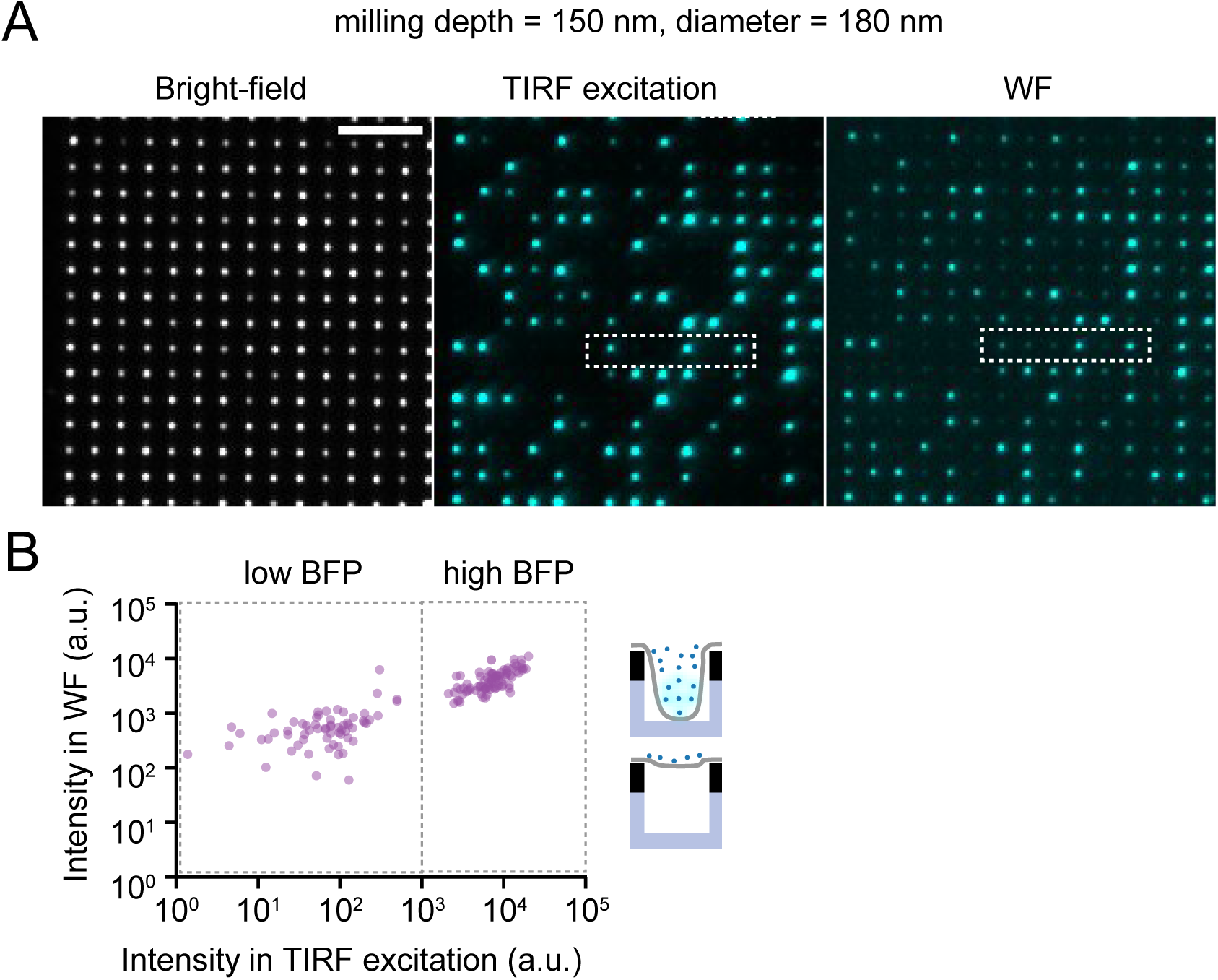
TIRF illumination vs. widefield illumination. **A:** Bright-field image (left), and BFP fluorescence images acquired with either TIRF illumination (center) or WF illumination (right) of nanopores with milling depth ℎ = 150 nm and diameter d = 180 nm. Scale bar, 5 µm. The boxes indicate the location of the zoom-in shown in Figure 5 G. **B:** Scatter plot of BFP intensities in individual pores for TIRF and widefield illumination, with each dot representing a single pore. Note that the two populations can be distinguished much more readily when using TIRF excitation compared to widefield illumination. The number of pores analyzed is 150.

**Figure 6—Figure Supplement 1.**
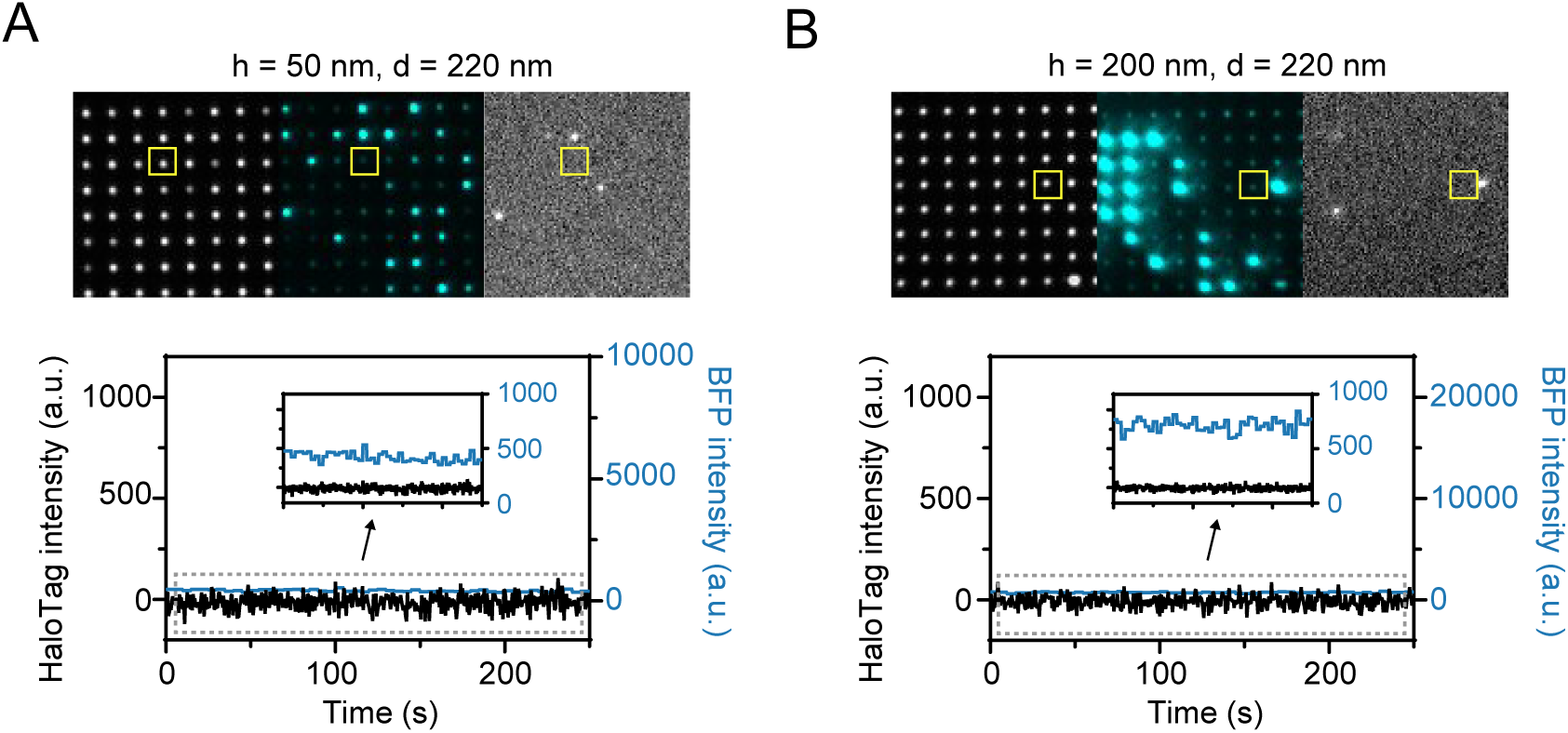
BFP and JFX650-HaloTag signal of pores showing low BFP signal. Bright-field, BFP, and JFX650-HaloTag signal from representative pores showing only low BFP signal with milling depths of 50 nm (left) or 200 nm (right). A fluorescence time trace of a single pore is shown for each condition. Insets provide a zoom-in on the grey box. While a low signal from BFP is observed, no single molecule events of the red fluorophore could be detected. Time interval, 5 s for BFP, 500 ms for JFX650-Halo

## References

Agam G, Gebhardt C, Popara M, Mächtel R, Folz J, Ambrose B, Chamachi N, Chung S, Craggs T, de Boer M, Grohmann D, Ha T, Hartmann A, Hendrix J, Hirschfeld V, Hübner C, Hugel T, Kammerer D, Kang H, Kapanidis A, et al. Reliability and accuracy of single-molecule FRET studies for characterization of structural dynamics and distances in proteins. Nat Methods. 2023;.

Al Masud A, Martin WE, Moonschi FH, Park SM, Srijanto BR, Graham KR, Collier CP, Richards CI. Mixed metal zero-mode guides (ZMWs) for tunable fluorescence enhancement. Nanoscale Adv. 2020; 2:1894–1903. http://dx.doi.org/10.1039/C9NA00641A, doi: 10.1039/C9NA00641A.

Alterovitz SA, Amirtharaj PM, Apell P, Arakawa ET, Ashok J, Barth J, Bezuidenhout DF, Birch JR, Birken HG, Blessing C, Bloomer I, Borghesi A, Callcott TA, Cardona M, Chang Yc, Cotter TM, Edwards DF, Eldridge JE, Fink J, Forouhi AR, et al. Handbook of Optical Constants of Solids, vol. 3. Palik ED, editor, Boston: Academic Press; 1998. https://www.sciencedirect.com/science/article/pii/B9780080556307500031, doi: 10.1016/B978-0-08-055630-7.50003-1.

Aouani H, Wenger J, Gérard D, Rigneault H, Devaux E, Ebbesen TW, Mahdavi F, Xu T, Blair S. Crucial Role of the Adhesion Layer on the Plasmonic Fluorescence Enhancement. ACS Nano. 2009; 3(7):2043–2048. https://doi.org/10.1021/nn900460t, doi: 10.1021/nn900460t, pMID: 19518085.

Assad ON, Gilboa T, Spitzberg J, Juhasz M, Weinhold E, Meller A. Light-Enhancing Plasmonic-Nanopore Biosensor for Superior Single-Molecule Detection. Advanced Materials. 2016; 29(9):1605442. https://onlinelibrary.wiley.com/doi/abs/10.1002/adma.201605442, doi: 10.1002/adma.201605442.

Auger T, Mathé J, Viasnoff V, Charron G, Di Meglio JM, Auvray L, Montel F. Zero-Mode Waveguide Detection of Flow-Driven DNA Translocation through Nanopores. Phys Rev Lett. 2014 7; 113:028302. https://link.aps.org/doi/10.1103/PhysRevLett.113.028302, doi: 10.1103/PhysRevLett.113.028302.

Baibakov M, Patra S, Claude JB, Moreau A, Lumeau J, Wenger J. Extending Single Molecule Förster Resonance Energy Transfer (FRET) Range Beyond 10 Nanometers in Zero-Mode Waveguides. ACS Nano. 2019; 0(ja):null. https://doi.org/10.1021/acsnano.9b04378, doi: 10.1021/acsnano.9b04378, pMID: 31283186.

Bard JAM, Goodall EA, Greene ER, Jonsson E, Dong KC, Martin A. Structure and Function of the 26S Proteasome. Annual Review of Biochemistry. 2018; 87(1):697–724. https://doi.org/10.1146/annurev-biochem-062917-011931, doi: 10.1146/annurev-biochem-062917-011931, pMID: 29652515.

Bharadwaj P, Novotny L. Spectral dependence of single molecule fluorescence enhancement. Optics Express. 2007; 15(21):14266–14274. doi: 10.1364/oe.15.014266.

Boyd GT, Yu ZH, Shen YR. Photoinduced luminescence from the noble metals and its enhancement on roughened surfaces. Phys Rev B. 1986 6; 33:7923–7936. https://link.aps.org/doi/10.1103/PhysRevB.33.7923, doi: 10.1103/PhysRevB.33.7923.

Bronson JE, Fei J, Hofman JM, Gonzalez RL, Wiggins CH. Learning Rates and States from Biophysical Time Series: A Bayesian Approach to Model Selection and Single-Molecule FRET Data. Biophysical Journal. 2009; 97(12):3196–3205. https://www.sciencedirect.com/science/article/pii/S0006349509015136, doi: 10.1016/j.bpj.2009.09.031.

García AJ, Boettiger D. Integrin–fibronectin interactions at the cell-material interface: initial integrin binding and signaling. Biomaterials. 1992; 20(23):2427–2433. https://www.sciencedirect.com/science/article/pii/S0142961299001702, doi: 10.1016/S0142-9612(99)00170-2.

Gérard D, Wenger J, Bonod N, Popov E, Rigneault H, Mahdavi F, Blair S, Dintinger J, Ebbesen TW. Nanoapertureenhanced fluorescence: Towards higher detection rates with plasmonic metals. Physical Review B. 2008 1; 77:045413. https://link.aps.org/doi/10.1103/PhysRevB.77.045413, doi: 10.1103/PhysRevB.77.045413.

Gregor I, Chizhik A, Karedla N, Enderlein J. Metal-induced energy transfer. Nanophotonics. 2019; 8(10):1689– 1699. doi: 10.1515/nanoph-2019-0201.

Grimm JB, Xie L, Casler JC, Patel R, Tkachuk AN, Falco N, Choi H, Lippincott-Schwartz J, Brown TA, Glick BS, Liu Z, Lavis LD. A General Method to Improve Fluorophores Using Deuterated Auxochromes. JACS Au. 2021; 1(5):690–696. https://doi.org/10.1021/jacsau.1c00006, doi: 10.1021/jacsau.1c00006.

Hellenkamp B, Schmid S, Doroshenko O, Opanasyuk O, Kühnemuth R, Rezaei Adariani S, Ambrose B, Aznauryan M, Barth A, Birkedal V, Bowen ME, Chen H, Cordes T, Eilert T, Fijen C, Gebhardt C, Götz M, Gouridis G, Gratton E, Ha T, et al. Precision and accuracy of single-molecule FRET measurements—a multi-laboratory benchmark study. Nature Methods. 2018-09; 15(9):669–676. https://www.nature.com/articles/s41592-018-0085-0, doi: 10.1038/s41592-018-0085-0.

Hinterdorfer P, Oijen A, editors. Handbook of Single-Molecule Biophysics. 1 ed. Springer New York, NY; 2009. doi: 10.1007/978-0-387-76497-9.

Holzmeister P, Pibiri E, Schmied JJ, Sen T, Acuna GP, Tinnefeld P. Quantum yield and excitation rate of single molecules close to metallic nanostructures. Nature Communications. 2014; 5(1):5356. https://www.nature.com/articles/ncomms6356, doi: 10.1038/ncomms6356.

Hoyer M, Crevenna AH, Correia JRC, Quezada AG, Lamb DC. Zero-mode waveguides visualize the first steps during gelsolin-mediated actin filament formation. Biophys J. 2022; 121(2):327–335. doi: 10.1016/j.bpj.2021.12.011.

Jackson JD. Classical electrodynamics. New York: Wiley; 1962.

Jiang X, Bruzewicz DA, Thant MM, Whitesides GM. Palladium as a Substrate for Self-Assembled Monolayers Used in Biotechnology. Analytical Chemistry. 2004; 76(20):6116–6121. https://doi.org/10.1021/ac049152t, doi: 10.1021/ac049152t, pMID: 15481961.

Jiao X, Peterson EM, Harris JM, Blair S. UV Fluorescence Lifetime Modification by Aluminum Nanoapertures. ACS Photonics. 2014; 1(12):1270–1277. doi: 10.1021/ph500267n.

Kaminski F, Sandoghdar V, Agio M. Finite-Difference Time-Domain Modeling of Decay Rates in the Near Field of Metal Nanostructures. Journal of Computational and Theoretical Nanoscience. 2007; 4(3):635–643.

Klughammer N, Barth A, Dekker M, Fragasso A, Onck P, Dekker C. Diameter Dependence of Transport through Nuclear Pore Complex Mimics Studied Using Optical Nanopores. bioRxiv. 2023; https://www.biorxiv.org/content/early/2023/02/18/2023.02.18.529008, doi: 10.1101/2023.02.18.529008.

Klughammer N, Dekker C. Palladium zero-mode waveguides for optical single-molecule detection with nanopores. Nanotechnology. 2021 feb; 32(18):18LT01. https://doi.org/10.1088/1361-6528/abd976, doi: 10.1088/1361-6528/abd976.

Kubitscheck U. Fluorescence Microscopy: From Principles to Biological Applications. 2nd ed ed. John Wiley & Sons, Incorporated; 2017. https://public.ebookcentral.proquest.com/choice/publicfullrecord.aspx?p=4834057.

Larkin J, Henley RY, Jadhav V, Korlach J, Wanunu M. Length-independent DNA packing into nanopore zero-mode waveguides for low-input DNA sequencing. Nature nanotechnology. 2017; 12(12):1169. doi: 10.1038/nnano.2017.176.

Lenne PF, Rigneault H, Marguet D, Wenger J. Fluorescence fluctuations analysis in nanoapertures: physical concepts and biological applications. Histochemistry and Cell Biology. 2008 9; 130(5):795. https://doi.org/10.1007/s00418-008-0507-7, doi: 10.1007/s00418-008-0507-7.

Lerner E, Barth A, Hendrix J, Ambrose B, Birkedal V, Blanchard S, Börner R, Sung Chung H, Cordes T, Craggs T, Deniz A, Diao J, Fei J, Gonzalez R, Gopich I, Ha T, Hanke C, Haran G, Hatzakis N, Hohng S, et al. FRET-based dynamic structural biology: Challenges, perspectives and an appeal for open-science practices. eLife. 2021; 10:e60416.

Levene MJ, Korlach J, Turner SW, Foquet M, Craighead HG, Webb WW. Zero-mode waveguides for singlemolecule analysis at high concentrations. science. 2003; 299(5607):682–686. doi: 10.1126/science.1079700.

Liu YJ, Le Berre M, Lautenschlaeger F, Maiuri P, Callan-Jones A, Heuzé M, Takaki T, Voituriez R, Piel M. Confinement and Low Adhesion Induce Fast Amoeboid Migration of Slow Mesenchymal Cells. Cell. 2015; 160(4):659– 672. https://www.sciencedirect.com/science/article/pii/S0092867415000082, doi: 10.1016/j.cell.2015.01.007.

Liu Z, Lavis L, Betzig E. Imaging Live-Cell Dynamics and Structure at the Single-Molecule Level. Mol Cell. 2015; 58(4):644–659. doi: 10.1016/j.molcel.2015.02.033.

Los GV, Encell LP, McDougall MG, Hartzell DD, Karassina N, Zimprich C, Wood MG, Learish R, Ohana RF, Urh M, Simpson D, Mendez J, Zimmerman K, Otto P, Vidugiris G, Zhu J, Darzins A, Klaubert DH, Bulleit RF, Wood KV. HaloTag: A Novel Protein Labeling Technology for Cell Imaging and Protein Analysis. ACS Chemical Biology. 2008; 3(6):373–382. https://doi.org/10.1021/cb800025k, doi: 10.1021/cb800025k.

Love JC, Estroff LA, Kriebel JK, Nuzzo RG, Whitesides GM. Self-Assembled Monolayers of Thiolates on Metals as a Form of Nanotechnology. Chemical Reviews. 2005; 105(4):1103–1170. https://doi.org/10.1021/cr0300789, doi: 10.1021/cr0300789, pMID: 15826011.

Love JC, Wolfe DB, Haasch R, Chabinyc ML, Paul KE, Whitesides GM, Nuzzo RG. Formation and Structure of Self-Assembled Monolayers of Alkanethiolates on Palladium. Journal of the American Chemical Society. 2003; 125(9):2597–2609. https://doi.org/10.1021/ja028692+, doi: 10.1021/ja028692+, pMID: 12603148.

Malekian B, Schoch RL, Robson T, Ferrand Drake del Castillo G, Xiong K, Emilsson G, Kapinos LE, Lim RYH, Dahlin A. Detecting Selective Protein Binding Inside Plasmonic Nanopores: Toward a Mimic of the Nuclear Pore Complex. Frontiers in Chemistry. 2018; 6:637. https://www.frontiersin.org/article/10.3389/fchem.2018.00637, doi: 10.3389/fchem.2018.00637.

Martin WE, Srijanto BR, Collier CP, Vosch T, Richards CI. A Comparison of Single-Molecule Emission in Aluminum and Gold Zero-Mode Waveguides. The Journal of Physical Chemistry A. 2016; 120(34):6719–6727. https://doi.org/10.1021/acs.jpca.6b03309, doi: 10.1021/acs.jpca.6b03309, pMID: 27499174.

Milo R, Phillips R. Cell Biology by the Numbers. Garland Science, Taylor & Francis Group; 2015. http://book.bionumbers.org/, doi: 10.1201/9780429258770.

Miyake T, Tanii T, Sonobe H, Akahori R, Shimamoto N, Ueno T, Funatsu T, Ohdomari I. Real-Time Imaging of Single-Molecule Fluorescence with a Zero-Mode Waveguide for the Analysis of Protein-Protein Interaction. Analytical Chemistry. 2008; 80(15):6018–6022. https://doi.org/10.1021/ac800726g, doi: 10.1021/ac800726g, pMID: 18563914.

Mooradian A. Photoluminescence of Metals. Physical Review Letters. 1969; 22(5):185–187. https://link.aps.org/doi/10.1103/PhysRevLett.22.185, doi: 10.1103/PhysRevLett.22.185.

Moran-Mirabal JM, Torres AJ, Samiee KT, Baird BA, Craighead HG. Cell investigation of nanostructures: zero-mode waveguides for plasma membrane studies with single molecule resolution. Nanotechnology. 2007 apr; 18(19):195101. https://doi.org/10.1088%2F0957-4484%2F18%2F19%2F195101, doi: 10.1088/0957-4484/18/19/195101.

Müller BK, Zaychikov E, Bräuchle C, Lamb DC. Pulsed Interleaved Excitation. Biophys J. 2005; 89(5):3508–3522. doi: 10.1529/biophysj.105.064766.

Novotny L, Hecht B. Principles of Nano-Optics. Cambridge University Press; 2006. doi: 10.1017/CBO9780511813535.

Patra S, Claude JB, Wenger J. Fluorescence Brightness, Photostability, and Energy Transfer Enhancement of Immobilized Single Molecules in Zero-Mode Waveguide Nanoapertures. ACS Photonics. 2022; 9(6):2109– 2118. https://doi.org/10.1021/acsphotonics.2c00349, doi: 10.1021/acsphotonics.2c00349.

Purcell EM. Spontaneous Emission Probabilities at Radio Frequencies. Phys Rev Lett. 1946; 69(681):839–839. doi: 10.1007/978-1-4615-1963-8_40.

Rhoads A, Au KF. PacBio Sequencing and Its Applications. Genomics, Proteomics & Bioinformatics. 2015; 13(5):278–289. https://www.sciencedirect.com/science/article/pii/S1672022915001345, doi: 10.1016/j.gpb.2015.08.002.

Richards CI, Luong K, Srinivasan R, Turner SW, Dougherty DA, Korlach J, Lester HA. Live-Cell Imaging of Single Receptor Composition Using Zero-Mode Waveguide Nanostructures. Nano Letters. 2012; 12(7):3690–3694. https://doi.org/10.1021/nl301480h, doi: 10.1021/nl301480h, pMID: 22668081.

Rigneault H, Capoulade J, Dintinger J, Wenger J, Bonod N, Popov E, Ebbesen TW, Lenne PF. Enhancement of Single-Molecule Fluorescence Detection in Subwavelength Apertures. Phys Rev Lett. 2005 9; 95:117401. https://link.aps.org/doi/10.1103/PhysRevLett.95.117401, doi: 10.1103/PhysRevLett.95.117401.

Samiee KT, Foquet M, Guo L, Cox EC, Craighead HG. λ-Repressor Oligomerization Kinetics at High Concentrations Using Fluorescence Correlation Spectroscopy in Zero-Mode Waveguides. Biophysical Journal. 2005; 88(3):2145 – 2153. http://www.sciencedirect.com/science/article/pii/S0006349505732768, doi: https://doi.org/10.1529/biophysj.104.052795.

Samiee KT, Moran-Mirabal JM, Cheung YK, Craighead HG. Zero Mode Waveguides for Single-Molecule Spectroscopy on Lipid Membranes. Biophysical Journal. 2006; 90(9):3288–3299. https://linkinghub.elsevier.com/retrieve/pii/S0006349506725115, doi: 10.1529/biophysj.105.072819.

Schrimpf W, Barth A, Hendrix J, Lamb DC. PAM: A Framework for Integrated Analysis of Imaging, Single-Molecule, and Ensemble Fluorescence Data. Biophysical Journal. 2018; 114(7):1518–1528. https://www.sciencedirect.com/science/article/pii/S0006349518302959, doi: https://doi.org/10.1016/j.bpj.2018.02.035.

Tanii T, Akahori R, Higano S, Okubo K, Yamamoto H, Ueno T, Funatsu T. Improving zero-mode waveguide structure for enhancing signal-to-noise ratio of real-time single-molecule fluorescence imaging: A computational study. Phys Rev E Stat Nonlin Soft Matter Phys. 2013; 88(1):012727. doi: 10.1103/physreve.88.012727.

VandeVondele S, Vörös J, Hubbell JA. RGD-grafted poly-l-lysine-graft-(polyethylene glycol) copolymers block non-specific protein adsorption while promoting cell adhesion. Biotechnology and Bioengineering. 2003; 82(7):784–790. https://onlinelibrary.wiley.com/doi/abs/10.1002/bit.10625, doi: 10.1002/bit.10625.

Wenger J, Conchonaud F, Dintinger J, Wawrezinieck L, Ebbesen TW, Rigneault H, Marguet D, Lenne PF. Diffusion Analysis within Single Nanometric Apertures Reveals the Ultrafine Cell Membrane Organization. Biophysical Journal. 2007-02; 92(3):913–919. https://www.sciencedirect.com/science/article/pii/S0006349507708986, doi: 10.1529/biophysj.106.096586.

Wu M, Liu W, Hu J, Zhong Z, Rujiralai T, Zhou L, Cai X, Ma J. Fluorescence enhancement in an over-etched gold zero-mode waveguide. Optics express. 2019; 27 13:19002–19018. doi: https://doi.org/10.1364/OE.27.019002.

Xu C, Webb WW. 11. In: Lakowicz JR, editor. Multiphoton Excitation of Molecular Fluorophores and Nonlinear Laser Microscopy Boston, MA: Springer US; 2002. p. 471–540. https://doi.org/10.1007/0-306-47070-5_11, doi: 10.1007/0-306-47070-5_11.

Yan X, Hoek TA, Vale RD, Tanenbaum ME. Dynamics of Translation of Single mRNA Molecules In Vivo. Cell. 2016; 165(4):976–989. https://www.sciencedirect.com/science/article/pii/S0092867416304779, doi: 10.1016/j.cell.2016.04.034.

Yang S, Klughammer N, Barth A, Datasets underlying the paper Zero-mode waveguide nanowells for singlemolecule detection in living cells. Zenodo; 2023. https://doi.org/10.5281/zenodo.8060099, doi: 10.5281/zen-odo.8060099.

